# Integrated platform for multi-scale molecular imaging and phenotyping of the human brain

**DOI:** 10.1101/2022.03.13.484171

**Authors:** Juhyuk Park, Ji Wang, Webster Guan, Lars A. Gjesteby, Dylan Pollack, Lee Kamentsky, Nicholas B. Evans, Jeff Stirman, Xinyi Gu, Chuanxi Zhao, Slayton Marx, Minyoung E. Kim, Seo Woo Choi, Michael Snyder, David Chavez, Clover Su-Arcaro, Yuxuan Tian, Chang Sin Park, Qiangge Zhang, Dae Hee Yun, Mira Moukheiber, Guoping Feng, X. William Yang, C. Dirk Keene, Patrick R. Hof, Satrajit S. Ghosh, Matthew P. Frosch, Laura J. Brattain, Kwanghun Chung

## Abstract

Understanding cellular architectures and their connectivity is essential for interrogating system function and dysfunction. However, we lack technologies for mapping the multi-scale details of individual cells in the human organ-scale system. To address this challenge, we developed a platform that simultaneously extracts spatial, molecular, morphological, and connectivity information of individual cells from the same human brain, by integrating novel chemical, mechanical, and computational tools. The platform includes three key tools: (i) a vibrating microtome for ultra-precision slicing of large-scale tissues without losing cellular connectivity (MEGAtome), (ii) a polymer hydrogel-based tissue processing technology for multiplexed multiscale imaging of human organ-scale tissues (mELAST), and (iii) a computational pipeline for reconstructing 3D connectivity across multiple brain slabs (UNSLICE). We demonstrated the transformative potential of our platform by analyzing human Alzheimer’s disease pathology at multiple scales and demonstrating scalable neural connectivity mapping in the human brain.

**One-Sentence Summary:** We developed an integrated, scalable platform for highly multiplexed, multi-scale phenotyping and connectivity mapping in the same human brain tissue, which incorporated novel tissue processing, labeling, imaging, and computational technologies.

## Main Text

A detailed mapping of the anatomical and molecular architectures of brain cells and their brain-wide connectivity is essential for understanding human brain function, the etiology of brain injuries, and the impact of brain diseases (*1–3*). Neuroimaging techniques (e.g., MRI, fMRI, DWI) have vastly increased our knowledge of the functional and structural organization of the human brain (*4–8*), but these methods fail to capture the fine structural, cellular, and molecular details due to their limitations in spatial resolution (*9*, *10*). Advanced histological approaches have provided a new window into local organization of cells but fail to capture brain-wide 3D spatial information. Indeed, most histological studies resort to a subsampling approach, where serial sections are stained for different markers. Co-expression analyses rely on indirect inferences between sections, limited by the host of the antibodies and the number of channels available for imaging.

With the development of high-throughput single-cell/single-nucleus RNA-seq technology (*3*, *11*), many more cell types can now be distinguished by their transcriptomic profiles, but these methods inherently limit the acquisition of spatial information and anatomical details of cells. Advances in spatial transcriptomics have enabled systematic analysis of cellular composition within tissue contexts (*12*). Although these tools have dramatically advanced our understanding of the local cellular landscape in the brain, scaling these approaches to encompass entire human brains across many individuals remains challenging. Moreover, transcriptomic analysis alone cannot provide key information critical to understanding functional properties of cells, such as cellular morphology, connectivity, subcellular architectures, and protein posttranslational modifications and tracking.

Recently, significant progress has been made in integrating information across modalities to produce more comprehensive brain atlases. In a 2016 study (*13*), the Allen Brain Institute for Brain Science mapped the entire human brain using 50 µm-thick slices by alternating staining for Nissl, non-phosphorylated neurofilament protein, and parvalbumin, and combined this information with anatomical MR images of the same brain. This brain atlas was the first to integrate neuroimaging data with cyto- and chemoarchitectural data, thereby resulting in more thorough parcellations of brain regions and cells, as well as a comprehensive quantification of the complex interplay of microstructural features in the human brain. However, the limited resolution and number of features that can be recovered using these approaches cannot capture the multi-scale and high-dimensional complexity of the human brain. Therefore, there is a need to develop technologies that allow us to capture multi-scale multi-omic properties of individual cells and their brain-wide connectivity.

To address this challenge, we developed a fully integrated scalable technology platform to establish a three-dimensional (3D) human brain cell atlas at subcellular resolution by simultaneously mapping brain-wide structures and high-dimensional features (e.g., spatial, molecular, morphological, microenvironment, nanoscopic, and connectivity information) of cells acquired from the same whole human brains (**Fig. 1**). We accomplished this by seamlessly integrating new chemical, mechanical, and computational tools, enabling highly multiplexed multi-scale 3D proteomic phenotyping of human brain tissues. We demonstrated its utility and scalability by processing the whole human brain hemispheres and investigating Alzheimer’s disease (AD) pathology.

**Fig. 1.**
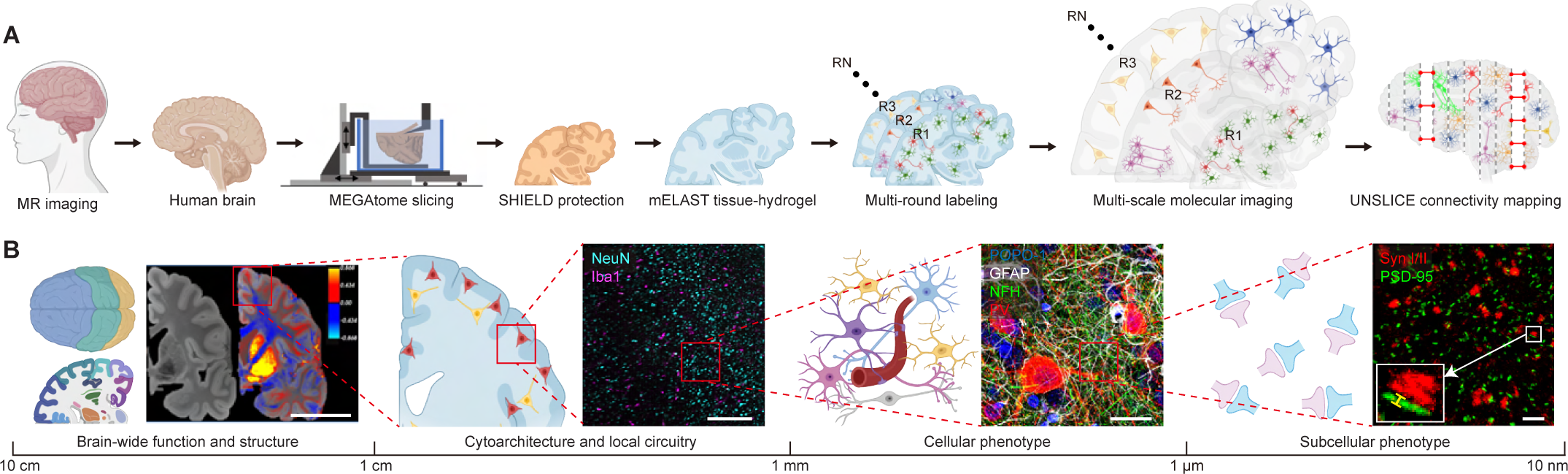
Integrated platform for multi-scale molecular interrogation of the human brain. (**A**) The workflow of the platform includes slicing, processing, labeling, imaging, and computational stitching of the human brain tissues. After MR imaging, human brain hemispheres are sliced into millimeter-thick slabs using MEGAtome. The slabs are SHIELD-processed and transformed into mELAST tissue-hydrogels. The tough, elastic and size-adjustable mELAST tissue-hydrogels undergo multi-round labeling and imaging at multiple scales to extract high-dimensional features. The images acquired from the sliced human brain tissues are computationally reintegrated using the UNSLICE pipeline for multi-level connectivity mapping. (**B**) 3D multi-scale profiling of the human brain captures brain-wide structures, cytoarchitecture and local circuitry, cellular and subcellular features from the same tissue. Scale bars, 5 cm, 300 μm, 25 μm, and 500 nm (inset, 25 nm), from left to right.

### MEGAtome enables high-precision slicing of ultra-large biological tissues

Though state-of-the-art tissue clearing and processing methods (*14–18*) can render intact tissues optically transparent, the working distance (WD) for the objectives of existing microscope systems cannot cover the entire volume of human and large animal organs. In addition, residual variations in refractive index within cleared tissues cause loss of resolution, therefore mechanical sectioning is still required for imaging of large tissues.

Vibratomes are tools for sectioning soft tissues (*19*, *20*). Conventional vibratomes, however, are limited to small samples and often cause tissue damage. For example, Leica VT1200 (one of the most used commercial vibratomes) has a maximum sample size of 33 mm [width] × 20 mm [height] × 50 mm [length], unsuitable for processing human organs. Furthermore, due to their limited blade vibrating speed and large out-of-plane blade vibration, cut surfaces often suffer from abrasion, tears, and deformation. Commercial vibratomes are also not optimized to uniformly slice highly heterogeneous tissues, such as human brains, which exhibit greater anatomical complexity and heterogeneity in their mechanical properties. Failure to slice with precision causes loss of valuable information required for in silico reconstruction of organs, of particular relevance for human brain studies where connectivity is critically associated to functionality.

To address these limitations, we developed a highly versatile vibratome, termed MEGAtome (**M**echanically **E**nhanced **G**reat-size **A**brasion-free vibra**tome**), that enables ultra-precision slicing of a wide range of biological samples, from small organoids (∼0.5 mm in diameter) to intact human brain hemispheres (approximately 65 mm [width] × 180 mm [height] × 130 mm [length]) and large arrays of animal organs (**Fig. 2**). MEGAtome minimizes tissue damage and maximally preserves intra- and inter-slice information by enabling high-frequency blade vibration with increased amplitude and low out-of-plane vibration displacement (**fig. S1**).

**Fig. 2.**
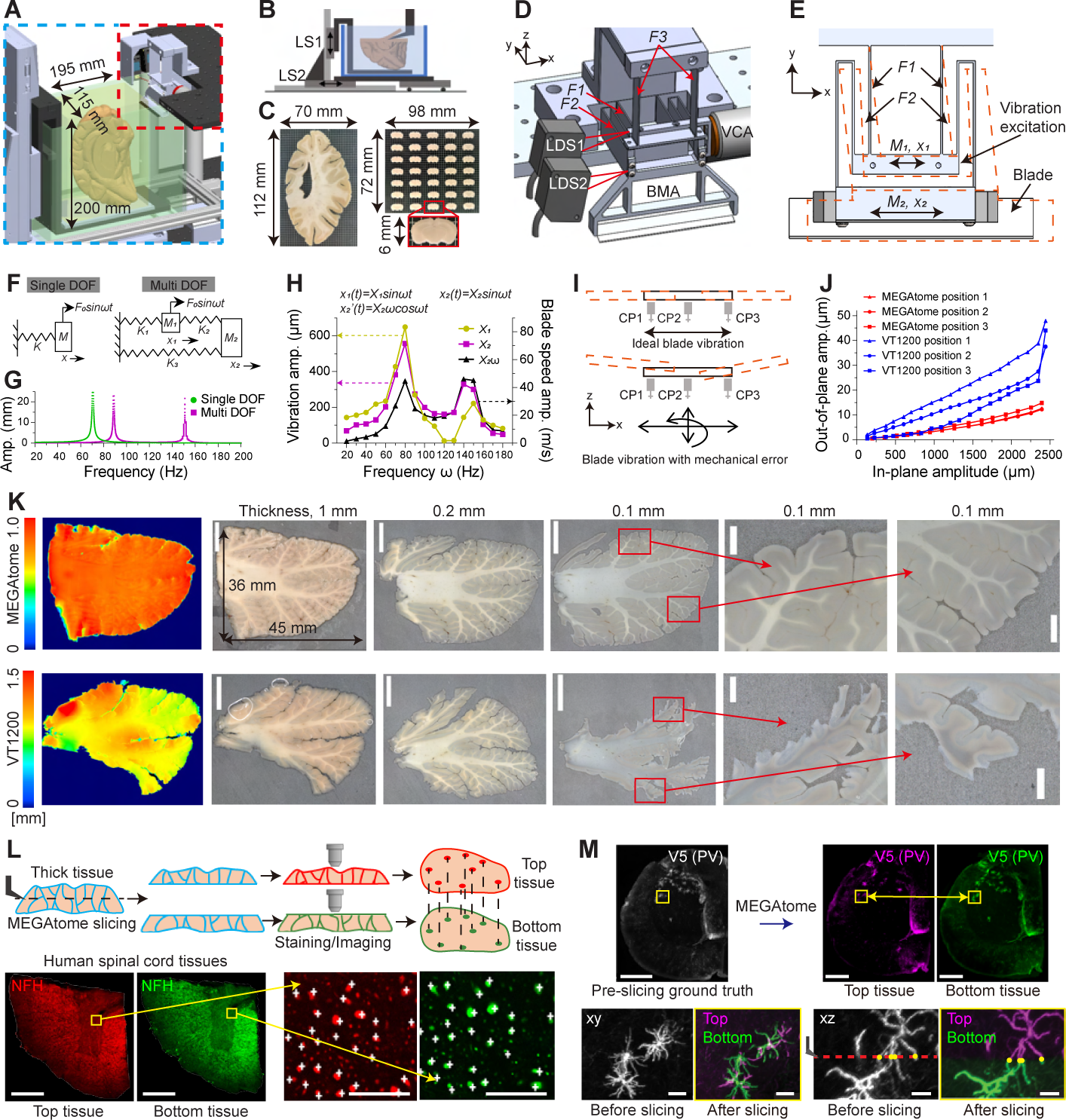
Mechanical design and characterization of MEGAtome for scalable, information lossless slicing. (**A**) MEGAtome consists of two main modules: a blade vibration generation and control platform (in red frame), and a sample mounting and feeding platform (in blue frame). (**B**) The sample mounting and feeding platform is configured through two linear stages (LS). (**C**) MEGAtome-sliced whole human brain hemispheres and mouse brain arrays. (**D-E**) Mechanical design of the blade vibration generation and control mechanism shown in a 3D view (D), and in xy plane (E). (**F**) A single degree of freedom (DOF) model (left, common commercial vibratomes) and a multi-DOF vibration model (right, MEGAtome), in which *K_1_*, *K_2_*, and *K_3_* represents the stiffnesses of *F_1_*, *F_2_*, and *F_3_*, respectively. (**G**) Simulation results showing the forced vibration response of a single DOF system and a multi DOF system. (**H**) Measurements of *x_1_* and *x_2_* indicate that the first and second resonant modes of the mechanism occur near 80 and 140 Hz, respectively. (**I**) Demonstration of ideal blade vibration and real blade vibration accompanied by parasitic errors. (**J**) Measurements of blade out-of-plane parasitic vibration amplitude versus in-plane vibration amplitude of VT1200 vibratome and MEGAtome. (**K**) MEGAtome generated 1.0, 0.2, and 0.1 mm-thick slices from whole human cerebellums (top row). The 1 mm-thick slices were scanned by an optical profiler showing the surface height variance. Scale bars, 10 mm (whole slice figures) and 2 mm (enlarged figures). (**L**) MEGAtome does not cause a loss of connectivity information of NFH+ axons in the cut surfaces of human brain spinal cord tissues after slicing. Scale bars, 1 mm (left figures) and 40 μm (right enlarged figures). (**M**) MEGAtome preserves neuron micromorphology and connectivity in PV-Cre MORF mouse brain tissues while slicing. Scale bars, 500 μm (top row) and 50 μm (bottom row). The images were obtained with the 20x/0.5NA (L) and 4x/0.2NA objectives (M).

Analytical modeling and experimental data have shown that three elements benefit the precision and scalability of vibratome sectioning: (i) high blade vibration frequency, (ii) high vibration amplitude, and (iii) minimal out-of-plane blade displacement (*21–23*). To achieve these crucial elements, we designed a new blade vibration generation and control mechanism based on a multi-degree-of-freedom (multi-DOF) flexure series (**Fig. 2, A to E**). The blade vibrating frequency and amplitude are controlled by the frequency and amplitude of the driving current.

The advantage of multi-DOF over single-DOF (in most commercial vibratomes) lies in its capability to operate at higher frequencies and increased amplitudes (**Fig. 2F**). Simulations and experiments (*21*, *24*) reveal two resonant frequencies for our vibration system, approximately at 80 and 145 Hz (**Fig. 2, G and H**). Comparison of the input and output vibration amplitude at their optimal frequencies (**fig. S2**) reveal MEGAtome’s linear increase in blade vibration amplitude with input, reaching 3.4 mm peak-to-peak at 140 Hz, while VT1200 performs best at 2.4 mm peak-to-peak at 85 Hz. These findings demonstrate that MEGAtome overcomes the trade-off between blade vibrating frequency and amplitude, achieving a 2.33 times higher blade vibrating speed than VT1200 at 1.65 times higher frequency.

To minimize the out-of-plane parasitic blade vibrations, we increased the system’s out-of-plane stiffness through the design of flexure series (**Fig. 2, D, E, and I**). The configuration of the flexures, particularly the vertical orientation of *F_3_*, ensures high stiffness in out-of-plane directions. To experimentally characterize this, we used high-precision capacitance probes (CP) to measure the out-of-plane displacement of the same blade mounted on both machines. The data demonstrated that the MEGAtome exhibits lower out-of-plane vibration amplitudes compared to VT1200, even though MEGAtome operates at much higher frequency (**Fig. 2J**).

It is worth noting that MEGAtome is not suitable for sectioning hard samples, such as bone, paraffin-embedded, or frozen samples due to their high sample resistance, which disrupts the blade’s vibration accuracy. However, MEGAtome excels in sectioning (i) soft tissues (e.g., fresh or chemically fixed tissues) (ii) dense and highly myelinated tissues (e.g., human brainstem and spinal cord), and (iii) heterogeneous samples (e.g., human organs with connective tissues and embedded blood vessels) (**figs. S1 and S3**). Next, we compared the precision of MEGAtome and VT1200 by slicing large human cerebellum tissues (36 x 45 x 20 mm) from the same brain into 1.0, 0.2, and 0.1 mm-thick sections and measuring their surface profiles. Visual inspection and evaluation of optical profiles of surface height variance images showed that MEGAtome produces intact slices with excellent surface evenness, preserving fine tissue architectures (**Fig. 2K**). VT1200 not only generated highly uneven tissue sections, but also caused severe damage to the cerebellar gyri and sulcus structures.

To confirm that MEGAtome achieves information-lossless tissue slicing, the distribution pattern of NFH+ neuronal fibers from adjacent sections was compared (**Fig. 2L**). Single fiber resolution matching at cross sections indicates that MEGAtome slicing caused no damage or loss of fiber connectivity information. Additionally, we used a mononucleotide repeat frameshift (MORF) mouse brain expressing V5 peptide (*25*, *26*) to collect images before and after slicing (**Fig. 2M**). Fiber connectivity and morphology of the neurons remained intact after slicing, highlighting MEGAtome as an ideal solution for slicing tissues while preserving their multi-scale integrity and cellular connectivity information.

### Whole-mount slicing and imaging of large-scale tissue samples

Mechanical slicing into millimeter-thick slabs is necessary to image large samples, such as human organs. Minimizing the number of dissected tissue blocks from large samples is advantageous as it can drastically reduce human labor, cost, and information-loss. For instance, whole-mount sectioning of an entire human brain hemisphere into intact thick coronal slabs is highly desirable as slicing the same brain into cube-shaped tissue blocks with the same thickness results in thousands of pieces. Unfortunately, no available vibratomes can realize whole-mount precision slicing of human-organ scale tissues. MEGAtome successfully sliced a whole-mounted human brain hemisphere and generated continuous, uniform, and intact 4 mm-thick coronal slabs (**Fig. 3, A and B** and **movie S1**). To whole-mount the human brain hemisphere securely for repeated slicing, we developed a protocol for stable, chemically crosslinked hydrogel embedding of the brains (**Fig. 3A**). With our mounting protocol and MEGAtome, we achieved slicing of an entire human brain hemisphere into 40 consecutive 4 mm-thick slabs within 8 hours (**fig. S4**).

**Fig. 3.**
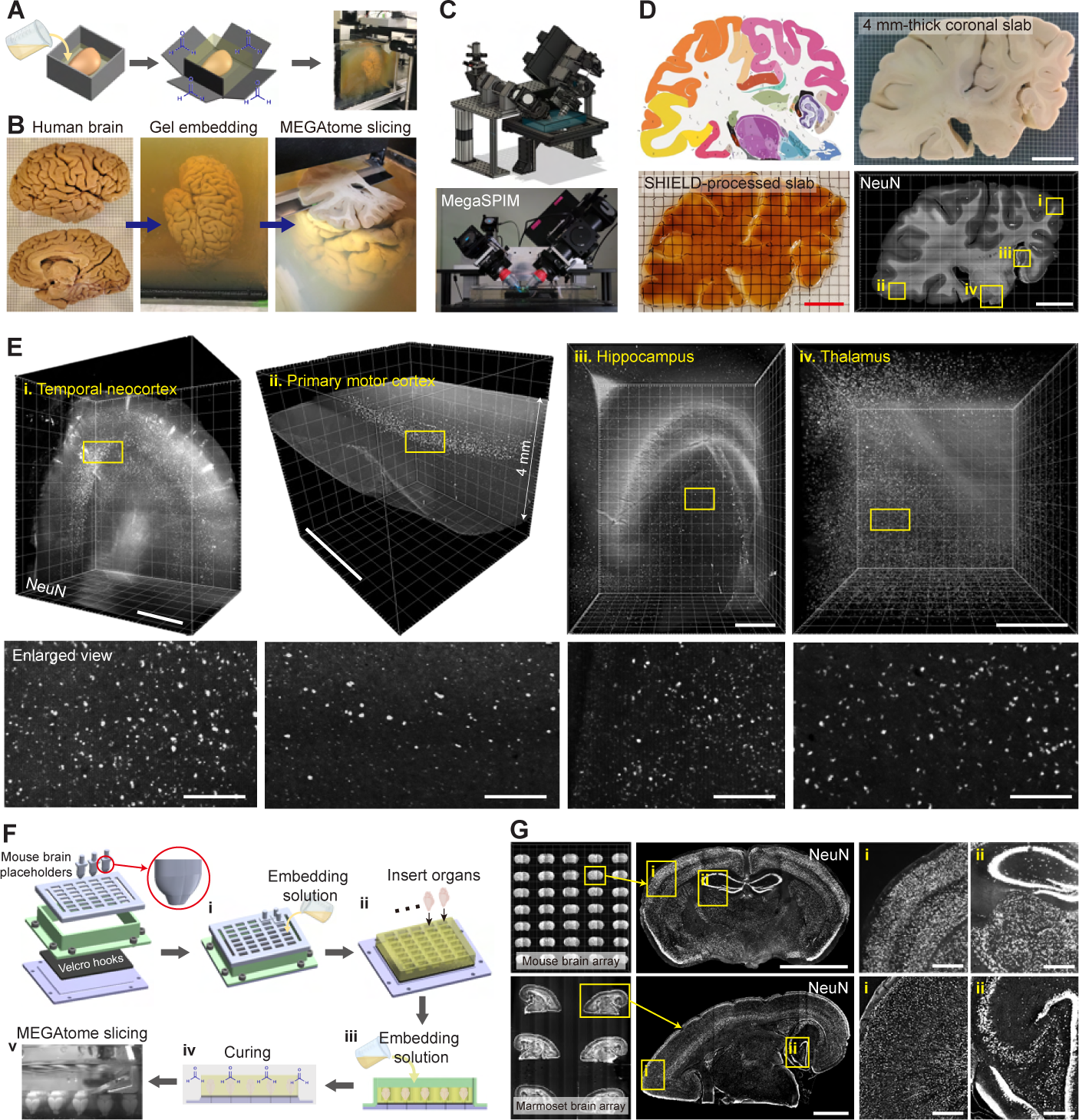
Large-scale slicing and high-throughput imaging of the whole human brain hemisphere and an array of animal organ-scale tissues. (**A**) A schematic drawing of the whole mounting of an intact human brain hemisphere for MEGAtome slicing. To gel-embed a whole human brain hemisphere, liquid gel is first poured onto the hemisphere, then solidified, followed by chemical crosslinking of the gel with the brain in PFA solution. After sufficient crosslinking, the gel with the hemisphere is mounted onto MEGAtome for slicing. (**B**) A banked human brain hemisphere was embedded in gel and subsequently sliced by MEGAtome into 40 consecutive 4 mm-thick uniform slabs. (**C**) MegaSPIM, a rapid, obliquely operated, multicolor light sheet microscopy allows for the imaging of large-scale tissues. (**D**) High-throughput imaging of a 4 mm-thick SHIELD-cleared human brain coronal slab captures all NeuN+ neurons in the slab, encompassing multiple cortex regions, hippocampus, striatum and thalamus. Scale bar, 2 cm. (**E**) 3D image volume and enlarged xy plan views of NeuN+ cells from each brain region. Scale bars, 3 mm (top row) and 200 μm (bottom row). (**F**) Custom-designed tools facilitate the gel-embedding of intact tissues and their subsequent vibratome slicing for areas measuring 72 x 80 mm: (i) Liquid gel is poured over the assembled tools, (ii) animal organs are positioned in the holder, (iii) the organs are completely immersed in extra liquid gel, (iv) the gel-organ mixture is solidified and fixed in PFA, and (v) MEGAtome slices the array. (**G**) MEGAtome, in conjunction with MegaSPIM, enables high-throughput imaging of NeuN+ neurons from the arrays of 35 mouse brains and 6 marmoset hemispheres. Scale bars, 3 mm (second column) and 500 μm (enlarged view). The images were obtained with the 2x/0.1NA objective (E, F, and H). Anatomical annotations in E from Allen Brain Reference Atlases (Adult Human, Modified Brodmann), http://atlas.brain-map.org (*13*).

For imaging large tissue slabs, we developed a multicolor inverted axially-swept light-sheet fluorescence microscopes, termed MegaSPIM (**Fig. 3C**). MegaSPIM can capture multi-scale data with objectives ranging from 2x to 25x, providing near-isotropic resolution and voxel sizes (**fig. S5** and **table S1**). For cellular resolution imaging of full human brain slabs, a 2x/0.1NA objective provides a long WD of up to ∼12 mm. The microscope is equipped with a continuous wavelength, multicolor laser source operating at 405, 488, 561, 647, and 785 nm. Custom-designed microscope stages can accommodate tissue dimensions of up to 180 x 380 mm, enabling single-cell resolution imaging of an expanded coronal human brain hemi slab (**Fig. 4I, i**). Equipping the system with a 16.7x/0.4NA objective enables imaging of expanded tissues at single-axonal resolution (**Fig. 4I, iv**).

**Fig. 4.**
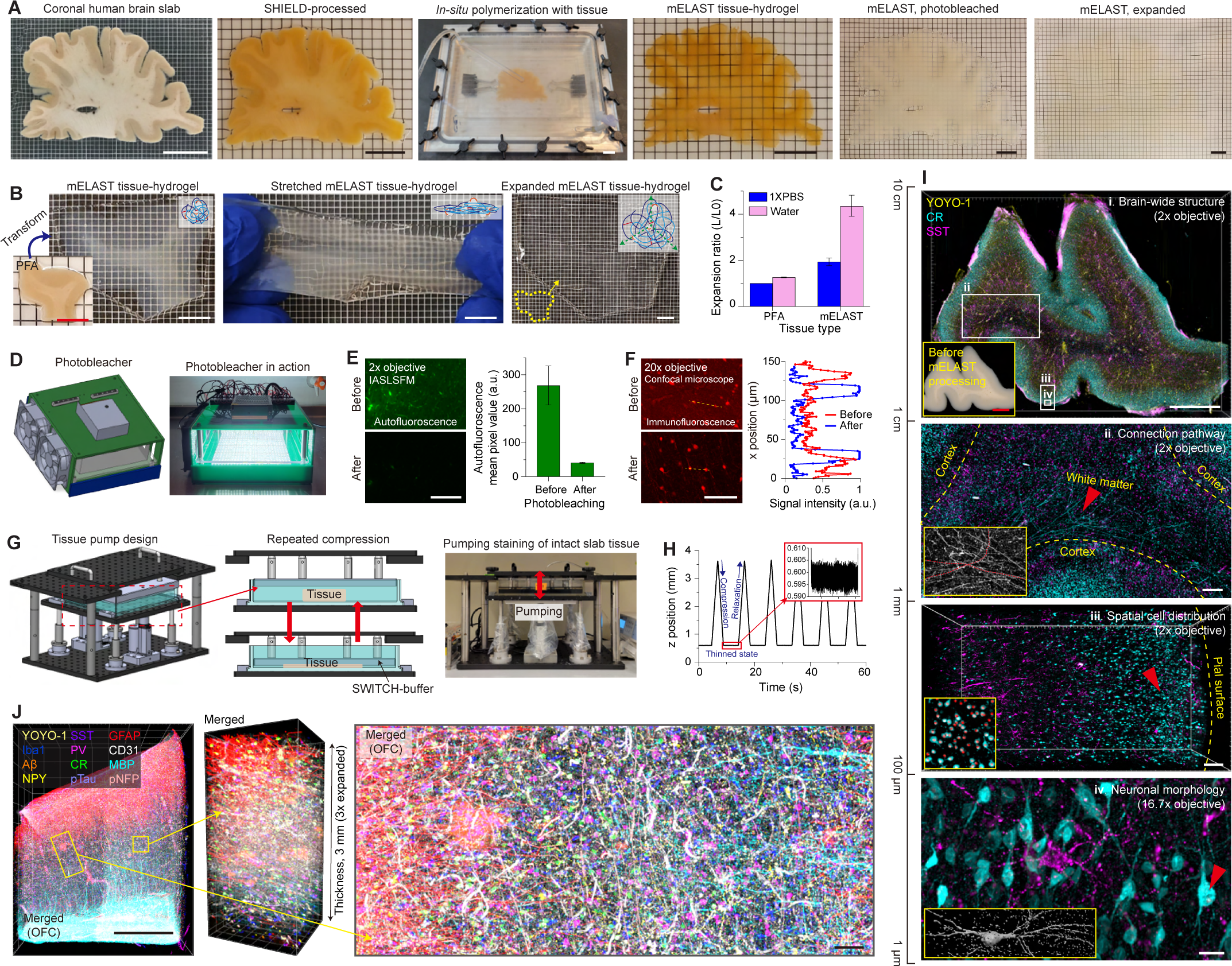
mELAST tissue processing platform for multiplexed multi-scale molecular imaging of thick human brain tissues. (**A-C**) mELAST transforms large-scale tissue into a transparent, permeable, tough-elastic, and size-adjustable material using a novel polymer hydrogel. (**A**) The photos show the processing workflow of an intact coronal human brain slab. Scale bar, 2 cm. (**B**) The mELAST human brain tissue-hydrogel is highly elastic and expandable in water. Scale bar, 1 cm. (**C**) Size-adjustability of PFA tissues and mELAST tissue-hydrogels is compared in 1XPBS and water. L0 is the mean length of the original brain. L is the mean length in 1XPBS or water. (**D**) A photobleaching device is used to remove autofluorescence from lipofuscin and other autofluorescence pigments in human brain tissue. (**E**) Mean background signal values of MegaSPIM images before and after photobleaching are compared. Scale bar, 300 μm. (**F**) Photobleaching allows for high signal-to-noise imaging. Scale bar, 100 μm. (**G**) A human-brain-scale tissue pump enables accelerated clearing, washing, staining, and destaining of large-scale mELAST tissue-hydrogels. (**H**) A graph measures the distance between the pump head and the chamber floor during the operation of the pump. The inset graph shows the accuracy of the device. (**I**) Multi-scale imaging of an mELAST-processed coronal human brain hemisphere tissue. The slab-scale mELAST tissue, stained using SWITCH-pumping with YOYO-1 (a nuclei dye), anti-CR, and anti-SST antibodies, was imaged using MegaSPIM, capturing multi-scale information. Scale bars, 5 mm [i], 1 mm [ii], 200 μm [iii], and 20 μm [iv]. (**J**) Highly multiplexed imaging of an mELAST human brain tissue, simultaneously mapping neuronal and non-neuronal cell subtypes. Scale bars, 2 mm (left figure) and 100 μm (enlarged view). The images were obtained with the 2x/0.1NA (E, I-i, ii, iii, and J), 20x/0.5NA (F), and 16.7x/0.4NA objectives (I-iv).

Using the MegaSPIM system, we demonstrated high-throughput cellular-resolution imaging of a whole human brain hemisphere (**Fig. 3, D and E**). **Figure 3D** shows a SHIELD-preserved and cleared 4 mm-thick coronal human brain slab and annotation of subregions. Following staining with anti-NeuN antibodies, we completed imaging of the entire slab in 6 hours using MegaSPIM, capturing the 3D distribution of NeuN+ neurons (**Fig. 3E**). Using a single MegaSPIM, a whole brain hemisphere can be imaged at single-cell resolution in approximately 100 hours. We believe MEGAtome and MegaSPIM are also suitable for whole-mount precision slicing and high-resolution imaging of other human organs.

Recent large-scale animal studies have greatly increased the need for high-throughput and cost-effective array-based imaging of tissues (*27*, *28*). However, current approaches involve manual handling of individual samples in a piecemeal manner, limiting throughput, increasing labor, and introducing sample-to-sample variations. To address this challenge, we developed a device and protocol to create high-density sample arrays compatible with MEGAtome for high-throughput processing and imaging of various biological samples (**Fig. 3F** and **fig. S6**). The tool ensures that each sample is positioned in the same exact orientation with minimal spacing to maximize slicing and imaging efficiency. With this protocol, we demonstrated high-throughput imaging of NeuN+ cells in mouse (whole brain slabs, 7×5) and marmoset (hemisphere brain slabs, 3×2) tissue arrays, within 4 hours (**Fig. 3G**).

### mELAST enables multiplexed multi-scale imaging of large-scale tissues

Tissue transformation into a macromolecule-permeable and optically transparent hydrogel is a proven approach to investigate complex 3D biological systems (*14*). CLARITY (*15*), which hybridizes tissues with a synthetic polymer hydrogel, facilitates holistic imaging of intact tissues. Tissue expansion strategies (*29–31*) enable super-resolution imaging by imparting superabsorbent properties to tissue-hydrogel hybrids. Transmuting thick tissues into elastic materials using a polymer hydrogel with slip links enables rapid probe transport (*32*). Nevertheless, an integrated technology enabling multiplexed multiscale imaging of the large-scale human brain is still lacking.

To tackle this challenge, we developed a novel tissue processing platform termed mELAST (**m**agnifiable **E**ntangled **L**ink-**A**ugmented **S**tretchable **T**issue-hydrogel) by integrating SHIELD (*33*), SWITCH (*34*), MAP (*35*) and ELAST (*32*) technologies (**Fig. 4A**). SHIELD (*33*) protects endogenous biomolecules by epoxide-based chemical fixatives, ensuring the homogeneous preservation of molecules and tissue architecture. Proteins can be permanently preserved in fully delipidated SHIELD-treated tissue during iterative staining, imaging, and destaining cycles. After SHIELD treatment, mELAST transforms tissues into an elastic, thermochemically stable, and reversibly expandable tissue-hydrogel hybrid via *in-situ* polymerization and subsequent post-processing (**Fig. 4, A and B** and **figs. S7 and S8**) (*32*, *35*). By engineering a synthetic hydrogel with highly entangled and super-absorbent polymeric chains to hybridize biological tissues, we achieved 4.5-fold linear expansion in water while maintaining elasticity, integrity, and high durability of the tissue-hydrogel (**Fig. 4C**). mELAST successfully preserved protein epitopes (**fig. S9** and **table S2**) and no structural information was lost during periodic compression or tissue expansion (**fig. S10**). The processing protocol is highly-scalable and universally applicable even for entire coronal human brain slabs (**Fig. 4A**) and other species and organs without further optimization (**figs. S7C** and **S11**).

Autofluorescence materials in biological tissues, such as lipofuscin in brains (*36*) can degrade the signal and image quality during data acquisition. To address this issue, we built a human brain-scale tissue photobleaching device for uniform photobleaching of thick tissue autofluorescence (**Fig. 4D**). This approach significantly reduced tissue autofluorescence and improved the signal-to-noise ratio (SNR) during data acquisition (**Fig. 4, E and F** and **fig. S12**).

To achieve rapid, uniform, and cost-effective immunostaining, we developed a SWITCH-pumping method that controls reaction kinetics through SWITCH (*34*) during the rapid probe delivery by mechanical thinning (*32*). A custom-designed pump device, scaled for coronal human brain slabs, enabled highly reproducible mechanical tissue thinning (**Fig. 4G**). The tissue pump accelerates antibody delivery by reducing the characteristic tissue thickness and creating convective flow within nanoporous tissue-hydrogel via cyclic mechanical compression (**Fig. 4H**). During pumping-staining of tissue, we modulate antibody-antigen binding reactions using a SWITCH-staining buffer (**fig. S13A**). The SWITCH-pumping method drastically reduces the amount of antibodies required for staining large tissue volume while improving the staining uniformity, even for high copy-number targets, such as parvalbumin (PV), neurofilament heavy chain (NFH), and phosphorylated neurofilament proteins (pNFP) (**fig. S13, B and C**). Automated pumps of various sizes ensure scalable tissue labeling (**fig. S14**).

mELAST, in combination with SWITCH-pumping, enables multiscale molecular imaging of large human brain tissues (**Fig. 4I**). Using the SWITCH-pumping strategy, we achieved completely uniform primary staining of an mELAST-processed whole coronal human brain hemisphere slab (5.8 cm x 4.2 cm x 2.5 mm) with nuclear dye (YOYO-1), anti-calretinin (CR) and anti-somatostatin (SST) antibodies within 12 hours, without any signal gradients in the entire tissue volume (**Fig. 4I, i-iii** and **movie S2**). Multi-scale information was successfully acquired from the 3-fold linearly expanded tissue-hydrogel, including brain wide structure, connection pathway, spatial cell population, and neuronal morphology (**Fig. 4I, iv** and **movie S3**).

Multiplexed imaging allows for detailed analyses of cell types in tissues overcoming the spectral limitations of conventional fluorescence microscopy (*37*). With mELAST, we achieved highly multiplexed labeling and imaging of volumetric human brain tissues (**Fig. 4J**). After each round of staining and imaging, tissue underwent complete antibody elution and then re-stained with the next set of antibodies. The universal applicability of our method enabled uniform staining of thick tissues with most antibodies under identical conditions, without additional optimization. The unprecedented structural stability of the mELAST tissue-hydrogel allowed the samples to withstand extreme mechanical stresses (∼60,000 cycles of 4-fold compression) without tissue damage. After completing 7 rounds of staining and imaging, we coregistered the multi-round datasets at single-cell resolution using a computational algorithm, simultaneously mapping 11 major neuronal and non-neuronal markers in 3D (**Fig. 4J** and **figs. S15 to S17**). Taken together, our innovative tissue-hydrogel technology, mELAST, facilitates large-scale tissue processing, multiplexed molecular staining with a single protocol, and volumetric imaging across various scales.

### Multi-scale interrogation of human brain tissues in Alzheimer’s disease

We applied the mELAST pipeline for interrogating the molecular and structural details of human brain tissues derived from two donors across multiple brain regions (**Figs. 5 and 6**): Donor 1 was a non-demented control (a 61-years-old female), and Donor 2 had Alzheimer’s disease dementia (an 88-years-old female). We sliced the two donor hemibrains using MEGAtome and obtained intact coronal slabs at the same neuroanatomical level to ensure we studied cerebral gyri (e.g., superior frontal gyrus [SFG], rostral gyrus [RoG], cingulate gyrus [CgG], gyrus rectus [ReG], orbital gyrus [OrG], and mid frontal gyrus [MFG]).

**Fig. 5.**
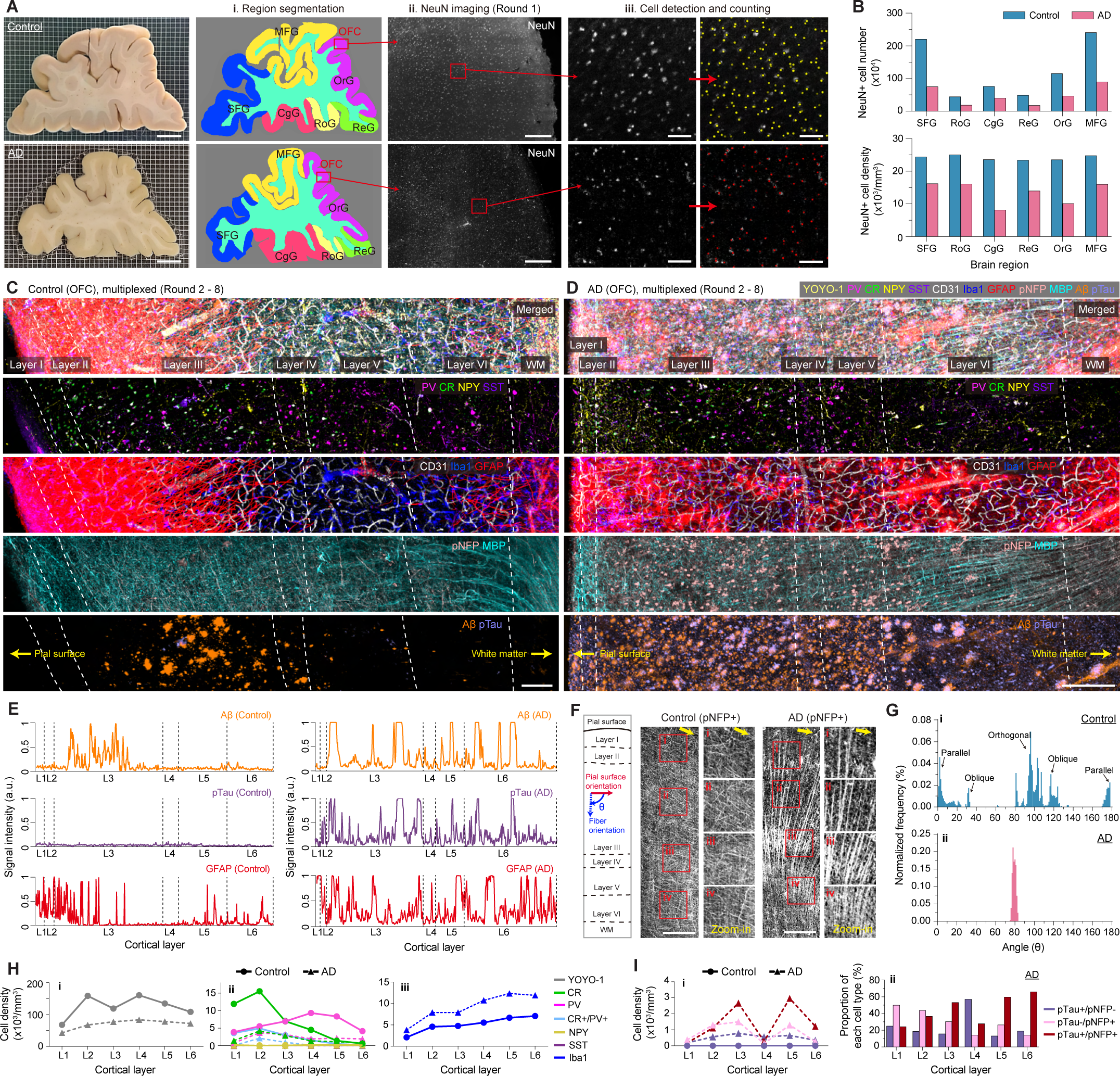
Comparative multi-scale imaging and phenotyping of human brains, Part 1: Single-cell level. (**A-B**) Processing and imaging of intact coronal human brain hemisphere slabs allow for holistic quantitative analysis of individual cells in large tissue volumes. (**A**) Immunolabeled NeuN+ cells in the slabs from both control and AD cases were imaged: (i) Segmented cortical subregions in the whole slab images, (ii) enlarged views of orbitofrontal cortex (OFC) from OrG, and (iii) NeuN+ cells detected and automatically counted for each cortex subregion. Scale bars, 1 cm (first column), 500 μm (third column), and 100 μm (fourth and fifth column) (**B**) The graphs display the difference in number and density of NeuN+ cells across each brain subregion. (**C-D**) Multiplexed staining and imaging of mELAST-processed human brain tissues from the OFC in control and AD slabs, described in Fig. 5A: nuclei (YOYO-1), neuronal subtypes (PV, CR, NPY, SST), astrocytes (GFAP), microglia (Iba1), oligodendrocyte (MBP), axons (pNFP), endothelial cells (CD31), senile plaques (Aβ), and neurofibril tangles (pTau). Scale bar, 200 μm. (**E**) Comparative signal intensity profiles of GFAP, Aβ, and pTau across the cortical column from the two cases. (**F-G**) pNFP+ cortical fiber orientation analysis. (**F**) Representative images show that pNFP+ fibers were oriented differently throughout the cortex of the control and AD tissues (Yellow arrows represent the pial surface direction). Scale bar, 300 μm. (**G**) Histograms of the angular orientation of pNFP+ fibers with respect to the cortical column direction (θ) in the control and AD tissues. (**H**) Layer-specific comparison of the density and distribution of (i) YOYO-1, (ii) PV+, CR+, NPY+, SST+, CR+/PV+ neurons and (iii) Iba1+ microglia in the OFC of the control and AD tissues. (**I**) Analysis of the spatial distribution pattern of cells associated with pTau and pNFP. Layer-specific comparison of (i) cell density and (ii) proportion of each subtype of pTau+/pNFP-, pTau-/pNFP+ and pTau+/pNFP+ cells in each layer. The images were obtained with the 2x/0.1NA objective (A, C, D, and F).

**Fig. 6.**
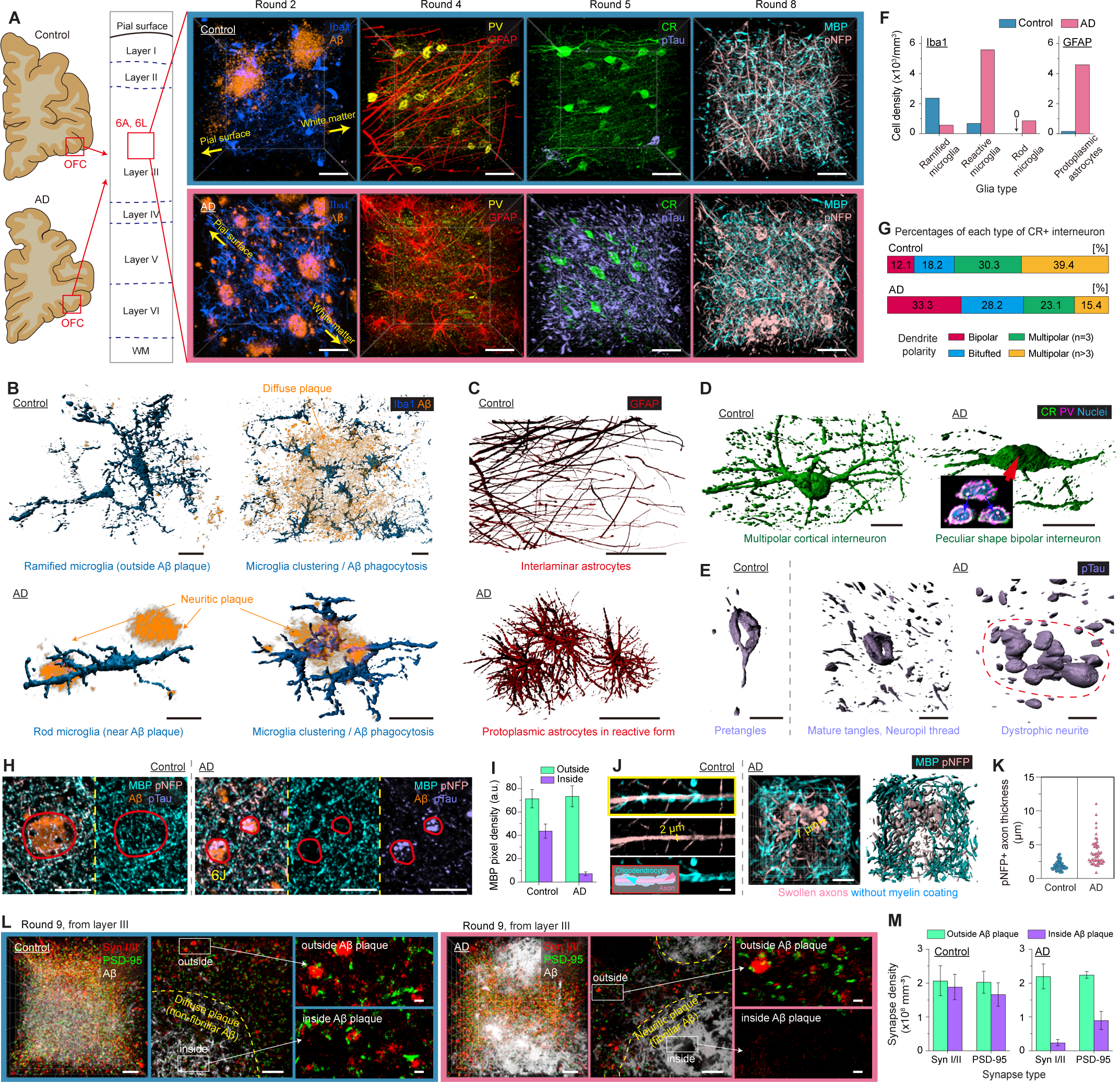
Comparative multi-scale imaging and phenotyping of human brains, Part 2: Subcellular level. (**A**) Fine morphological details of cells in the same expanded mELAST tissue-hydrogels (layer III) described in Fig. 5, C and D. Scale bar, 50 μm. (**B-E**) Comparison of the detailed morphology of each cell type, illustrated with representative 3D rendering images: (**B**) Iba1+ microglia and associated Aβ plaques (scale bar, 20 μm), (**C**) GFAP+ astrocytes (scale bar, 50 μm), (**D**) CR+ and CR+/PV+ neurons (scale bar, 20 μm), and (**E**) pTau+ cells and fibers (scale bar, 20 μm). (**F**) Cell density of each glia type classified based on their morphology. (**G**) Percentages of CR+ interneurons in the control and AD cases, classified by their dendrite morphology: bipolar, bitufted, multipolar (n=3), and multipolar (n>3). (**H-J**) Multi-scale imaging and analysis of axonal damage. (**H**) Representative single-cell resolution images of pNFP+ axons, MBP+ oligodendrocytes, pTau+ fibers, and Aβ plaques. Scale bar, 50 μm. (**I**) Analysis of MBP pixel density from regions inside and outside Aβ plaques in the control and AD tissues. (**J**) pNFP+ axon and MBP+ oligodendrocyte morphology at the subcellular level, represented by the marked region in Fig. 6H. Scale bars, 5 μm (left figure) and 20 μm (right figure). (**K**) Distribution of thickness of representative pNFP+ axons in the control and AD tissues. (**L**) 3D mapping of nanoscopic chemical synapses (Syn I/II and, PSD-95) and Aβ plaques in the same 4.5x expanded control and AD mELAST human brain tissue-hydrogels described in Fig. 6A. Scale bars, 5 μm and 500 nm (zoom-in view). (**M**) Comparison of the density of Syn I/II and PSD-95 in the control and AD tissues, both inside and outside Aβ plaques. The images were obtained with the 2x/0.1NA (H), 16.7x/0.4NA (A-E, and J), and 63x/1.2NA objectives (L).

After SHIELD treatment, the slabs were delipidated, stained with nuclear dyes (YOYO-1) and anti-NeuN antibodies, and imaged at single-cell resolution with the 2x objective (**Fig. 5A**). Nuclear staining enables detection of all nuclei for coregistration of multi-round datasets at the single-cell level. The pan-neuronal marker NeuN facilitates cytoarchitectural parcellations. A custom-built algorithm was used for 3D-segmentation of brain subregions in the volumetric datasets, followed by cell counting and density calculation for each segmented brain region (**Fig. 5B**).

Using our 3D cell phenotyping algorithm, NeuN+ cells were automatically detected in both slabs (**Fig. 5A** and **table S3,** detection accuracy=95.1%) (*38*). Cellular profiling of the cortical areas contained in the entire slabs resulted in a total of 7,464,727 NeuN+ cells for the control case and 2,890,858 NeuN+ cells for the AD case. The NeuN+ cell density was 46.5% lower in AD derived tissue compared to control (13.0×10^3^/mm^3^ vs 24.3×10^3^/mm^3^), a more severe loss than previous reports (22-40%) (*39*). For the control tissues, no regional differences were observed in NeuN+ cell density (ranging from 24.4 to 25.0×10^3^/mm^3^), whereas in AD, the NeuN+ cell density in CgG and OrG was lower compared to other regions.

To further investigate cellular and molecular changes in the regions with high neuronal loss, we performed multiplexed immunohistological imaging of orbitofrontal cortex (OFC) samples from OrG (**table S4**). The OFC is a large region of the frontal cortex known for its role in high-order cognition tasks such as decision-making (*40*). Connectivity in the OFC is impaired in later stages of AD, suggesting its potential contribution to non-memory-related behavioral changes associated with AD (*41*, *42*).

To better understand the cell type- and structure-specific changes in the OFC in AD, we performed multi-round immunostaining and multi-scale imaging of the mELAST-processed tissues, using a set of neuronal subtype markers (parvalbumin [PV], calretinin [CR], neuropeptide-Y [NPY], and somatostatin [SST]), glia subtype markers (ionized calcium-binding adaptor molecule 1 [Iba1] for microglia, glial fibrillary acidic protein [GFAP] for activated astrocytes, and myelin basic protein [MBP] for oligodendrocytes), structural pan-axonal marker (phosphorylated neurofilament proteins [pNFP]), pathogenic protein markers (beta amyloid [A ] and phosphorylated (Ser214 and Thr212 residues) tau [pTau]), endothelial cell marker (cluster of differentiation 31 [CD31/PECAM-1]), and synaptic markers (synapsin I/II [Syn I/II] and postsynaptic density protein 95 [PSD-95]) to map key molecules associated with cellular, structural, connectivity, and pathological processes. The multi-channel datasets were coregistered at single-cell resolution (**Fig. 5, C and D, figs. S14 and S18**, and **movies S4 and S5**).

We observed distinct distribution patterns of the pathology-associated Aβ and pTau proteins in AD compared to control (**Fig. 5, C to E** and **movie S6**). In AD, Aβ senile plaques enveloping pTau+ neurofibrillary tangles were densely distributed throughout the entire cortical area, as well known (*43*). In contrast, Aβ plaques in control were primarily localized in layer III, with very few pTau+ tangles. The distribution of GFAP also differed between the donors. In the control tissue, processes from interlaminar astrocytes at the pial surface were broadly observed to extend into layers III and IV. However, in AD, the density of interlaminar astrocytes was lower, and their processes extended only into the boundary of layers II and III, consistent with previous observations (*44*). Since the interlaminar astrocytes play an important role in stabilizing the physical structure of the brain, the reduction of their density and reach may point to compromised structural stability in AD (*45*). We also observed severe reactive astrogliosis across the cortex in AD, a process characterized by an abnormal increase in the number of astrocytes due to neuroinflammation (*46*, *47*).

Next, we investigated how axonal projection patterns in the OFC are altered in AD by characterizing pNFP+ fiber orientations (**Fig. 5, F and G** and **fig. S19**). In the control tissue, we observed large populations of pNFP+ fibers oriented parallel, orthogonally, and obliquely to the pial surface, while in the AD tissue these fibers were primarily orthogonally oriented, consistent with previous findings on axonal orientation shifts in AD tissue (*48*). In the control tissue, orthogonally oriented fibers were present in all cortical layers, while obliquely and parallel oriented fibers were primarily found in layers I, III, IV, and V. In the AD tissue, all six layers exhibited similar distributions of orthogonally oriented fibers, indicating a shift in pNFP+ axonal orientation in AD.

To study how spatial organization of different cell types is affected in AD, we measured the density of nuclei and each neuron subtype in each cortical layer (**Fig. 5H**, **tables S5 to S7**, and **fig. S20**). Nuclei density measurement showed a lower total cell density in AD compared to control. CR+ (in layers I-III), PV+ (in layers III-V), and CR+/PV+ (in layers I-III) interneurons displayed lower density in AD, a pattern potentially associated with neurodegeneration, impaired homeostasis and calcium-based signaling, consistent with the literature (*49*, *50*). Additionally, the density of Iba1+ microglia was higher (1.81 times on average) across all cortical layers in AD compared to control (**Fig. 5H**) suggesting increased immune activation.

Next, we investigated the relationship between various cell types, phosphorylated neurofilaments (pNFP) and neurofibrillary tangles (NFTs) (**Fig. 5I and table S8**). NFTs, a hallmark feature of AD, are abnormal accumulations of hyperphosphorylated tau (pTau), neurofilaments, and other cytoskeleton proteins, which causes neuronal dysfunction and ultimately death (*48*, *51–53*). We used anti-pNFP antibodies to visualize phosphorylated epitopes on medium and heavy subunits of neurofilament proteins. Unlike control, in which no pNFP+ cell bodies were found, many pNFP+ cell bodies were detected in AD. The pNFP+ cells were highly concentrated in layers III and V (77.1%), the main residence sites for pyramidal cells. No co-localization between pNFP and the interneuronal markers (PV, CR, NPY, and SST) was observed, consistent with reports for selective vulnerability of pyramidal neurons in AD (**tables S6 to S7**) (*52*, *53*). Most NFTs-containing neurons (82.6%) were pNFP+ (pTau+/pNFP+, 54.2%; pTau-/pNFP+, 28.4%) and only 17.4% were pTau+/pNFP-cells, implying that phosphorylation of neurofilament proteins and their accumulation in specific neuronal populations may precede perikaryal pTau aggregation (**table S8**) (*54*).

To further investigate whether subcellular architectures are differentially altered in AD, we performed high-resolution imaging of layer III where Aβ plaques showed high accumulation in both cases (**Fig. 6A**). Most Aβ plaques in AD were neuritic, dense-core plaques known to be closely associated with neuronal and synaptic loss (*55*), whereas Aβ plaques in control were largely diffuse forms (**Fig. 6, A and B** and **movie S7**). As previously observed (*56*), the majority of Iba1+ microglia were in the highly-branched, ramified form (77.6%), but microglia in the reactive form were observed inside and around diffuse Aβ plaques (22.4%) in the control tissue (**Fig. 6F**, left). In the AD tissue, most microglia (79.6%) were reactive, showing that the OFC may exhibit a greater difference in the ratio of ramified/reactive microglia compared to other cortex regions analyzed in the literature (*57*). Additionally, rod-shaped microglia, which are observed in numerous pathological conditions, but whose function and role remain unknown, were exclusively identified in AD (12.2%), but not in control (0%). The elongated body axis of all rod-shaped microglia found in the region of interest were oriented perpendicular to the pial surface (**fig. S21**) (*58*).

A comparative morphological evaluation of GFAP+ astrocytes in layer III was conducted (**Fig. 6, A and C**). In the control tissue, GFAP+ structures were predominantly composed of long, unbranched processes of interlaminar astrocytes located near the pial surface. In AD tissue, these long processes were not detected. Instead, most GFAP+ cells were protoplasmic astrocytes in reactive form, with 31.9-fold higher density of protoplasmic astrocytes in AD versus control (**Fig. 6F**, right) (*44–47*, *59*).

Dendrite morphology analysis of CR+ interneurons revealed differences in arborization patterns between AD and control cases (**Fig. 6, A, D and G**) (*60*). While the majority of CR+ cells in control showed a multipolar dendrite morphology with complex branching patterns (69.7%), bipolar/bitufted neurons with less branching were more numerous in the AD case (61.5%). Interestingly, some bipolar/bitufted CR+ neurons in AD had peculiar shapes, with their cell bodies in very close contact with each other.

In the AD tissue, pTau+/pNFP+ fibers were abundant, indicating the accumulation of pTau in axons, whereas in control, only a few pretangles and neuropil threads were observed (**Fig. 6, A and E**). While pTau-/pNFP+ axons in the control tissue were thin (thickness=1.94±0.64 μm) and largely myelinated, many of the pTau+/pNFP+ axons inside the Aβ plaques in the AD tissue appeared swollen (3.89±2.00 μm) and demyelinated (**Fig. 6, H to K**) (*61*, *62*). In the AD tissue, MBP+ pixel density inside the Aβ plaques was 10-fold lower than that outside the plaques, where in control, we observed only 38.8% reduction in MBP pixel density (71.3±6.9 outside the plaques vs 43.6±5.0 inside the plaques) (**Fig. 6I**) (*63*, *64*).

Finally, to investigate whether synaptic structures are affected in AD, we performed super-resolution imaging of the same region in both AD and control tissues that were linearly expanded fourfold (**Fig. 6, L and M, table S4**, and **movie S8**). We assessed pre- and postsynaptic density (Syn I/II and PSD-95) inside and outside the Aβ plaques. While the synaptic density within the diffuse Aβ plaques was similar to that outside of the plaques in the control case, we found a 10-fold decrease in density of Syn I/II inside the neuritic Aβ plaques compared to that outside (*65–67*). Interestingly, the synapse density outside of the Aβ plaques in AD and control was similar. Collectively, these phenotyping results demonstrate that our mELAST technology pipeline enables extraction of pathological changes at multiple scales from the same clinical human samples.

### UNSLICE enables multi-level connectivity mapping of the human brain

A computational pipeline capable of registering multiple slabs with single-fiber accuracy is required for mapping the 3D neural connectivity across mechanically sliced tissues (**fig. S22**). Previous computational tissue reconstruction methods often work with thin, undeformed, or laterally small tissue sections (*68–71*), making them unapplicable for large, heavily-processed human brain hemisphere slab-scale images. Other methods designed for large mammalian tissue have not achieved accurate reconstruction of densely immunolabeled neurites in human brain tissue at single-fiber resolution (*72*). As such, accurate multi-scale reconstruction of thick tissue volumes at both axonal and organ scales has yet to be accomplished.

To address these challenges, we developed a computational pipeline termed UNSLICE (**U**nification of **N**eighboring **S**liced-tissues via **L**inkage of **I**nterconnected **C**ut fiber **E**ndpoints), which accurately aligns anatomical features at macro-, meso- (blood vessels), and micro-scale (axons) levels. Intensity or feature-based registration methods (e.g., Harris corner detector (*34*)) commonly used for 2D image registration, do not scale well to inter-slab registration of large, thick, highly processed, and immunostained tissue due to deformation. Thus, to achieve accurate inter-slab registration at multiple scales, more relevant and reliable features are required.

To this end, UNSLICE takes advantage of fluorescent labeling of vasculature to match blood vessel endpoints at the cut surfaces to define a smooth deformation field for intact tissue reconstruction. Blood vessels are ideal for inter-slab registration due to their long-range continuity and connectivity, high SNR staining profile, and dense, ubiquitous distribution throughout the brain (*73*, *74*). By ensuring vessel connectivity across consecutive tissue slabs, we achieve effective macroscopic and mesoscopic alignment between multiple slabs.

UNSLICE utilizes the following steps for registering consecutive tissue slabs at vessel resolution (**Fig. 7A** and **figs. S23 and S24**): [1] image pre-processing, [2] tissue surface detection, [3] tissue surface flattening (*75*), [4] identifying matching “anchor points”, [5] detecting, segmenting, and skeletonizing surface vessel endpoints (*76*), [6] matching endpoints between the two surfaces, [7] applying thin plate spline deformation to fuse the volumes, [8] annotating additional endpoints in the newly transformed frame, [9] iteratively repeating steps [7-8] for progressively improved connectivity alignment. This semi-automated process achieves effective and accurate registration between consecutive tissue slabs at the vasculature resolution. Furthermore, by applying this method to image channels containing axonal or neural fiber information, we can fine-tune the vasculature-scale registration to restore axon connectivity across sliced tissue slabs.

**Fig. 7.**
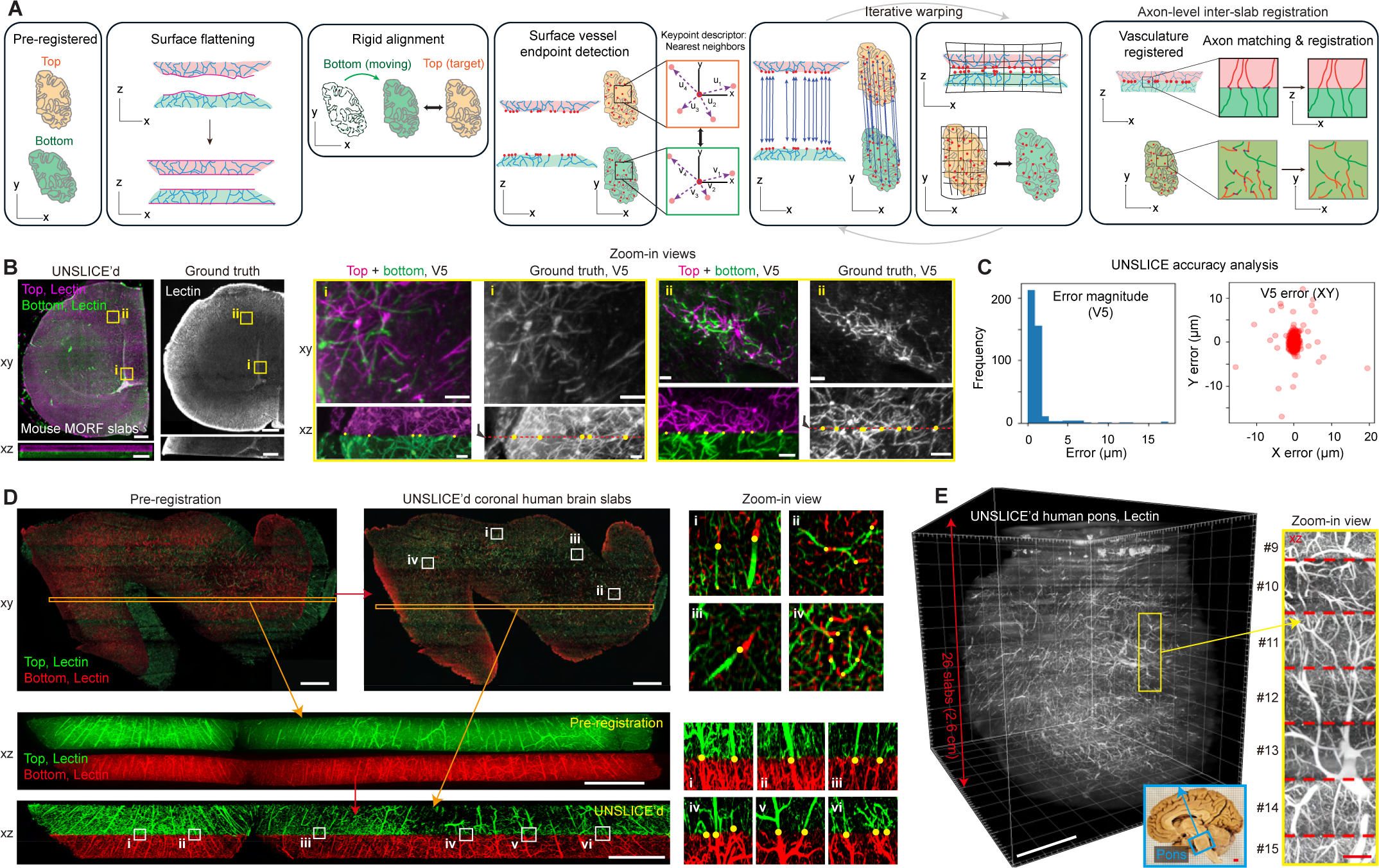
UNSLICE computational pipeline enables accurate, multi-level inter-slab registration of large-scale sliced tissue blocks. (**A**) The multi-level inter-slab registration pipeline, termed UNSLICE, involves pre-registration, surface flattening, rigid alignment, detection of surface vessel endpoints, and iterative warping for vasculature registration. Axon-level inter-slab registration is achieved by starting with the vasculature-registered slabs, then utilizing finer features to achieve progressively more detailed registration. (**B**) UNSLICE enables 3D reconstruction of genetically labeled PV+ neurons in MORF mouse brains. Scale bars, 500 μm (macroscopic view) and 50 μm (zoom-in views). (**C**) Validation of UNSLICE was conducted by quantifying the distance (error) between fiber endpoints at the cut surface in three sampled subsections, based on ground-truth image data before and after slicing at cellular resolution (average error ∼1.29 μm). (**D**) UNSLICE registration of two adjacent coronal human brain hemisphere slabs, based on vascular endpoints (lectin channel) of the consecutive slabs (xy and xz optical cross sections). Scale bar, 2 mm. Inset images show blood vessels connected across the cut surface. Scale bar, 2 mm. (**E**) 3D rendering of a fully reconstructed human pons. Scale bar, 1 cm. A zoomed-in view of the xz plane shows the registered vasculature at multiple interfaces. Scale bar, 500 µm. The images were obtained with the 2x/0.1NA (D and E) and 4x/0.2NA (B) objectives.

To demonstrate UNSLICE’s capability for achieving single capillary and axonal-scale inter-slab registration, we applied it to two different types of sliced tissue datasets, each with an original intact tissue “ground truth”: [1] Genetically labeled PV-Cre MORF mouse hemisphere slabs (**Fig. 7B** and **movie S9**) (*25*, *26*), and [2] NFH labeled expanded mELAST human brain tissue (**fig. S25A**). We quantified the accuracy of inter-slab registration by sampling three subregions per dataset (**Fig. 7C** and **fig. S25B**). We computed the distance between perfectly matched corresponding fiber endpoints in both the ground truth and UNSLICE’d datasets, revealing an average error of 1.29 μm for PV+ neurites in MORF mouse brain slabs and 1.13 μm for NFH+ neurofibers in human brain samples. The results demonstrated the robustness of UNSLICE across varying tissue types, stains, objectives, and datasets, successfully restoring connectivity of PV+ and NFH+ fibers through the cut surfaces.

We applied UNSLICE to consecutive image volumes from millimeter-thick coronal human brain hemisphere slabs (**Fig. 7D**, **fig. S26**, and **movie S10**) and twenty-six human pons slabs (**Fig. 7E and figs. S27 and S28**), demonstrating its feasibility for the scalable reconstruction of the human brain. The computational flattening step in UNSLICE proved especially advantageous in this dataset (**fig. S24**) due to the significantly larger lateral area and irregular shape of the multiple slabs, which posed a challenge in preserving their original shape during tissue processing and imaging. Furthermore, by combining manually defined correspondences for warp initialization with automated vessel endpoint matching in an iterative and parallelized fashion, the user has flexibility in choosing the level of precision required for inter-slab registration. The successful application of UNSLICE, as displayed in the 3D rendering and zoom-in views, showcases the scalability of our pipeline for proteomic and projectomic analysis of large-scale human organ tissue.

### Axonal projectome mapping and pathological analysis of human brain

Vasculature alone is not sufficiently dense to achieve the single-axon resolution registration required for projectome mapping and axon tracing. Therefore, we utilized cell type markers staining for fibers and neurites for axonal resolution registration in addition to vascular staining. The vasculature deformation field was initially used for macroscopic and vessel-resolution registration, and then the fiber endpoints of other cell type markers such as GFAP, PV, and NFH were used to refine the registration results at the fiber-scale (**Fig. 8A**). This refinement enabled multi-channel spatial reconstruction of multiple slabs at single-fiber scale (**fig. S29** and **movie S11**). Additionally, because UNSLICE’s topographic feature matching and outlier removal steps were designed to tolerate both false negatives and positives in fiber endpoint detection, the same vesselness filter, segmentation, and skeletonization steps can be used for automated fiber endpoint detection, even with endpoint detection F1 scores in the 85%-90% range (**fig. S24**).

**Fig. 8.**
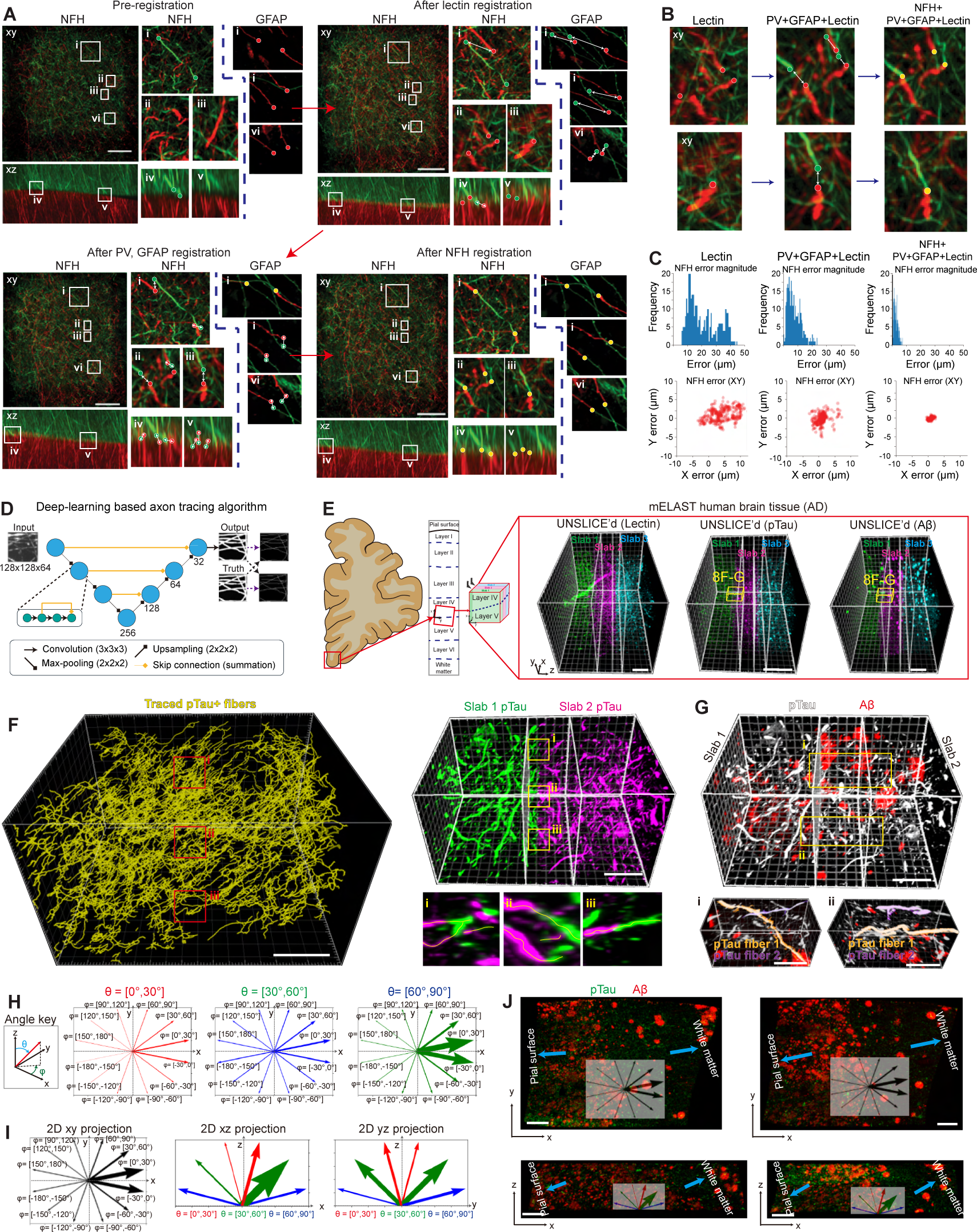
Multi-channel inter-slab registration and volumetric axon tracing for single-fiber level connectivity mapping in human brain. (**A-C**) UNSLICE enables multi-channel reconstruction at single-fiber resolution. (**A**) Demonstration of NFH and GFAP registration at single-fiber resolution within a single field of view of the larger cortical tissue. Scale bar, 100 μm. Inset images depict maximum intensity projections (MIP) of NFH and GFAP in xz and xy planes. (**B**) Fibers can be progressively connected more accurately using multi-channel registration. (**C**) Histogram of absolute errors (top) and x-y lateral connectivity errors (bottom) for NFH+ axonal fibers during each registration step (vasculature registration, PV/GFAP fine-tuning, and NFH fine-tuning). (**D-J**) Synergizing UNSLICE with MEGAtome and mELAST technologies facilitates pTau+ fiber mapping in AD human brain tissue. (**D**) An AD human brain tissue was reconstructed with UNSLICE at single-fiber level. Scale bar, 0.5 mm. (**E**) A deep-learning based axon tracing algorithm performs end-to-end detection of axon centerlines using a residual 3D U-Net architecture. (**F**) High-throughput, volumetric tracing of pTau+ fibers across cut human brain tissue slices. Insets (i)-(iii): Zoomed in xz MIP overlaid with tracing results showing single pTau+ fibers tracing across tissue slices. Scale bar, 150 μm. (**G**) An Aβ (red)/pTau (grayscale) overlay of the same subvolume from (F). Insets (i)-(ii): Zoomed-in subvolumes depicting two segmented long traced pTau+ fibers among Aβ plaques. Scale bar, 150 μm. (**H**) pTau+ fiber orientation histogram of the traced fibers from the subvolume in (F), binned by polar (φ) and azimuthal angle (θ). The thickness of each vector was proportional to fiber counts in each direction. (**I**) Histograms from (H) projected into the xy, xz, and yz planes. (**J**) The majority of the fibers were oriented parallel to the cortical column.

Progressive improvement in the accuracy of individual NFH+ axon connectivity through the tissue sections was observed as additional landmarks from different cell type markers were incorporated (**Fig. 8, B and C**). This multi-stage registration approach also enhanced the connectivity of sparse astrocytic GFAP+ astrocytic fibers. Specifically, the mean error magnitude of NFH+ fiber connectivity was 20.8 μm after the vasculature-based registration, reduced to 7.85 μm with the addition of GFAP and PV landmarks, and finally minimized to 2.35 μm following the inclusion of NFH landmarks. The distribution of x- and y-axis NFH+ fiber connectivity error also shrinks from a spread of 10 μm (in each lateral dimension) after lectin registration, to ∼5 μm after PV/GFAP registration, and finally to ∼1 μm after NFH registration.

Automated and semi-automated neuron tracing and segmentation algorithms have previously been developed for analyzing high-resolution images of single neurons or sparsely labeled groups of neurons (*77–79*). These algorithms are primarily optimized for two types of tasks: [1] subcellular segmentation and tracing at the nanoscale in electron microscopy images or [2] reconstruction of single neuron morphology in fluorescence microscopy images. While these algorithms are suitable for tracing high-resolution, high SNR, and sparsely labeled light microscopy images, to the best of our knowledge, no algorithms are available for high-accuracy tracing of densely packed axons in immunostained tissue volumes. Toward this end, we have developed a deep-learning based axon segmentation and tracing algorithm based on the 3D extension of U-Net architecture (**Fig. 8D**) (*80*). For end-to-end centerline detection, the algorithm employs a centerline-Dice (clDice) loss function (*81*, *82*), allowing us to incorporate parameter-free skeletonization directly into the optimization process.

The UNSLICE and automated axon tracing algorithms enabled the reconstruction of AD human tissue blocks at multiple levels and seamless connection of pTau+ axons across cut surfaces (**Fig. 8E and movie S12**). Multiple pTau+ fibers that traversed the tissue slab seams were reconnected, segmented, and traced (**Fig. 8, F and G** and **fig. S30**). Our pipeline allowed us to analyze the orientation of the traced pTau+ fibers throughout the whole reconstructed volume (**Fig. 8, H and I**). We found that most fibers were oriented in the direction corresponding to an azimuthal angle θ=30° to 60° (with 0° corresponding to perfect alignment with the z-axis and 90° corresponding to perfect alignment with the x-y plane) and a polar angle φ=-30° to 30°, corresponding to fibers aligned with the x-axis. We also estimated the approximate orientation of the cortical column and compared our fiber orientation results to it. Qualitatively, we found that the majority of pTau+ fibers were generally aligned parallel to the cortical column (**Fig. 8J**). Combined with our histological and imaging platforms, this demonstrates that our computational pipeline for inter-slab registration and axon tracing is accurate, scalable, and capable of enabling large-scale connectivity mapping of human brain tissues.

## Discussion

In summary, we developed multiple novel technologies and seamlessly integrated them to establish a scalable platform that enables fully integrated anatomical and molecular phenotyping of cells in human-organ-scale tissues with unprecedented resolution and speed.

MEGAtome realizes precision slicing of ultra-large tissues with minimal loss of information. MEGAtome overcomes the current trade-off in state-of-the-art machines between blade vibration frequency and amplitude. MEGAtome’s multi-DOF system amplifies blade vibrating speed at high frequencies while minimizing the blade mechanical errors. MEGAtome is coupled to a synergistic gel-embedding protocol to facilitate high-throughput imaging of ultra-large-scale samples, preserving biological connectivity information.

The unique expandability and compressibility of the mELAST tissue-hydrogel enables multiplexed, multiscale volumetric imaging of human brain tissues with negligible information loss. With this framework, we could simultaneously capture protein expression, cellular morphology, neural projection, and synapse distribution information from the same brain tissues, a highly desirable feature given inter-individual variation (*83*). The exceptional mechanical and chemical stability, rooted in polymeric networks of the mELAST framework enables adaptive and iterative characterization of cell types and disease markers. This unique feature maximizes the utility of rare and highly valuable human clinical tissues and allows the resulting datasets to evolve continuously beyond the scope of individual studies, enabling the community to jointly probe the processed tissues and continue to extract information from the same samples.

Multi-scale phenotyping of the two human brains (non-demented control and AD dementia) using the mELAST platform elucidated distinctive molecular and structural features of the nervous system affected in AD. Cellular-level imaging allowed for quantitative analysis of various cell subtypes. Subcellular-level imaging of the same brain tissues captured dissimilarities in the morphological properties of cells in areas where different types of Aβ plaques are concentrated. Nanoscopic imaging of these regions revealed altered 3D distribution patterns of synapses within and outside the plaques. The comparative analysis shown in **Figs. 5 and 6** uncovered apparent pathological features in AD tissue, including neuronal loss (*39*, *52*, *53*), tauopathy (*43*, *54*), β-amyloidosis (*55*), astrogliosis (*44–47*), microgliosis (*56–58*), axonopathy (*61*, *62*), myelinopathy (*63*, *64*), and synaptopathy (*65–67*), consistent with literature. Notably, pTau-associated axonal swelling, correlated with severe axonopathy/demyelination and synapse loss in regions surrounded by neuritic Aβ plaques, supports neuroimaging studies that suggest severe damage to the connectivity of the OFC in the late stages of AD (*42*).

Deciphering the neural connectivity can provide long-sought insights into the essential features of brain circuitry and function Yet, mapping neural connectivity in the human brain remains challenging as current interrogation techniques (e.g., viral labeling) are restricted to experimental animal systems (*72*, *84*). To our knowledge, for the first time, we demonstrated reconstruction of inter-slab axonal connectivity at single-fiber resolution in human brain tissue. The iterative nature of our analytical strategy ensures that the accuracy of this reconstruction and tracing will continue to improve as we increase the dimensionality of the integrated 3D datasets by adding additional rounds of 3D fiber-resolution images. The comprehensive circuit mapping enabled by this platform will enable the identification of non-canonical connections, enhancing our understanding of human brain circuitry.

We envision that this scalable technology platform will allow researchers to interrogate a statistically significant number of human organs, adequately covering the spectrum of human demographic diversity and individual variability. Such baseline characterization of cellular, subcellular, and nanoscopic features will provide an immediately useful reference frame for the research community to study disease-specific changes in cell composition, spatial distribution, cellular connectivity, and subcellular architectures. This will advance our understanding of the human organ functions and disease mechanisms to spur development of new therapies.

## Materials and Methods Summary

The postmortem human brain hemispheres were first embedded in a gel for mounting and then sliced using MEGAtome. We assessed the slicing quality by measuring the surface roughness with a VR machine and analyzing the vibration frequency. The sliced human brain tissues underwent SHIELD processing and were hybridized with mELAST polymer hydrogel for permanent protein preservation and transformation of them into elastic, expandable materials. We then immunostained the engineered human brain tissues either passively or actively using a SWITCH-pumping method with antibodies and dyes. The tissues were imaged at multiple scales using a confocal microscope or MegaSPIM, employing various magnification objectives after optical clearing. We repeated the staining, imaging, and destaining processes in multiple rounds for image multiplexing. The imaging data were stitched together, post-processed, and subjected to computational analyses, including brain subregion segmentation, cell/synapse counting, morphological reconstruction/classification, and analyses. The images from the sectioned and engineered tissues were reintegrated using UNSLICE based on the end-point detection and matching at single axonal accuracy. The stitching accuracy of blood vessels and neurofibers in the UNSLICE’d images was evaluated, and their characteristics were analyzed through fiber tracing. Full materials and methods are available in Supplementary Materials.

## Acknowledgments

We thank the entire Chung laboratory for support and discussions. Resources that may help enable general users to establish the methodology are freely available online (http://www.chunglabresources.org). The cartoon in Figure 1 was created with BioRender.com.

## Author contributions

J.P., J.W., W.G., and K.C. conceived the idea and designed the experiments. J.P., J.W., W.G, and K.C. designed the platform and wrote the paper with input from other authors. J.P. developed mELAST technology and performed tissue processing, staining, and imaging of the brain tissues. J.W. developed MEGAtome and tissue-embedding method, and sliced the large-scale tissues. W.G. developed UNSLICE and performed inter-slab stitching of the images. N.B.E. and J.S. developed MegaSPIM. L.K. developed the computational pipeline for processing MegaSPIM images with help from M.M. L.K. developed co-registration of multi-round images. L.G., D.P., M.S., D.C., and L.J.B. developed the deep learning-based axon tracing algorithm and performed axon tracing on datasets with help from W.G. X.G. performed quantitative analysis of cells. C.Z. helped assemble machines and segmented the slab images. S.S.G. developed DANDI platform. S.M. and M.M. managed data storage. L.K. and S.S.G. were involved with standardizing the data. L.K., S.M., and S.S.G. curated, organized, and made available the data on DANDI. M.E.K. helped enhance image quality and performed synapses counting. W.G. stitched and post-processed MegaSPIM images. J.W. developed tissue pumps and photobleachers. S.W.C. and J.P. developed slab-scale SHIELD-clearing protocol. J.P. coregistered multi-round images. J.P. screened and validated antibodies with help from D.H.Y. Y. T. prepared SeTau647-conjugated secondary antibodies. J.P. formulated the RI-matching media. C.S.-A. helped buffer preparation. C.S.P. and X.W.Y. provided MORF mouse brain samples. Q.Z. and G.F. provided marmoset brain samples. M.P.F. provided the human brain tissue specimens. M.P.F., C.D.K. and P.R.H. helped data interpretation. K.C. supervised all aspects of the work.

## Data and materials availability

In this study, the human brain microscopy image data from the SHIELD and mELAST tissues acquired using MegaSPIM for the BICCN (RRID:SCR_015820) by Juhyuk Park and colleagues accessed/deposited in the DANDI Archive (RRID:SCR_017571) under accession number DANDI:000108. Data may be accessed at: https://dandiarchive.org/dandiset/000108. Neuroglancer view of the data on DANDI: https://biccn.github.io/Quarterly_Submission_Receipts/000108-dashboard.html <u>(The overlapped chunks in each slab are “stitched” using the rigid transform available with each chunk)</u>. All other datasets used in this paper are available upon request. Code used for the paper can be found at the following Github repositories: https://github.com/chunglabmit/spimstitch (post-acquisition pre-processing e.g., illumination correction, tile stitching, and file conversion), https://github.com/chunglabmit/unslice (UNSLICE), and https://github.com/chunglabmit/multiround-alignment-ui (multi-round image co-registration). The cell counting code will be released upon institutional review for open-source. The axon tracing code will be released upon institutional review for open-source.

## Supplementary Materials

Materials and Methods

Figs. S1 to S30

Tables S1 to S8

References

Movies S1 to S12

