## Supplementary Materials for "Integrated platform for multi-scale molecular imaging and phenotyping of the human brain"

Juhyuk Park *et. al.*

**This PDF file includes:**

Materials and Methods
Figs. S1 to S30
Tables S1 to S8
Captions for Movies S1 to S12
References

**Other Supplementary Materials for this manuscript include the following:**

Movies S1 to S12

### Materials and Methods

Mechanical configuration of MEGAtome

MEGAtome includes two main modules: a sample mounting and feeding platform and a blade vibration generation and control mechanism. The sample mounting and feeding platform was configured through two commercial high-precision linear stages (M-531, M-511; Physik Instrumente): one stage was oriented horizontally and the other one was oriented vertically. The vertical stage was mounted on the moving piece of the horizontal stage through an L-shaped adapter bracket. A sample carrying platform, where the sample is fixed on, was mounted on the vertical stage. A pair of linear guiding rails and bearings were configured as a passive linear stage. It was connected to the M-531 linear stage through a moving platform, on which sit the buffer solution tank (Fig. 2B). Thus, the large weight of the solution-filled tank was stably supported by both the M-531 stage and the passive stage. When the sample was mounted, the sample was fully immersed in the buffer solution in the tank. During operation, the horizontal stage feed the mounted sample towards the vibrating blade to realize slicing. It returned the sample to its starting position after a slice was generated. Then the vertical stage elevated the sample to the next cutting position. Thickness of the slices was determined by the step distance of the vertical stage. This process was repeated until the slicing process was finished.

The blade vibration generation and control mechanism mainly include three pairs of flexures ($F_1$, $F_2$ and $F_3$), a blade mounting assembly, a voice coil actuator (NCC03-15-050-2X; H2W Technologies), and two sets of laser displacement sensors (ZX2; Omron). Flexure $F_1$, $F_2$ and $F_3$ are thin aluminum beams that are flexible in the blade vibrating direction. Flexure $F_1$ connects an intermediate mass, where the vibration input was exerted, and the stationary ground. Flexure $F_2$ connects the intermediate mass and the blade mounting assembly, where the vibration output was expected. Flexure $F_3$ connects the blade mounting assembly and the station ground vertically to eliminate the out-of-plane displacement. All the flexures were machined with CNC process from 7075 aluminum alloy. The blade mounting assembly accommodates both off-the-shelf blades and custom-made blades as long as 120 mm. The blade mounting assembly was also designed to be lightweight to avoid lowering the natural frequency of the mechanism. The voice coil actuator consisted of a stationary magnet and a moving coil. When given an oscillating current, an oscillating force was generated to the moving coil which was attached to the blade mounting assembly. The amplitude and frequency of the force is linear to the amplitude and frequency of the current. The laser displacement sensors were used to measure the blade vibration input and output feedback.

120 mm long blades were machined by Cadence through CNC process. The blades were made from 316 stainless steel, the blade thickness was 0.25 mm, and the hone angle was 13°.

Electronic system configuration of MEGAtome

To control the voice coil actuator, we used an I/O device (USB-6363; National Instruments) to generate analog current output signal. The signal generator was connected to a current mode driver (LCAM-5/15; H2W Technologies), which also drives the actuator. The same I/O device was used to receive analog input signal from the laser displacement sensors for feedback signal acquisition. The sensing and actuating system was controlled by a custom-developed program written in LabView. This program allows adjusting signal frequency and amplitude. We also set up two cameras (BFS-U3-13Y3M-C USB3; FLIR) to monitor the sample slicing process from top and side views.

Single-DOF system vs Multi-DOF system

Current commercial and custom vibratomes that are compliant flexure-based can achieve relatively high blade vibration speed but have their inherent limitations. They are designed based on single-degree-of-freedom systems (Fig. 2F, left) (*21*, *24*), in which vibration input is directly applied to a blade mounting assembly (M) which is constrained by a spring component (K). Such a system has a trade-off between its vibration frequency and amplitude: a system with high spring stiffness can operate at a high frequency but is limited to low amplitude due to the allowable material deformation in the spring; a system with low spring stiffness can achieve large amplitude but only at a lower frequency. Additionally, direct vibration drive can cause noise and mechanical error due to mechanical misalignment. On the other hand, MEGAtome's multi-DOF design is not subjected to the trade-off as its forced vibration response involves multiple design elements: two masses ($M_1$, $M_2$) and three springs ($K_1$, $K_2$, and $K_3$). The multi-DOF system can be modeled as a forced mass-spring vibration system that consists of two moving masses ($M_1$, the intermediate mass connected to the VCA; $M_2$, the mass of the blade and the blade mounting assembly) and three springs ($K_1$, $K_2$, and $K_3$ representing their stiffnesses, respectively). A sinusoidal force is applied to the intermediate mass $M_1$ to indirectly drive the blade mounting assembly $M_2$. The system has two resonant frequencies, near where $X_1$ (vibrating amplitude of $M_1$) and $X_2$ (vibrating amplitude of $M_2$) are significantly increased due to mechanical resonance. Taking advantage of this property, we can drive the multi-DOF system near its higher natural frequency (~140 Hz) to achieve high blade vibration frequency as well as amplified vibration amplitude.

Modeling of the vibration system

The mechanical design of MEGAtome can be modeled as a multi-DOF vibration system in its vibrating direction. Because the purpose of the simulation is to find patterns of the system's response and the resonant frequencies instead of finding the exact vibration displacement, it is reasonable to assume the material damping can be neglected. Thus, the forced equation of motion for the system can be written as:

$$\begin{bmatrix} M_1 & 0 \\ 0 & M_2 \end{bmatrix} \begin{Bmatrix} \ddot{x_1} \\ \ddot{x_2} \end{Bmatrix} + \begin{bmatrix} K_1 + K_2 & -K_2 \\ -K_2 & K_2 + K_3 \end{bmatrix} \begin{Bmatrix} x_1 \\ x_2 \end{Bmatrix} = \begin{Bmatrix} F(t) \\ 0 \end{Bmatrix} \qquad (1)$$

where $M_1$ and $M_2$ are moving masses, $K_1$ , $K_2$ and $K_3$ are stiffnesses along the x-axis of flexure $F_1$, $F_2$ and $F_3$. $F(t)$ is the force exerted by the voice coil actuator. Given that $F(t) = F_0 cos\omega t$, the forced response can be written as:

$$\begin{Bmatrix} x_1 \\ x_2 \end{Bmatrix} = \begin{Bmatrix} X_1 \\ X_2 \end{Bmatrix} cos\omega t$$

$$\begin{Bmatrix} X_1 \\ X_2 \end{Bmatrix} = \frac{1}{A} \begin{Bmatrix} (K_2 + K_3 - M_2\omega^2)F_0 \\ K_2 F_0 \end{Bmatrix}$$

$$A = \begin{vmatrix} K_1 + K_2 - M_1\omega^2 & -K_2 \\ -K_2 & K_2 + K_3 - M_2\omega^2 \end{vmatrix} = M_1 M_2 (\omega^2 - \omega_1^2)(\omega^2 - \omega_2^2) \quad (2)$$

where $\omega_1$ and $\omega_2$ are the two positive eigenvalues of Equation (2), representing the first mode and second mode natural frequency. When the excitation frequency $\omega$ is close to $\omega_1$ or $\omega_2$, the vibration amplitude $X_1$ and $X_2$ increase significantly.

The single-DOF vibration system models of current commercial vibratomes can be divided into two categories depending on the type of blade motion guide. Vibratomes that use traditional linear bearings, e.g., the ball bearing cartridge and guide rail combination, to constrain the blade vibration usually have large moving mass and large damping, resulting in low frequency (< 60 Hz) vibration generation. The other types of vibratomes, such as Leica VT1200, adopt flexures as their linear motion guide. They can achieve relatively high frequencies, such as 85 Hz. Such a system consists of a moving mass *M* and a spring component that has stiffness *K*. The forced equation of motion for the system can be written as:

$$M\ddot{x} + Kx = F(t) \quad (3)$$

Given that $F(t) = F_0 cos\omega t$, the forced response can be written as:

$$x = Xcos\omega t$$

$$X = \frac{F_0}{K(1-(\omega/\omega_0)^2)} \quad (4)$$

where $\omega_0 = \sqrt{K/M}$ is the natural frequency of the system. Amplified vibration can be achieved near the mechanism's natural frequency $\omega_0$. However, when driven beyond $\omega_0$, $X$ decreases drastically. Increasing system stiffness can increase the driving frequency but lower the amplitude of the response.

<u>Simulation of forced vibration response</u>

Simulation of forced vibration response for both multi-DOF and single-DOF systems were performed in Python based on Equations (2) and (4). To obtain flexure stiffnesses of the mechanism, finite element simulation was performed in SolidWorks with the real geometric parameters of the flexures. The flexures of MEGAtome are 1 mm thick beams machined with 7075 aluminum alloy. The simulated $K_1$ , $K_2$ and $K_3$ are 27.5 N/mm, 17.8 N/mm, and 20 N/mm, orderly. $M_1$ is 0.1 kg, including the mass of the moving coil of the actuator and the moving mass connecting to the coil. $M_2$ is 0.06 kg, including the mass of the blade mounting assembly. For the simulation of a single-DOF system, it is assumed that $K = K_1$, $M$ = 0.14 kg which includes the mass of the moving coil, the blade mounting assembly, and the connecting piece between the two parts. It is assumed that $F_0$ = 10 N for the simulation of both multi-DOF and single-DOF systems.

<u>Characterization of MEGAtome operation</u>

To characterize the blade vibration output vs input of MEGAtome, two laser displacement sensors (ZX2; Omron) were used to measure the displacement of $M_1$ and $M_2$. The sensors were mounted to the stationary ground of the machine. They were configured to emit laser beams perpendicular to the surfaces being measured. The sensors' resolution was 1.5 μm and the measurement range was ±5 mm. The same sensor was used to measure the real blade vibration output of Leica VT1200.

To characterize the blade out-of-plane displacement vs in-plane displacement of MEGAtome and Leica VT1200, three high-precision capacitance probes (CPL290; Lion Precision) were used to measure out-of-plane displacement of the blade at three different locations, including the left edge, right edge, and the center of the blade. The non-contact probes detect small displacement by sensing changes of the capacitance of the electric field between the probe and the conductive object surface. The resolution of the sensors was 5 nm RMS. At the same time, a laser displacement sensor was used to measure the in-plane blade vibration. Measurements for both machines were performed with the same 80 mm-long blade (D554G50; Sturkey).

Acquisition of human, mouse, and marmoset tissues
Human tissues. Whole human brain hemispheres in formalin solution were obtained from Massachusetts General Hospital. The brains were collected and banked in accordance with approval from the local Institutional Review Board. The formalin solution was replaced with 1XPBSN made with modified 10X PBS solution, pH 7.2 (2P1072, Teknova, CA, USA) and 0.02% sodium azide (N, NaN3, S2002, MilliporeSigma) upon arrival, and the hemispheres were stored at 4°C until use.

Mouse tissues. All experimental protocols were approved by the MIT Institutional Animal Care and Use Committee and the Division of Comparative Medicine and were in accordance with guidelines from the National Institute of Health. PFA fixed wild-type female mouse brains (10 weeks old) for the brain array experiment (Fig. 3F-G) were purchased from Hilltop Lab Animals.

Marmoset tissues (*25*). All animal experiments were approved by the Institutional Animal Care and Use Committee of Massachusetts Institute of Technology and were performed under the guidelines from the National Institute of Health. Adult common marmosets (2-4 years old) were housed in AAALAC-accredited facilities. The housing room was maintained at 23.3±1.1 °C in the relative humidity of 50±20 %, and in a 12-hour light/dark cycle. The animals were housed in dedicated cages with enrichment devices and had unrestricted access to food and water. For histological examinations, the animals were deeply sedated by intramuscular injection of Ketamine (20-40 mg/kg) or Alfaxalone (5-10 mg/kg), followed by intravenous injection of sodium pentobarbital (10-30 mg/kg). When pedal withdrawal reflex was eliminated and/or respiratory rate was diminished, animals were transcardially perfused with 0.5 ml 1000 IU/ml heparin and 100-200 ml cold 1XPBS by gravity. Then the descending aorta of the animals was clamped and a peristaltic pump was used to infuse another 200-300 ml ice-cold PFA perfusion solution (4% PFA(w/v) in PBS). Brains were removed from the skulls and subject to post-PFA fixation.

Human brain tissue gel embedding and slicing
A five-side container was made for human brain whole hemisphere gel embedding with inside dimension of 250 mm [length] x 200 mm [width] x 150 mm [height]. The container was made from ¼ in. clear acrylic sheets that were laser cut to size. The container's 5 sides were taped together so that they can be detached later. An acrylic substrate (195 mm x 115 mm) that has Velcro hooks attached to one side was placed inside the acrylic container, standing against the 200 mm x 150 mm wall. 1XPBS with 10% gelatin solution (Gelatin Type B, 9000-70-8, Fisher Chemical) was made and poured into the

container to about 15 mm in height. Gelatin was then cured at 4°C for about 2 hours until the gel became solidified. A human brain hemisphere, which had meninges removed from, was then placed on top of the gel, with the midsagittal plane facing down. More 1XPBS with 10% gelatin solution was poured into the container until the solution was about 15 mm above the top of the hemisphere. The gel was then brought to cure at 4°C for about 24 hours. Next, the container's four side walls were detached, having the side of the gel block exposed. The gel block was then placed inside a large container. 4% paraformaldehyde (PFA) solution was made using a 32% solution (15714-S; Electron Microscopy Sciences) and was filled in the container to fully immerse the gel block. The gel block was fixed for 7 days at room temperature (RT), while PFA solution was refreshed or 1-2 times. After fixation, the gel block was washed in 1XPBSN using Gibco 10X PBS (70011-044; Life Technologies) for 1 day at RT. Then, the gel block was mounted onto the sample platform inside the solution tank of MEGAtome in coronal slicing position. The solution tank was filled with 1XPBSN solution to submerge the sample. The custom made 120 mm long stainless-steel blade was used to slice the whole hemisphere sample. The sample feeding speed was set to 0.2 mm/s for 4mm slices cutting 0.1 mm/s for 1 mm slices cutting.

To gel embed and slice smaller human brain tissues such as cerebellum, pons and spinal cord, the same general protocol was followed as the whole hemisphere embedding and slicing process, except the embedding container was made to be 60 mm [length] x 50 mm [width] x 50 mm [height]. PFA fixation was 2 days at RT for these small samples.

Gel embedding and slicing for arrays of mouse and marmoset brain tissues

To embed an array of mouse brains in the gel, a custom designed container was assembled by fastening a container base piece with a container wall piece by screws. The inside dimension of the container was 72 mm x 98 mm x 28 mm. The base piece was CNC machined from aluminum with tapped holes. The wall piece was 3D printed. After the container was assembled, a plastic substrate, 71 mm x 97 mm in size, that has Velcro hooks attached was placed in the container. Next, a rack was placed on top of the container, sitting on the top plane of the container wall. The rack has patterned rectangular cutouts to fit 5 x 7 placeholders. The placeholders were designed to match the shape of a typical mouse brain on the bottom part. The top part of the placeholders was in rectangular shape to match the cutout size of the rack. 35 placeholders were inserted to the rack hanging in the container. 1XPBS with 10% gelatin was poured into the container. The gel was cured at 4°C for about 2 hours. Placeholders were then removed one by one; the rack was also removed afterwards. Mouse brain fitted cavities were formed in the gel, where 35 mouse brains were positioned. 1XPBS with 10% gelatin was poured inside the container to fully submerge the brains. The gel was cured at 4°C for about 2 hours. After the gel solidified, the container wall was disassembled from the base. The gel block that contained 35 brains were fixed in 4% PFA solution for 2 days, followed by 1XPBS washing for 1 day. The sample was then mounted on MEGAtome. 80 mm-long off-the-shelf blades (D554G50, Sturkey) were used for slicing. The sample feeding speed was set to 0.2 mm/s for generating slices with >1 mm thickness, it was set to 0.1 mm/s for 200 μm and 500 μm slices.

Based on the same idea of mouse brain array embedding, we demonstrated embedding and slicing of an array of 6 marmoset brain hemispheres. The marmoset brain hemisphere-shaped placeholder was designed based on the 3D surface scan of a marmoset brain left hemisphere.

Tissue slice surface characterization

A non-contact surface profiler (Keyence, VR-5000) was used to scan the surface of tissue slices and quantify surface roughness. When the profiler illuminates the slice surface with structured light beams, height variance on the surface causes distortion to the structured light beams. When receiving the distorted reflected light beams, the profiler can recreate the surface profile and calculate surface roughness parameters, such as Ra, which is the average of the microscopic peaks and valleys of the surface area being measured. Surface profile figures were generated in the VR-5000 Analyzer software.

Chemicals

The SHIELD-perfusion solution (*32, 33*) consists of 2.5 w/v% of polyglycerol 3-polyglycidyl ether (GE38 or P3PE, CVC Thermoset Specialties of Emerald Performance Materials, Moorestown, NJ, USA) and 4% paraformaldehyde (PFA) made from a 32% stock solution (15714-S, Electron Microscopy Sciences, Hatfield, PA, USA) in a buffer composed of 0.1 M sodium phosphate buffer (diluting P2072, Teknova, Hollister, CA, USA) and 50 mM sodium chloride (diluting S5845, Teknova) (*32, 33*). After vortex mixing all the chemicals in a 50 ml conical tube, the clear supernatant was used following centrifugation for 10 minutes at 4°C and 7,200 r.c.f. The SHIELD-OFF solution is identical to the SHIELD perfusion solution, except that it does not contain PFA. The SHIELD-ON solution is a weak-base physiological buffer solution (pH~9) that triggers epoxy-based crosslinking of biomolecules and is made from 0.05 M sodium carbonate (S7795, MilliporeSigma, St. Louis, MO, USA) and 0.05 M sodium bicarbonate (S5761, MilliporeSigma). The post-SHIELD solution was prepared by dissolving 2 w/v% GE38 in the SHIELD-ON solution.

The mELAST solution was composed of 40 w/v% acrylamide monomers (AA, A9099, MilliporeSigma) 0.005% bis-acrylamide crosslinkers (BisAA, 2% bis solution, 161-0142, Bio-Rad Laboratories, Hercules, CA, USA), and 0.05% VA-044 thermal initiators (27776-21-2; Wako Chemicals, Richmond, VA, USA) for 1 mm thick tissues and 0.03% VA-044 for mouse whole-organ tissues. All reagents except VA-044 were dissolved in 0.5X PBS (phosphate-buffered saline), which was diluted from a modified 10X PBS solution, pH 7.2 (2P1072, Teknova, CA, USA) and kept at 4°C until use.

The NaCh clearing solution for passive delipidation of SHIELD-processed tissues was made by dissolving 12 w/v% cholic acid sodium salt (NaCh, J13630A1, Fisher Scientific), 150 mM lithium hydroxide monohydrate (LiOH, 43171, Alfa Aesar, Haverhill, MA, USA), 250 mM boric acid (B6768, MilliporeSigma), 70 mM sodium sulfite (S0505, MilliporeSigma) in deionized water (DIW). The SDS clearing solution for protein denaturation, hydrolysis, and destaining was made by dissolving 200 mM sodium dodecyl sulfate (SDS, 75746, MilliporeSigma) in the same buffer solution, replacing 12% NaCh. 1X PBS, 1X PBST, 1XPBSN or 1XPBSTN was made by diluting 10X PBS into DIW with or without 0.02% sodium azide (N, NaN3, S2002, MilliporeSigma) and 1%

TritonX-100 (T, X100, MilliporeSigma). 5 v/v% normal donkey serum (NDS, ab7475, abcam) was included in all the staining buffers for preblocking. The SWITCH staining solutions (PBSNaCh) consisted of 1X (for passive staining) or 0.2X (for SWITCH-pumping staining) PBS with 5% NaCh and 0.02 w/v% NaN3. dPROTOS, a refractive index (RI) matching solution (RI~1.52, pH~7.2) was composed of 530 g iohexol, 150 mL 10 w/v% 4-methoxyphenol (M18655, MilliporeSigma) in dimethyl sulfoxide (DMSO, D8418, MilliporeSigma), 150 mL 2,2'-thiodiethanol (TDE, 166782, MilliporeSigma), 200 mL DIW, and 150 ul 10 w/v% triethanolamine (TEOA, T58300, MilliporeSigma).

Human brain samples acquisition

Postmortem PFA-fixed human brain hemispheres were obtained from the Massachusetts General Hospital brain bank. Donor 1, a non-demented control, was a 61-year-old female who had no clinical evidence of neurological impairment and showed low levels of AD neuropathologic changes (ADNC). Donor 2, diagnosed with Alzheimer's Disease (AD) dementia, was an 88-year-old female with clinically suspected and neuropathologically confirmed AD dementia.

SHIELD tissue processing

All the experimental protocols using mice were approved by the MIT Institutional Animal Care and Use Committee and the Division of Comparative Medicine and were in accordance with guidelines from the National Institute of Health. Wild-type (WT) and Thy1-eGFP-M line mice (2-4 months old) were housed under a 12 hours' light/dark cycle with unrestricted access to food and water. Mice were transcardially perfused with 20-30 mL of ice-cold 1X PBS at a flow rate of 5 ml/min. After confirming that the fluid running from the mice was completely clear, 35 mL of ice-cold SHIELD-perfusion solution was transcardially injected to the mice at the same flow rate. Following perfusion, the mouse organs were extracted from and incubated in the remaining 15 mL SHIELD-perfusion solution for 48 hours with gentle shaking in a 4°C cold room. The mouse organs were transferred to the SHIELD-OFF solution and incubated there for 24 hours with gentle shaking in the cold room, followed by transfer to the SHIELD-ON solution in the 37°C incubator for another 24 hours with gentle shaking. The SHIELD processed mouse organs were washed with 1XPBSN for 2 days with the solution refreshed three times.

After vibratome slicing the SHIELD-processed mouse brain into 0.2- or 1-mm thickness, the tissues were passively cleared in the NaCh clearing solution at 37°C for 1 day (for 0.2 mm sections) or 2 days (for 1 mm sections). The whole mouse organs were delipidated in the SDS clearing solution at 45°C for 14 days, with the buffer being refreshed every 2 days.

Coronal human brain slabs, human pons slabs, or organ-scale tissue arrays, sliced with MEGAtome, were incubated in the post-SHIELD solution in the cold room (4°C). The incubation period was 2 days for 1 mm thick slices and 4 days for 2 mm thick slices, with the solution refreshed once. 4 mm thick slabs were first incubated in the SHIELD-OFF solution (with 2% GE38, instead of 2.5%) for 4 days and then in the post-SHIELD solution for 4 days in the cold room (4°C). Subsequently, the pre-incubated 1-, 2-, or 4-mm thick tissues were directly transferred to a fresh post-SHIELD solution at room temperature for the 'reaction ON' step for 24 hours. The remaining GE38 in the

SHIELD-processed human brain slabs was washed out with 1XPBSN at 37°C for 1 day (for 1 mm thick slabs), 1.5 days (for 2 mm thick slabs) or 2 days (for 4 mm thick slabs), refreshing the buffer solution three times per day.

The SHIELD-processed brain slabs were delipidated in the NaCh clearing solution at 45°C for 5-7 days (for 1- or 2-mm thick slabs) or 21 days (for 4 mm thick slabs), with the solution refreshment every 3 days. After clearing, the remaining NaCh molecules in the brain tissues were washed out with 1% 1XPBSTN at 45°C for 2-3 days (for 1- or 2-mm thick slabs) and for 4 days (for 4 mm thick slabs). The tissues were then stored in 1XPBSN at 4°C until further use.

mELAST tissue-hydrogel processing

SHIELD-cleared tissues were incubated and equilibrated in the mELAST solution in the cold room (4°C), for 1 day (for 1 mm thick mouse brain tissues and small human brain tissue slices), 2 days (for 1 mm thick intact coronal human brain slab), and 4-5 days (for intact mouse organs) to ensure sufficient infusion of monomers deep within the tissues. For small tissues (e.g., mouse brain tissues), polymerization was conducted in a gelling cartridge made of slide glasses with spacers matching tissue thickness (*32*). After filling the gap of the cartridge with the mELAST solution, the tissues were completely submerged in the solution. The top slide glass was covered with an adhesive tape and mechanically secured with binder clips. The cartridge was then placed in a 50 ml conical tube, which was purged with 12 psi of Nitrogen gas for 1 minute before the lid was tightly closed. The tube was placed horizontally on a tube rack at room temperature and left to gel for 6-7 hours.

Hydrogelation of mouse organs was carried out using a 15 ml conical tube. This tube was cut with the cap and filled with 2 ml of the mELAST solution. The SHIELD-cleared and mELAST solution-equilibrated mouse organs were then submerged in the solution. Subsequently, the tube was placed inside a 50 ml conical tube, followed by the same nitrogen gas purging and gelling processes.

Human brain slabs were hybridized with hydrogel by *in-situ* polymerization in an air-tight stainless gelling chamber (fig. S7D). SHIELD-cleared human brain slabs, equilibrated with the mELAST solution, were placed on a glass plate (Borosilicate Glass Sheet, 17.78 cm x 10.16 cm x 0.3175 cm, 8476K49, McMaster) inside a plastic container filled with the mELAST solution. Care was taken to avoid trapping air bubbles. A top glass plate (identical to the bottom plate) was then placed over the tissue in the mELAST solution, and the two glass plates were mechanically secured on each side with two stainless steel binder clips. The glass plates with the tissues were transferred to the gelling chamber (fig. S7D). The chamber was covered with a transparent polycarbonate plate, tightly sealed with sixteen screws, and purged with nitrogen gas at 10-12 psi for 10 minutes. The gelling process was conducted at room temperature for 6-7 hours.

After the *in-situ* polymerization with tissue, the mELAST tissue-hydrogels were trimmed from the gel formed inside the cartridges or between the glass plates. To remove the unreacted thermal initiator and monomers, the tissue-hydrogels were first washed in the SDS clearing solution at room temperature for 24 hours and then at 45°C for another 24 hours, with three times solution refreshments. Protein denaturation in the mELAST tissue-hydrogels was achieved by incubating them in pre-heated SDS clearing solution at 95°C for 30 minutes, followed by hydrolysis at 80°C for 6 hours (*31*, *35*, *85*). After the hydrolysis step, the tissue-hydrogels underwent post-clearing with the SDS clearing solution, using a tissue pump that repeatedly compressed the tissue to 1/4 of its thickness overnight. Following these processes, the tissue-hydrogels were pumping-washed with 1% 1XPBSTN for 5-6 hours, with several refreshments of the buffer solution. The tissue-hydrogels were then photobleached for 1-2 days at 4°C and stored in 1XPBSN at 4°C until use.

Photobleacher design and operation

The mELAST tissue-hydrogels were photobleached to enable high signal-to-noise ratio (SNR) imaging across a wide spectrum by removing autofluorescence (e.g., lipofuscin) commonly found in long-term PFA-fixed postmortem human brain tissues. The photobleacher includes a transparent acrylic chamber to contain samples and approximately 300 white LEDs (Fig. 4D). These LEDs emit strong light onto the top and bottom surfaces of the tissue samples. To prevent overheating from the LEDs, two sets of cooling fans were incorporated into the design. The mELAST tissue-hydrogels were placed in a zip bag filled with sufficient 1XPBSN, and this bag was then placed in the chamber of the photobleacher, which was situated in a cold room (4°C) for 1-2 days.

Tissue pump design and operation

We developed tissue pumps in three different sizes for compression-assisted immunostaining of mELAST tissue-hydrogels: small (30 mm x 30 mm), medium (85 mm x 85 mm) and large (150 mm x 150 mm) (Fig. 4G and fig. S14B). In this design, mechanical thinning was achieved through the relative linear movement between a pump head and a flat-bottomed container holding mELAST tissue-hydrogels and buffer solution. During each pumping cycle, two linear actuators (with maximum pushing force of 1.3 kN) raised the container upwards towards the stationary pump head until the tissue was compressed to a predetermined thickness, such as 0.5 mm for 2-3 mm thick tissue. The actuators maintained the compression for 40 seconds before returning to the starting position. An Arduino microcontroller was programmed to control the cyclic movement of the actuators fully automatically. Given the large lateral dimensions of mELAST-processed human brain tissue-hydrogels, it was crucial to maintain the plane of the container bottom's plane strictly parallel to the pump head plane for uniform compression staining throughout the entire tissue area. Therefore, we incorporated an extensively long linear movement guiding mechanism to control the tilting of the moving plane. To further ensure uniform compression, we designed a mechanism that redistributes the compressive force from the actuators to evenly distribute it at the bottom of the container. Our staining experiments using the pump demonstrated effectively uniform staining results. The machine also featured adjustable pumping and retracting speed, adjustable compression thickness, and a user-friendly interface. We characterized the tissue compression repeatability of the pump by measuring the displacement of the moving plane with a laser displacement sensor. The results showed that when the target compression thickness was set at 0.6 mm, the pump consistently achieved this target with approximately 99% repeatability (Fig. 4H).

SeTau647 conjugation with secondary antibody

The highly photostable SeTau647 dye was conjugated with Fc Specific Fab Fragments secondary antibodies for imaging mELAST-processed human brain tissue-hydrogels. 100 μg of Fab fragments underwent buffer exchange to 0.1 M sodium bicarbonate buffer using a Zeba desalting spin column (ThermoFisher, Cat. No. 89883, 7 kDa) to remove preservatives. Subsequently, SeTau647-NHS ester (Seta Biomedicals, K9-4149) was added to the antibody solution to achieve a final concentration of 0.5 mM. The reaction mixtures were then incubated for 15-16 hours in darkness at room temperature with gentle shaking. The mixtures were subsequently separated and purified via size-exclusion column chromatography (GE, Superdex 200 15-300 increase) in Akta pure 25L equipped with the fraction collector. The relevant fractions were concentrated using an Amicon mass cut-off spin column (Millipore Sigma-Aldrich, 30 kDa, 2 mL, Cat. No. UFC203024). The dye conjugation was analyzed using 4-15% SDS PAGE. The final fractions containing the desired conjugates were combined and their concentration was measured using the microBCA assay (Thermo). After determining the exact concentration, preservatives such as 40% glycerol, 0.02% sodium azide, and 0.1% w/v BSA were added to the antibody-oligo conjugates. The product was stored at 4°C until use.

Passive staining and imaging

Sectioned tissues were pre-incubated in 1XPBSTN with 5% normal donkey serum (for antibody validation in figs. S9 and S10) or the SWITCH staining solution (for other staining, 1XPBSNaCh with 5% NDS) at 37°C for 3 hours. For antibody validation (figs. S9 and S10) and synapse imaging (Fig. 6L), the tissues thicker than 1 mm were sliced into 200 μm thick sections using a Leica vibratome (Leica VT1200). The pre-incubated tissues were transferred to fresh staining buffer solution containing primary antibodies. After overnight staining with primary antibodies at 37°C, the tissues were washed with 1XPBSTN (for antibody validation) or 1XPBSNaCh (for other staining) for 3-4 hours, refreshing the solution three times. Secondary antibody staining (at a 1:3 molar ratio of primary to secondary antibodies) was performed overnight at 37°C in 1XPBSTN or 1XPBSNaCh. The tissues were again washed with 1XPBSTN or 1XPBSNaCh for 3-4 hours, with three times of solution refreshment. After staining and washing, SHIELD-cleared tissues were incubated in dPROTOS for 30 minutes for refractive index matching. The mELAST tissue-hydrogels were incubated in water for 4x physical expansion for 30 minutes for antibody validation (figs. S9-S10) and synapse imaging (Fig. 6L). The optically cleared tissues were imaged using a Leica confocal microscope or Andor Dragonfly high-speed confocal imaging system with the 20x/0.5NA, 25x/0.95NA or 63x/1.2NA water immersion objectives.

To achieve a higher signal-to-noise ratio chemical synapse imaging, brief fixation with formaldehyde was employed for synaptic target labeling. After primary staining, the samples were incubated in 4% formaldehyde in 1XPBS at room temperature for 1 hour with shaking for brief fixation. This was followed by a 1-hour wash in 50 mM Tris solution at room temperature with shaking and a 3-hour wash in 1XPBSTN with 5% NDS at room temperature with shaking. Finally, the samples were incubated with secondary antibodies in 1XPBSTN at 37°C overnight with shaking, followed by final washes in 1XPBSTN at 37°C for 4–6 hours, including at least three solution exchanges.

After expansion in DIW, the tissues were imaged using a Leica confocal microscope with a 63x/1.2NA water immersion objective.

For (i) MEGAtome validation (Fig. 2. L and M), (ii) intact slicing and imaging of coronal human brain hemisphere slabs and brain arrays (Figs. 3 to 5), and (iii) UNSLICE demonstration (Figs. 7-8), the tissues were sliced using MEGAtome for antibody labeling and imaging. The 1- and 4-mm thick SHIELD-cleared human brain slabs (both coronal and pons slabs) and 2 mm thick SHIELD-cleared mouse and marmoset brain arrays were stained with primary antibodies in 1XPBSTN with 5% normal donkey serum at 37°C for 2 days (1 mm thick), 4 days (2 mm thick), and 6 days (4 mm thick), followed by washing with 1XPBSNaCh for 1 day (1 mm thick), 2 days (2 mm thick), and 3 days (4 mm thick), refreshing the buffer twice daily. After primary staining, the tissues were stained with secondary antibodies and YOYO™-1 Iodide nuclei dye (Y3601, Invitrogen) in 1XPBSNaCh at 37°C for 2 days (1 mm thick), 4 days (2 mm thick), and 6 days (4 mm thick) and washed with 1XPBSTN for 1 day (1 mm thick), 2 days (2 mm thick), 3 days (4 mm thick) with two times buffer refreshment every day. For 4 mm thick tissues, 1% Triton-X was added 24 hours before the end of the primary and secondary staining as an ON step. The stained tissues were optically cleared with dPROTOS for 1 day (1 mm thick), 2 days (2 mm thick), or 3 days (4 mm thick). For refractive index (RI) matching with dPROTOS, the tissues were first incubated in a solution with a dPROTOS to water ratio of 1:2 for 1 day (2 mm thick) or 2 days (4 mm thick) at 37°C, followed by full dPROTOS incubation for 1 day at 37°C. The optically cleared tissue slabs were imaged using MegaSPIM with the 2x/0.1NA or 4x/0.2NA objectives. Details are summarized in table S4.

<u>Multi-round SWITCH-pumping staining and imaging</u>

Before SWITCH-pumping staining, mELAST tissue-hydrogels were pre-incubated in the SWITCH staining solution (0.2XPBSNaCh). The tissue pumps were used for staining to accelerate probes delivery deep into the tissue-hydrogel by repeated thinning (from 2-2.5 mm to 0.4-0.6 mm) and solution refreshment (40 seconds for compression and 2-3 seconds for refreshing). After 16-18 hours of pumping-staining of the tissue-hydrogels with primary antibodies in the SWITCH staining buffer, 20% Triton-X was added to the staining buffer at a 3:1 volumetric ratio, resulting in a final Triton-X concentration of approximately 5%. After a total 24 hours of primary antibody staining, the tissue-hydrogels were washed with 0.2XPBSNaCh for 12-16 hours, refreshing the buffer three times. The tissue-hydrogels were then stained with secondary antibodies and nuclei dyes (YOYO™-1) in 0.2XPBSNaCh for 16-18 hours with cyclic compression. Following secondary staining, the tissue-hydrogels were washed with 1XPBSTN for 3-4 hours using the tissue pumps. The stained tissue-hydrogels were optically cleared using dPROTOS and imaged using MegaSPIM with the 2x/0.1NA and/or 16.7x/0.4NA objectives. After each imaging round, the mELAST tissue-hydrogels were destained in preparation for the next round. For destaining, first, the tissue-hydrogels were pre-incubated in the SDS clearing solution overnight at 45°C. The bound antibodies and probes were eluted by applying thermal stress at 80°C for 1 hour, followed by pumping-destaining at room temperature for 3-5 hours with 2-3 times solution refreshment to wash out the unbound antibodies and probes. The destained tissue-hydrogels were then washed with 1XPBSTN using a tissue pump for 3-5 hours at room temperature to remove SDS molecules inside the tissue-hydrogels. Prior to the next staining round, the buffer was exchanged to 0.2XPBSNaCh with 5% NDS using a tissue pump for 2-3 hours at room temperature. The primary staining for the next round then commenced. Detailed information on each antibody staining and imaging procedure is listed in table S4. The 3D image volumes and supplementary movies showing cells from the multiplexed tissue-hydrogels were created using Imaris software.

MegaSPIM design and operation

Multicolor-Inverted Axially-Swept Lightsheet Fluorescence Microscopes (named MegaSPIM) was designed to accommodate tissues with large xy dimensions and capture multicolor data with simultaneous two-color imaging. The light source of IASLSFM is a multicolor laser combiner with a fiber optic output that delivers continuous wavelength light into the illumination arm of the system. The microscope was equipped with 405 nm, 488 nm, 561 nm, and 647 nm continuous wave laser sources for acquisitions with up to four colors. A cylindrical lens shapes the beam into a sheet, and an electrically tunable lens is used to sweep the light along the propagation axis, such that the lightsheet waist is always in-focus and acquired by the rolling shutter of the camera. This created a near-perfectly evenly illuminated optical section with consistent axial resolution and allowed for high-speed line-scanning detection. Illumination and detection on these systems was achieved through pairs of objectives set up orthogonally to one another and at a 45-degree angle to the sample. These objectives can be swapped out depending on resolution and speed requirements. For full human brain slab imaging, a 2x 0.1NA objective was used to acquire cellular resolution data. These are long working distance objectives with custom dipping caps to acquire image data through up to ~12 mm of depth. During multicolor imaging, detected light from the fluorescent probes is captured and sent through a dichroic mirror which separates the longer two wavelengths from the shorter two wavelengths and sends each to a dedicated camera. Color pairing is logically determined in software so that imaging is done simultaneously for 405 nm and 561 nm as a pair, and 488 nm and 647 nm as a pair. To account for chromatic shifts between different colors detected at the same focal plane, each detection arm has its own tube lens mounted to an actuator-controlled translation stage which allows for independent focusing of each color's image into its respective camera. The tissue slabs were scanned in long strips along the X-axis of the stage during acquisition. Each microscope's xy stage was a 200 mm x 200 mm MS-8000 from ASI with a "precision scan" X-axis, which allows very low velocity ripple during these long acquisition movements. Synchronization between the lasers, camera, tunable lens, and stage were controlled through ASI's Tiger controller and custom LabVIEW software. We also have a custom-built 400 mm x 200 mm xy stage assembly from ASI to allow imaging of the largest, most expanded human brain slabs which are too big for the MS8000. Calculated selection of the step size in X allowed us to acquire a near-isotropic voxel size. The nature of the 45-degree illumination and detection relative to the sample and stage resulted in a coupling of the X and Z axes. Accommodating this relationship with a careful x-step-size choice enabled high axial resolution as well as clean processing and visualization of the data. Voxel columns could be simply be re-shuffled to convert from the 45-degree-tilted coordinate space into typical xyz coordinate space.

Sample mounting for MegaSPIM imaging

RI-matched tissues were mounted on a custom-machined plastic holder made from black polycarbonate to minimize light scattering. Velcro strips, adhered to the plastic holder, were used to secure the agarose and provide stability during stage movement. A 1% agarose solution, prepared in the RI matching solution (dPROTOS), was poured onto the holder, which was equipped with the side walls made of Blu-Tack LLC Reusable Adhesive. The RI-matched tissues were then placed on the gel. Additional agarose/RI matching solution was poured over the RI-matched tissues to fix them on the holder. The entire plastic holder was submerged in a chamber filled with the RI matching solution (dPROTOS) and held at the bottom with four magnets. A layer of silicon oil on top of the immersion media prevented water evaporation from the solution. The mounted tissues were imaged using MegaSPIM.

MegaSPIM image processing pipeline

We built a custom pipeline that reconstructs the images from our MegaSPIM into three-dimensional volumes suitable for visualization and analysis. The MegaSPIM camera microscope objective orientation is at a 45-degree angle to the stage motion so that a single planar image is aligned along both the X and Z axes of stage motion. Planar images were acquired sequentially in synchrony with X stage motion to form a stack, then the stage was moved in Y to acquire subsequent stacks with approximately 200 voxels of overlap between each stack. The steps of the pipeline are:

- Conversion of the proprietary and uncompressed Hamamatsu DCIMG file format to lossy compressed JPEG 2000 images.
- Destriping and illumination correction of the planar images.
- Reconstruction of the planar image pixels into an orthogonal X, Y, Z voxel space.
- Alignment of adjacent stacks
- Either stitching of the stacks to create a unitary volume or recording of the alignment between stacks according to the BIDS specification (*86*).

The data was first converted from its raw form to JPEG 2000 planes. Lossy compression was used, with a maximum allowed loss of information of 80 db. This resulted in a typical reduction in storage space required for the image data of 7x. Each plane was then destriped according to our previously-published protocol (*87*) to remove the shadows that are cast in the lightsheet downstream of an occluding object. Briefly, the image was decomposed into three sets of wavelets in the direction parallel to, perpendicular to and at a diagonal to the light path, the wavelets parallel to the light sheet path are de-emphasized at a set of frequencies and the image was reconstructed to yield a result with the stripes largely removed. Image correction was then done by dividing the image intensity at each pixel by an illumination function at the corresponding pixel (*88*). The illumination function was created by scanning a uniformly-dyed field, smoothing the result. The stack volume was then reconstructed by creating a grid in X, Y and Z and interpolating each grid point's position to the corresponding plane and pixel in the corrected images. Adjacent stacks were scanned with overlap in order to calculate any offsets of one stack with respect to the other caused by deviations from expected due to slight stage or optics tilt or inaccuracies in the stage. The alignment step used a blob detector (blur with a difference of Gaussians followed by finding local maxima) to find punctate objects in the overlapped region. A subset of these was selected and for each, the Pearson correlation coefficient was calculated between the images in each overlapping stack within a 51-voxel window after blurring the images. The Pearson correlation coefficient was calculated, shifting the image by one in each direction and the direction with the highest coefficient was chosen and the process was repeated centering around the chosen voxel from the previous iteration. The process was repeated until no immediately surrounding voxel has a coefficient higher than the central voxel or until a certain number of iterations have been executed. The offset in X, Y and Z was kept if the coefficient is higher than a cutoff value (typically .95) and otherwise discarded. Finally, the calculated offset of one stack with respect to the other was computed by taking a median weighted by the correlation coefficient. The alignment result could be applied in one of two ways. The alignment could be used to reconstruct a unitary volume containing all the stacks, displaced by the recorded offsets or the offsets could be recorded in the ChunkTransformMatrix field of BIDS sidecar files. The volume could be reconstructed on the fly as specified by the BIDS specification. All software described above is open-source and available in the "spimstitch" package: https://github.com/chunglabmit/spimstitch. The software was written in Python, tested on Linux and easily installed using PIP.

Image upload to DANDI

Post alignment, the data were locally organized into a BIDS (*89*) dataset using the BIDS Microscopy specification (*90*). This arranges the dataset per participant. Each acquisition session is stored in a separate folder containing a photo of the slab and the different stains acquired on each slab during that session. In a few cases (LIST sample numbers), the data from the same slab were acquired in multiple sessions. The 3D images corresponding to each slab were stored as a multi-resolution pyramid using the OME-ZARR format (*91*), including the alignment information. Other associated metadata were stored in sidecar JSON files. All data was uploaded to the DANDI data archive (RRID:SCR_017571; (*92*) using the DANDI command line tools (https://pypi.org/project/dandi/). The DANDI archive stores data on AWS S3 and combines it with the OME-ZARR format, allowing direct visualization of the stitched slabs on the archive using tools such as Neuroglancer. A dashboard for this dataset is available at https://biccn.github.io/Quarterly_Submission_Receipts/000108-dashboard.html when uploads are finalized.

Co-registration of multi-round images

Custom software was written in Python to perform the alignment of a first round of imaging (the fixed round) to subsequent rounds (the moving rounds). Typically, the volume images were rotated, displaced and stretched with respect to each other. The software is in the form of a pipeline and GUI to run the pipeline. The output of the pipeline is a bidirectional warping field that can be used to translate coordinates from the space of the fixed round to that of the moving round and vice-versa. The GUI also contains tools to perform image warping of the moving volume to the space of the fixed volume. The pipeline’s starting point is a pair of imaged volumes from the fixed and moving space, typically of the sample stained with a nuclear stain. The pipeline starts with a rough manual alignment step followed by steps of finding pairs of points in each of the volumes that match, filtering out bad matches and computing a warping function. The last three steps are repeated until the warping function converges on a solution.

The user must perform a rough manual alignment as a starting point for the automatic alignment. The user selects pairs of corresponding points in downsampled images of both volumes. A minimum of four non-coplanar points must be selected to define the rotation and translation of the moving volume into the fixed volume space. Selecting more points allows the algorithm to more quickly converge on the final alignment. The point pairs are used to create a thin-plate spline transformation function (*93*) to convert coordinates in the moving frame of reference into the fixed frame of reference and vice-versa. The thin-plate spline function is then approximated by a grid of displacements.

The pair finding step of the pipeline takes a set of points of interest as one of its inputs. These points should ideally have clear monotonically decreasing intensities at increasing distances from the point to make the pair finding step's gradient descent work accurately. Nuclei centers work well for this. The centers are found by blurring the image using a difference of Gaussians and finding local maxima whose intensities are above a preset threshold and are at least a preset distance from any maxima of greater intensity. The pair finding step takes as inputs the moving and fixed images, the results of point finding and either the warping function from the manual alignment or the warping function calculated during the previous round of pair finding and filtering. A grid is laid over the fixed volume and the nearest nucleus center to each grid point is found. Grid points that do not have a nucleus center within a parameterized distance are discarded. A patch of the moving image around each of these points is warped into the space of the fixed image and a patch of the fixed image is also taken. Both patches are blurred with a Gaussian and the Pearson correlation coefficient of the two patches is taken as well as the correlation coefficient of the two patches, displaced by one voxel in each of 26 possible directions to form a 3x3x3 array. The location of the center of the moving image is updated to the coordinate with the highest correlation coefficient and the process is repeated until the highest correlation in the array is at the array center. This procedure results in a set of pairs that are purportedly at the same location in the fixed and moving volumes.

The pair filtering step takes the warping function and the set of pairs from the pair finding step as inputs. The fixed points are warped into the moving space and an affine transform mapping the moving points into the warped fixed-point space is calculated using RANSAC. This affine transform is used as a secondary warping that is applied to the already-transformed moving points. A vector is calculated between each warped fixed point and its transformed moving point partner. The cosine distance between this vector and the vectors of each of the point's two nearest neighbors is computed and the average of these two cosine distances is calculated to determine whether the displacements of the point of interest are similar to the two nearest neighbors. The resulting value is between 0 and 2, with 0 indicating that the displacements are in the same direction and 2 indicating that they are in opposite directions. If this mean cosine distance is above a parameterized threshold, the pair is discarded for not being in agreement with its neighbors.

The filtered pairs of coordinates are used to make a warping function using the same procedure used to calculate the manual alignment warping function. This new warping function takes the place of the previous warping function and pair finding and filtering is performed again.

The last step in the pipeline is a procedure that applies the warping function to each channel of the moving image to transform that channel into the fixed volume's space. The GUI is open-source at https://github.com/chunglabmit/multiround-alignment-ui. The software contains an installation script for Anaconda which will create an environment for the software stack on any Linux machine. Installation and usage instructions and a video tutorial can be accessed from the repository's README page.

We used this pipeline to co-register the multi-round images from the mELAST human brain tissues at single cell resolution (Fig. 4J and Fig. 5C-D). This approach is capable of accurately transposing many rounds of imaging into the same reference frame at single-cell resolution (fig. S14 and S17), which enabled quantification and analysis of cell subtypes based on many cell type markers, regardless of the staining patterns in most of the tissue types (Fig. 5). The same transformation matrix computed for the nuclei was applied to other channels, enabling multiplexing images (fig. S14B and S17B). The repeated experimental and computational processes generated a multiplexed proteomic map of the human brain orbitofrontal cortex (movies S4-5), revealing the 12-plexed 3D pattern of 11 major neuronal and non-neuronal subtypes and nuclei.

NeuroTrALE for brain region segmentation

Preprocessed and illumination corrected images were converted to precomputed volumes, then uploaded to MIT Supercloud (https://supercloud.mit.edu/), followed by a 4x downsampling for complete and smooth display in NeuroTrALE, a web-based data review and editing tool derived from Neuroglancer. Human brain slabs were segmented based on Allen Brain Institute Adult Human Brain Atlas, modified Brodmann (https://atlas.brain-map.org/atlas?atlas=265297126) and gyral (https://atlas.brain-map.org/atlas?atlas=138322605) (*13*). Corresponding coronal hemisphere slab and reference atlas were identified by comparing their shape and relative position. Regional segmentation was performed in NeuroTrALE using custom-developed polygon annotation functions. Polygons were drawn from scratch surrounding each individual brain region matching the atlas, and individual polygon points and edges could be moved, deleted, and subdivided to facilitate efficient tuning of annotations. Every $10^{th}$ z-slice was manually annotated within a region of interest, and post-processing steps applied linear interpolation to fill in the gaps. The resulting segmentation slices were merged into a 3D volumetric mask to be used for cell density counting within each segmented region.

Improving image quality

A series of image processing algorithms was applied to enhance signal-to-noise (SNR) ratio. After thresholding intensity ranges of the images, a skeletonization algorithm (*76*) was applied to detect non-specific signals such as blood vessels. The non-specific signals were then removed by a custom ImageJ script. The resulting images were then binarized to generate masks where 0 and 1 represent background and foreground signals respectively. Finally, the masks were multiplied to the original images.

pNFP+ fiber orientation analysis

OrientationJ (http://bigwww.epfl.ch/demo/orientation/) (*94*), an ImageJ plugin was used to compute the preferred angular orientation of each pixel in both pNFP-stained control and AD tissue images using finite difference gradients. The coherency (representing how similar a pixel's orientation is to its neighborhood) and energy (representing the gradient magnitude) of each pixel were also computed, and a threshold of 0.1 for coherency and 0.2 for energy were set to filter out spurious background pixels from being included. The pial surface orientation was estimated after manual cortical layer segmentation by averaging the orientation vectors that resulted from pairwise connection of adjacent sampled points along each cortical layer. The final orientation angle for each pixel was then computed by subtracting this pial surface orientation angle from the angle calculated using OrientationJ.

Cell detection and density analysis

The NeuN+ cells from both slabs were automatically detected using a 3D cell phenotyping algorithm (*38*). The Generalist 3D Neuronal Cell Classification algorithm detected and classified cells in 2 steps (CR, PV, Iba1, NPY, SST). First, all the cells in the volume were detected by nuclei staining segmentation using StarFinity (*95*). The quantitative information of the staining of other markers was extracted by the algorithm based on the nuclei segmentation results. Semi-automatic thresholding was applied to the quantitative measurements to perform the phenotypic-information-based categorization. After computational detection, obvious false positive cells were manually counted and excluded from cell density calculations. Detection accuracy (%) was calculated as 100*(manual cell count - undetected cell count - false positive count)/manual count.

Synapse detection and density analysis

A custom-built automated morphology-based cell detection framework was used for synapse detection. A curvature- and intensity-based detection algorithm identifies synapses in high-resolution images and produce center coordinates (3D). Detected synapses were counted per ROI (Region of interest) and normalized by the area of the regions ($mm^3$) to compute density. The final density per region was computed by averaging density scores of regions in each experimental condition. At least 10 candidate regions were used for each condition, and the minimum volume size per region was 100 $mm^3$.

Inter-slab registration (UNSLICE)

**Image pre-processing and illumination correction.** Images were converted to the easily chunked and parallelizable compressed zarr format, and then illumination corrected to even out the background artifacts that may occur near the surface of during lectin staining. A user-defined background intensity percentile of the image histogram in a local neighborhood around each pixel was subtracted from the pixel to correct for uneven illumination.

**Tissue surface detection and flattening.** For tissue surface detection, images were first significantly downsampled to mitigate computational expense. The tissue was then segmented from the background by applying graph cut segmentation (*96*) on the image.

Prior to graph cut segmentation, the image intensity histogram was clipped in order to reduce background noise (minimum clipping) and reduce variance of positive signal intensities (maximum clipping), since the image channel contains both autofluorescence and immunostaining signal which far exceeds the desired autofluorescence signal. After graph cut segmentation, morphological opening with a spherical element was performed to fill any potential holes in the tissue mask. Parameters for this were set manually after quickly sampling various parameter sets on subsets of the image and viewing the results. The specific parameters were (1) minimum clipping threshold, (2) maximum clipping intensity, (3) morphological opening size, and (4) graph cut smoothing parameter. In our experience, (1) and (2) are the only parameters requiring user tuning.

The top and bottom surfaces of the tissue were detected from the tissue segmentation by finding the uppermost or bottommost segmented pixel at each ($x$, $y$). To flatten the tissue, a point cloud grid was sampled from its two surfaces and triangular meshes were created from each surface point cloud. Based on the quality of surface detection, the user can opt to perform between zero to three outlier removal methods to clean up the generated point cloud: (1) gradient outlier removal, (2) statistical outlier removal, or (3) manual concave hull outlier removal. Gradient outlier removal was performed by estimating the gradient at each surface point by fitting a tangent plane locally at that point, and then removing all surface points with a gradient magnitude steeper than a user-defined threshold. Statistical outlier removal was performed by computing the average distance to each point's $k$ nearest neighbors, and removing all points that are 4 standard deviations above the average distance of all points. Manual concave hull outlier removal was performed by using the Neuroglancer point annotation tool to label the 2D concave hull of the tissue surface, generating a 2D mask of the surface by filling the concave hull, and masking the surface point cloud to eliminate points that are outside of the surface in 2D. Finally, after the filtered point cloud of surface points are generated, the target 2D projection for the points in the flattened reference frame for each surface are determined using Boundary First Flattening [2]. Thin point splines (TPS) were then used to interpolate the transformation vectors (from unflattened reference frame to flattened reference frame) for a regular grid over the entire unflattened image.

**Initial approximate slab alignment.** To perform initial slab alignment, matching manual anchor points were identified in the two adjacent slabs using the point annotation feature in Neuroglancer. These are typically large, easily identifiable features such as large arteries or vessels, and ideally span the entire cut surface, particularly near the edges of the tissue. An initial rigid transformation to align these anchor points was automatically calculated by minimizing the least squares error, followed by a nonrigid transformation computed using TPS with high regularization to ensure a smooth transformation.

**Automated tracing of vasculature.** Images were first filtered using a custom GPU implementation of the Optimally-Oriented Flux (OOF) filter (*97*), which for each pixel maximizes the outward image gradient flux flow across a sphere of given scale (or radius), as given by the expression below:

$$f(x; r, \hat{\rho}) = \int_{\partial S_r} ((v(x+h) \cdot \hat{\rho})\hat{\rho}) \cdot \hat{n} dA$$

*r* is the scale, or radius of the sphere, $v(x+h)$ is the image gradient at the sphere surface centered at a given pixel location *x*, $\hat{n}$ is the normal vector to the spherical surface, and $\hat{\rho}$ is the direction of the image gradient being projected onto the sphere. In order to generate a higher response at the vessel centerline, an Oriented Flux Asymmetry (OFA) measure (as defined below) that gives a high response at curvilinear structure edges is subtracted from the OOF measure to accentuate the vascular centerlines (*98*).

$$s(x; r, \hat{\rho}) = \int_{\partial S_r} v(x+h) \cdot \hat{\rho} dA$$

The OOF and OFA measures can be efficiently computed by converting the integrands into a set of linear filters, and then applying the Fast Fourier Transform on the integral. For OOF, this returns $\Lambda(x; r)$, the strongest eigenvalue response associated with the maximal flux direction and a certain radius, while for OFA, this returns $Q(x; r)$, the magnitude of the projected image gradient on the vessel cross-sectional plane (where the cross section eigenvectors are determined from the OOF computation).

$$\Lambda(x; r) = f(x; r\ , \widehat{e_i}(x, r)) = \lambda_i(x; r)$$

$$Q(x; r) = \sqrt{s^2(x; r, \widehat{e_1}) + s^2(x; r, \widehat{e_2})}$$

The filter should give the strongest response at the center of vessels and weak at the edges. Since OOF gives strong responses inside vessels and OFA gives strong responses at vessel edges, the overall filter response is given by:

$$M(x) = \big(0, \Lambda(x; r) - Q(x; r)\big)$$

Segmentation of the vessels was done by performing intensity thresholding or graph-cut segmentation in parallel on different chunks of the overall image. The specific intensity threshold for a given image was chosen manually based on user parameter tuning performed on sampled regions of the image.

**Feature matching and inter-slab registration.** As described in detail earlier, GPU-accelerated Optimally-Oriented Flux (OOF) followed by global intensity threshold segmentation were implemented to generate a mask of the vasculature, which was followed by parallelized medial-axis thinning on the vasculature mask to extract the vessel centerlines and transform the image into a graph of nodes, edges, and endpoints. The locations of endpoints lying near the previously determined surfaces were extracted and used as features for image registration.

A custom feature descriptor was used to correctly match corresponding vasculature endpoints in both slabs. Each endpoint feature descriptor consists of the relative (*x,y,z*) coordinates of its *k* (user-defined parameter) 3-D nearest neighbors. The search for potential feature correspondences for a given key point was localized to within a certain radius *r* (user-defined). Candidate key point matches within the radius were determined by evaluating the distance between the feature descriptors; however, in order to be more robust to imperfect vessel endpoint detection, the descriptor similarity was computed as the shortest distance between any $n \leq k$ (a user-defined parameter) spatial nearest neighbors using the Hungarian algorithm (*99*). For example, for $k = 6, n = 3$:

$$f_i = [x_{i1}\ y_{i1} z_{i1} x_{i2}\ y_{i2}\ z_{i2} x_{i3}\ y_{i3}\ z_{i3} x_{i4}\ y_{i4} z_{i4}\ x_{i5}\ y_{i5} z_{i5}\ x_{i6}\ y_{i6} z_{i6}] DS(f_1, f_2)$$
$$= \sum_{k \in S} \sum_{j \in S} \left|(x, y, z)_{1j} - (x, y, z)_{2k}\right|^2$$
$$S = \{(1,2,3), (1,2,4), (1,2,5), \ldots (3,5,6), (4,5,6)\}$$
where $f_i$ represents the keypoint descriptor vector for a point $i$, $(x_{ij}\ y_{ij} z_{ij})$ is the $j$th nearest neighbor 2D location to the point $i$, $DS(f_1, f_2)$ is the descriptor similarity between point 1 in one image and point 2 in the other image, and $S$ is the set of all $P(6,3)$ nearest neighbor permutations over which we compute the descriptor similarities. A good correspondence between a pair of keypoints was determined by computing the ratio of the best and second-best descriptor similarities and thresholding at 0.85:
$$\frac{DS(f_1, f_{closest})}{DS(f_1, f_{2nd-closest})} \leq 0.85$$
To further ensure a good correspondence, matching key points was done in the forward direction and in the reverse direction such that the correspondence satisfies the above inequality in both images.

Random sample consensus (RANSAC) with an affine model is applied locally in a radius about each detected matching point to filter out outlier feature correspondences, and thin plate splines interpolated from the keypoint correspondences were used to compute the nonrigid deformation from the moving to fixed surface. Point correspondences on the other (non-cut) surface of the tissue were defined as static to ensure good interpolation of the tissue volume, and a grid of static point correspondences at the z=0 optical section were also defined to eliminate non-physical TPS extrapolation at the cut surface. This is because the small number of points in the z-direction makes the transformation especially sensitive to small variations in z of the surface keypoint correspondences (both manual and automatically detected). To make the deformation computationally scalable, the TPS were computed exactly for a regular grid and approximated by interpolating the resulting deformation field with GPU-accelerated grid interpolation.

Inter-slab registration accuracy can be fine-tuned by manually annotating more matching endpoints in the newly registered image, followed by outlier removal via RANSAC, and finally defining another TPS transformation. This can be done iteratively to the desired level of accuracy.

**Inter-slab registration using multi-channel information.** The previous deformation field defined with vasculature features can be applied to warp other image channels. The above procedure for registering images can be applied to other channels to achieve finer resolution warping if using axon endpoint features. Specifically, axon endpoints can be detected automatically and/or manually and a new TPS transformation can be defined to further fine-tune the registration.

<u>Automated axon tracing</u>

**Network structure.** The U-Net architecture consists of convolutional layers interspersed with downsampling to form sequentially lower-resolution modules in an encoder path, followed by upsampling to restore the original resolution in a decoder path, with lateral skip connections between same resolution encoder and decoder modules. We used 3×3×3 convolutional layers followed by group normalization and exponential linear units. Summation joining was chosen for skip connections with an additional skip connection between the first and last convolutional layer in each module. We also used strided 2×2×2 max-pooling for downsampling, and strided transpose convolutions with max-pooling for upsampling. In our baseline approach, we trained a residual 3D U-Net with 4 resolution blocks to perform voxel-wise classification using binary cross entropy loss between the output and training images. The resulting segmentation was then binarized using a threshold of 0.5, and skeletonized using morphological thinning to obtain a single voxel wide centerline. To improve upon the baseline, we replaced binary cross entropy with an alternative loss function, centerline-Dice (clDice), which incorporates parameter-free skeletonization directly into the training procedure.

**Training details**. Each network was trained via stochastic gradient descent using the ADAM optimizer with an initial learning rate of $1\times10^{-4}$ and weight decay of $1\times10^{-3}$. For training, mini batches of 16 samples were randomly cropped from the input dataset, and augmented using random grayscale perturbations, random 90-degree X-Y rotations, and random flipping. For inference, samples were cropped using a sliding window with 50% overlap and blended using a 3D Hann window to minimize segmentation errors along the edge of the receptive field. We also incorporated test-time augmentation, by averaging predictions across 16 transformations for each input sample (unique combinations of X-Y 90-degree rotations and flipping along each axis). Test time augmentation has been shown to improve segmentation performance at the expense of a linear increase in inference time. For each model, training proceeded until convergence or until no improvement in validation loss was observed over a period of 100 epochs.

<u>pTau fiber orientation analysis</u>

Using the resulting graph of pTau fibers generated by the axon tracing algorithm, the orientation vector of each fiber was computed by subtracting one endpoint coordinate from the other. The azimuthal (θ) and polar (φ) angles were computed by transposing the Cartesian vectors (x,y,z) into spherical coordinates (r,φ,θ). Since the projection directionality of the pTau+ axons are not known (due to not knowing the location of each axon's cell body), we reflect the orientations of all fibers with $-90 < \theta < 0$ about the x, y, and z axes to constrain $0 < \theta < 90$. The orientations are then binned based on the (φ,θ) angles to generate the orientation histograms.

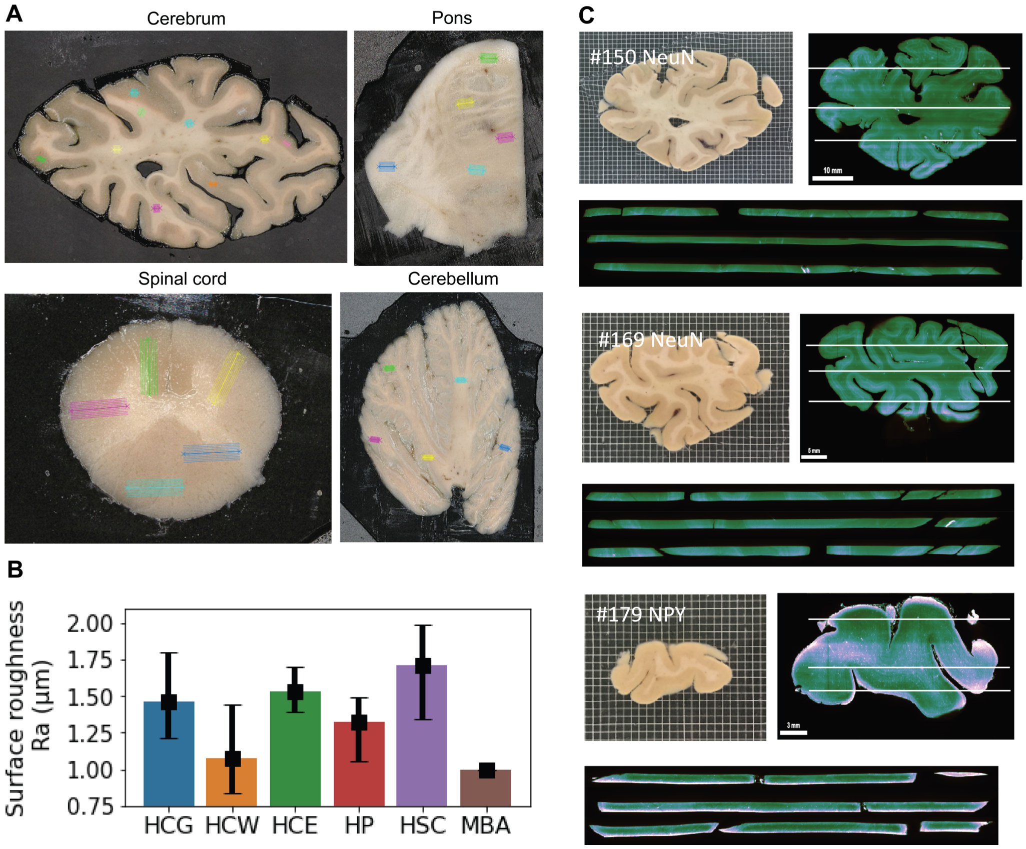


**Fig. S1.**

**MEGAtome slicing of the human brain.** (**A**) Slicing different regions of the human brain; cerebrum, pons, spinal cord, and cerebellum. (**B**) Measurement of the roughness of the sliced surfaces of different regions of human brain tissues. HCG: Human cerebral grey matter, HCW: human cerebral white matter, HCE: human cerebellum, HP: human pons, HSC: human spinal cord, MBA: mouse brain array (**C**) xz-plan view of light sheet microscopy images from the MEGAtome sliced and NeuN-stained human brain coronal slabs.

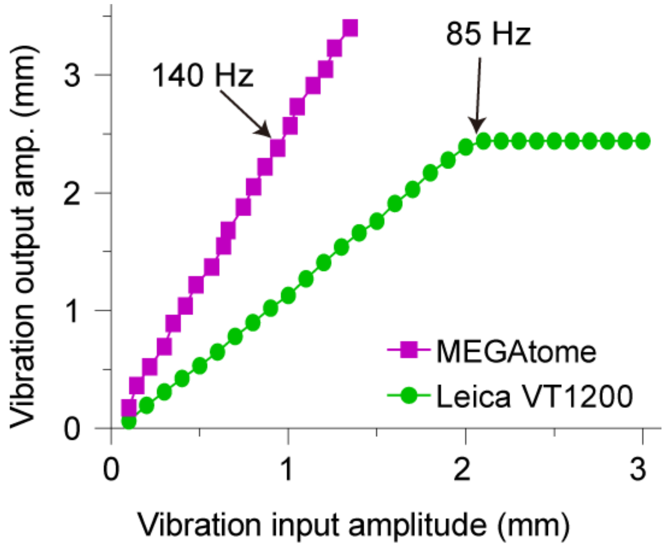


**Fig. S2.**

**Measurements of blade vibration output versus vibration input of Leica VT1200 vibratome and MEGAtome.**

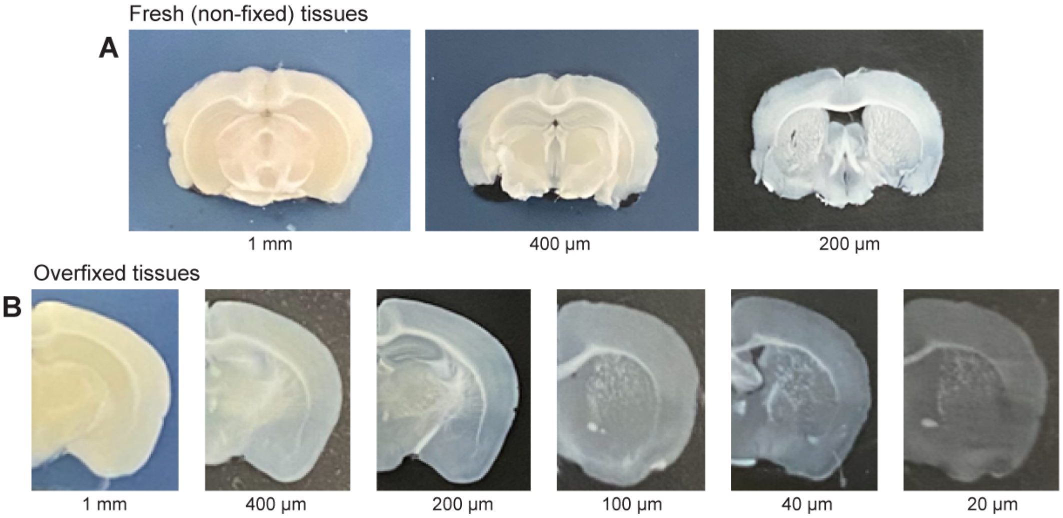


**Fig. S3.**

**MegaSPIM allows slicing of (A) fresh (non-fixed) tissues and (B) overfixed tissues.**

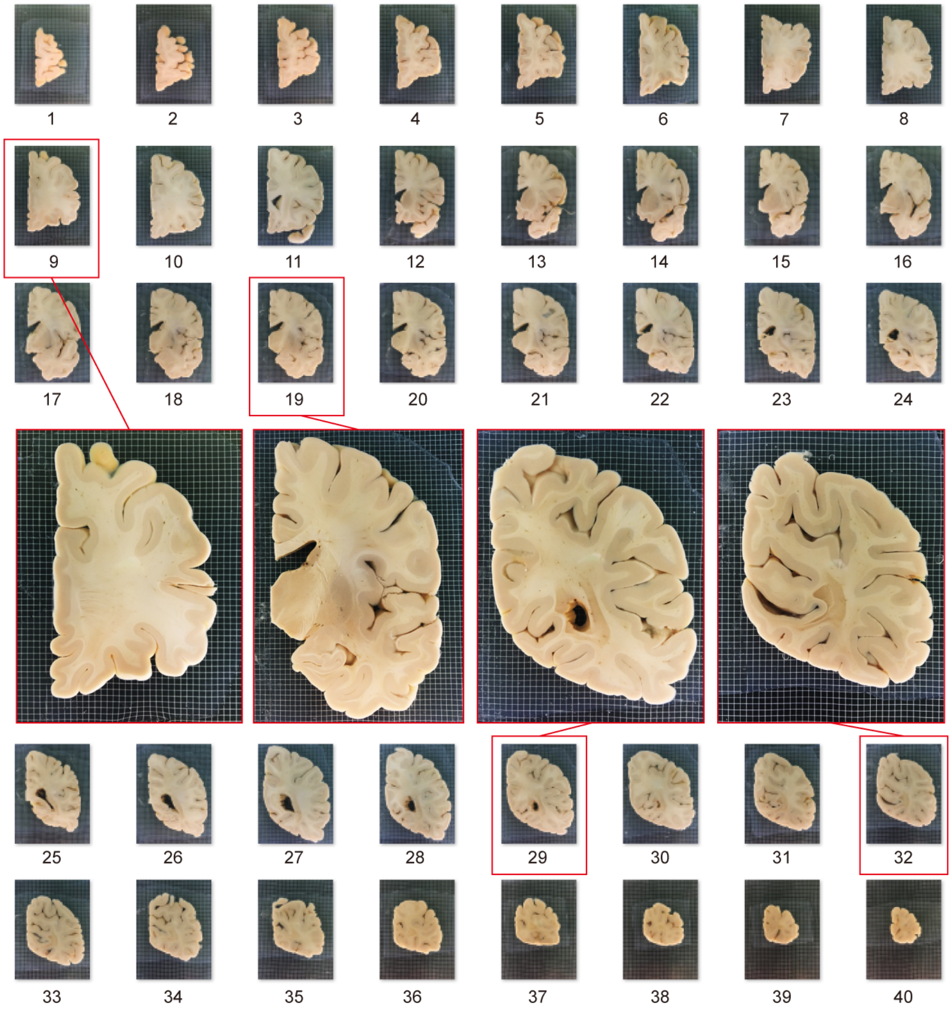


**Fig. S4.**

**A banked human brain hemisphere was embedded in gel and then sliced by MEGAtome into 40 consecutive 4 mm-thick uniform slices that kept complete tissue from the hemisphere.**

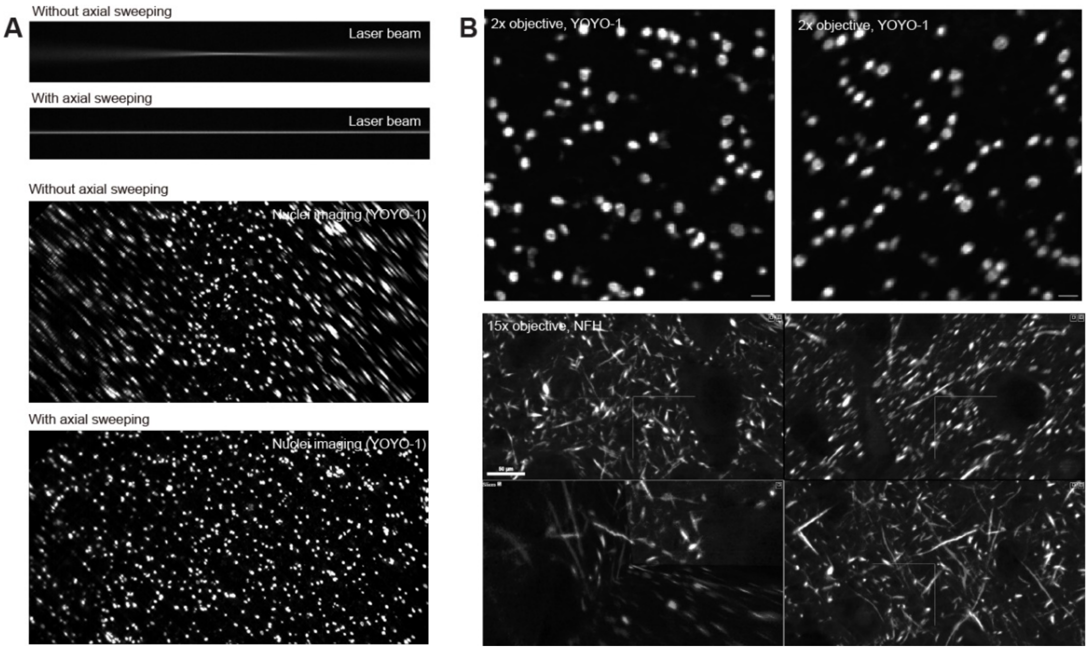


**Fig. S5.**

**MegaSPIM for multi-scale imaging of large-scale tissues.** (**A**) Images of the laser beam (top) and nuclei (bottom) without and with axial sweeping engaged across the field of view. (**B**) Images acquired from mELAST human brain tissues using the 2x (top, YOYO-1 stained nuclei) and 15x (bottom, NFH+ fibers) objectives.

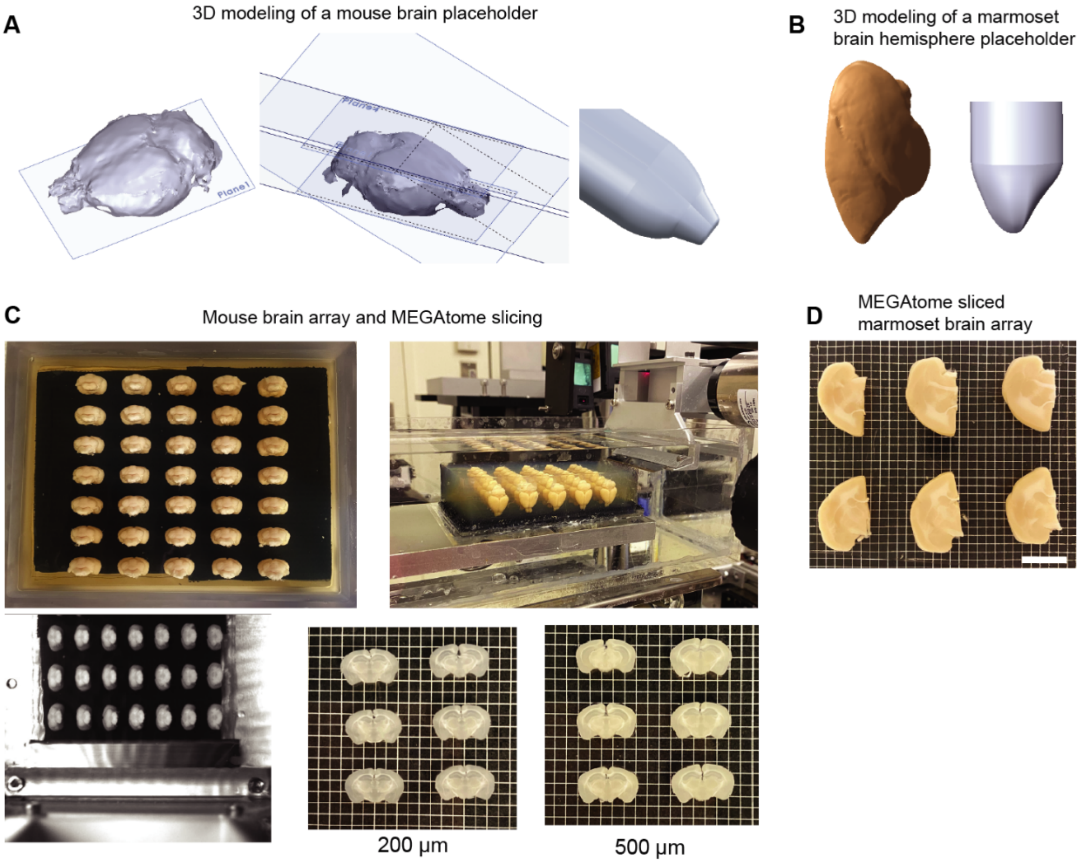


**Fig. S6.**

**3D modeling of the brains and MEGAtome-slicing of the brain arrays.** 3D rendering and modeling of the whole mouse brain (**A**) and marmoset brain hemispheres (**B**). (**C**) MEGAtome slicing of the mouse brain array. (**D**) MEGAtome-sliced marmoset brain array. One grid size: 2 mm x 2 mm.

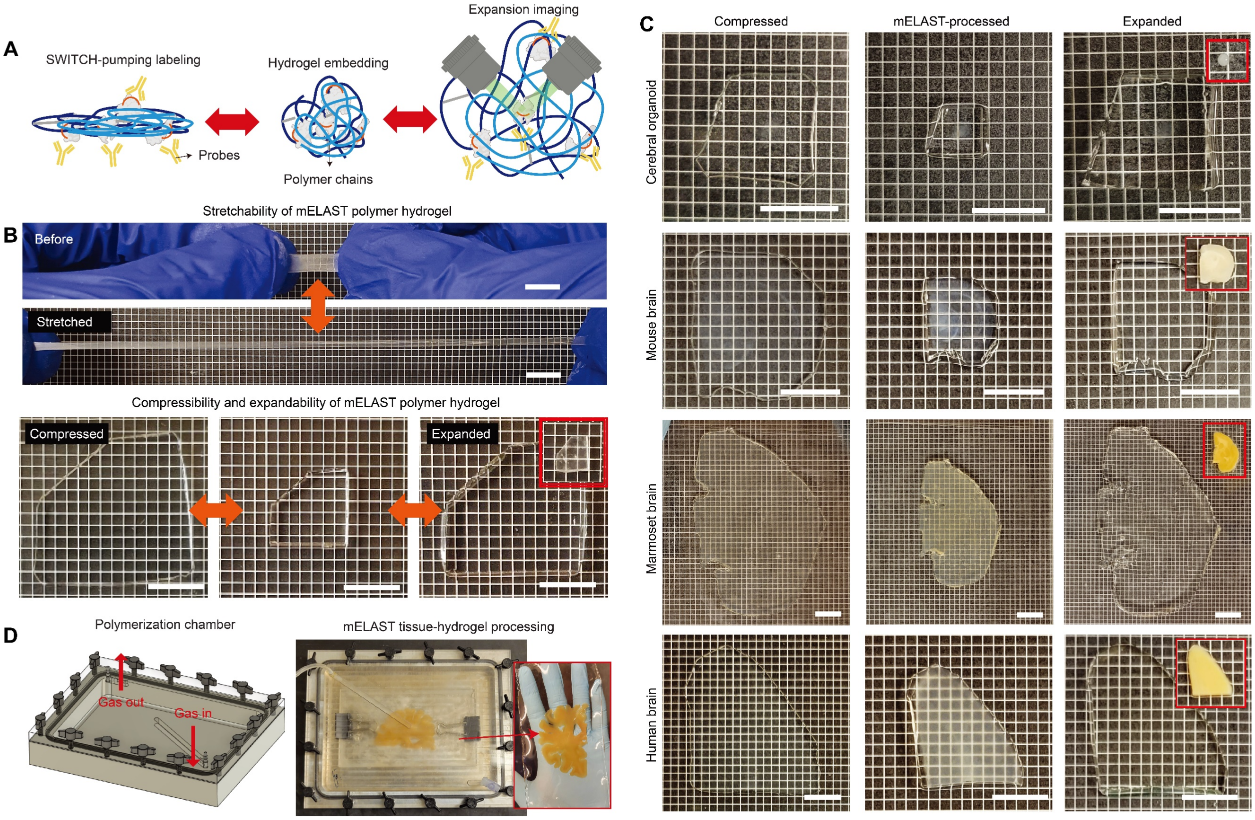


### Fig. S7.

**mELAST polymer hydrogel and tissue engineering.** (**A**) Schematic illustration of the mELAST mechanism. mELAST tissues can be rapidly, cost-effectively stained using the SWITCH-pumping staining method and imaged at multiple scales by physical expansion. (**B**) mELAST polymer hydrogel is stretchable, compressible and size-adjustable depending on buffer solution due highly entangled and super-absorbent polymeric chains. Scale bars, 1 cm. (**C**) mELAST-processed cerebral organoid, mouse, marmoset, and human brain tissue-hydrogels are elastic and expandable. Scale bars, 1 cm. (**D**) An air-tight chamber is used for mELAST gelling of human brain tissue. A SHIELD-delipidated tissue equilibrated in mELAST solution is placed between two glass plates with spacers, clamped with binder clips, and located inside the chamber. The chamber is closed and purged with Nitrogen gas for initiation of the polymerization.

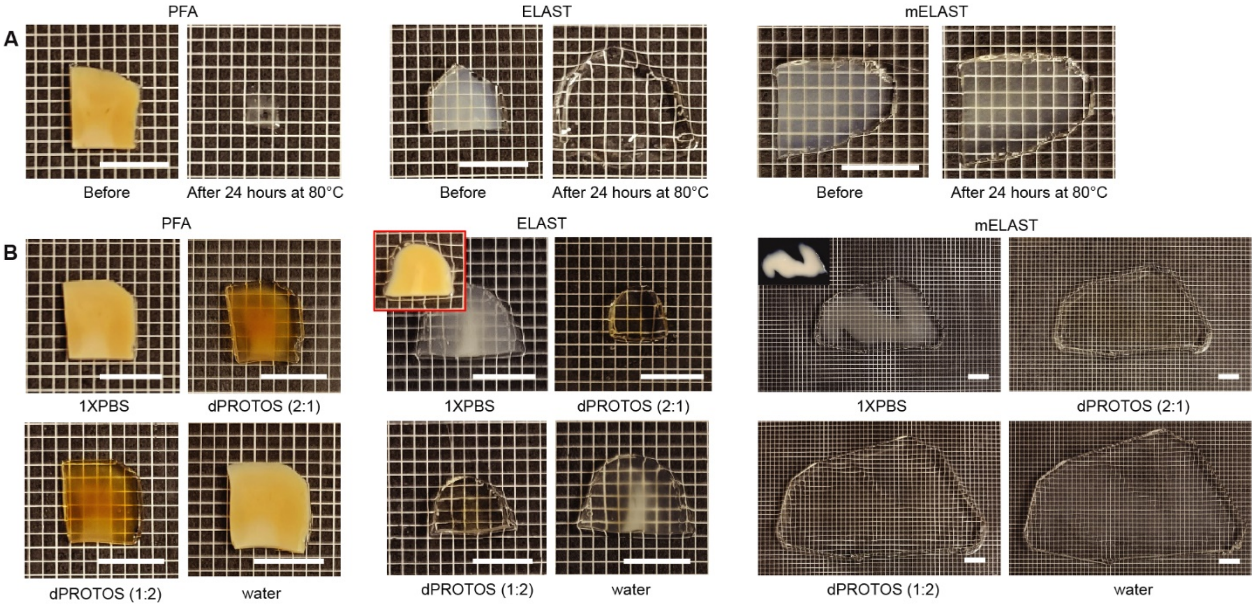


**Fig. S8.**

**Expansion and thermochemical stress test of mELAST human brain tissue-hydrogel.** (**A**) Thermal stress test to PFA, ELAST, and mELAST human brain tissue-hydrogels. After applying stress in the SDS clearing solution at 80°C for 24 hours, the tissue shapes before and after were compared. (**B**) Expansion test to PFA, ELAST, and mELAST human brain tissue gels in 1XPBS, dPROTOS (2:1 ratio with water for half-step and then full dPROTOS step), dPROTOS (1:2 ratio with water for half-step and then full dPROTOS step), and water. dPROTOS (1:2) was for collecting multi-scale dataset. Scale bars, 1 cm.

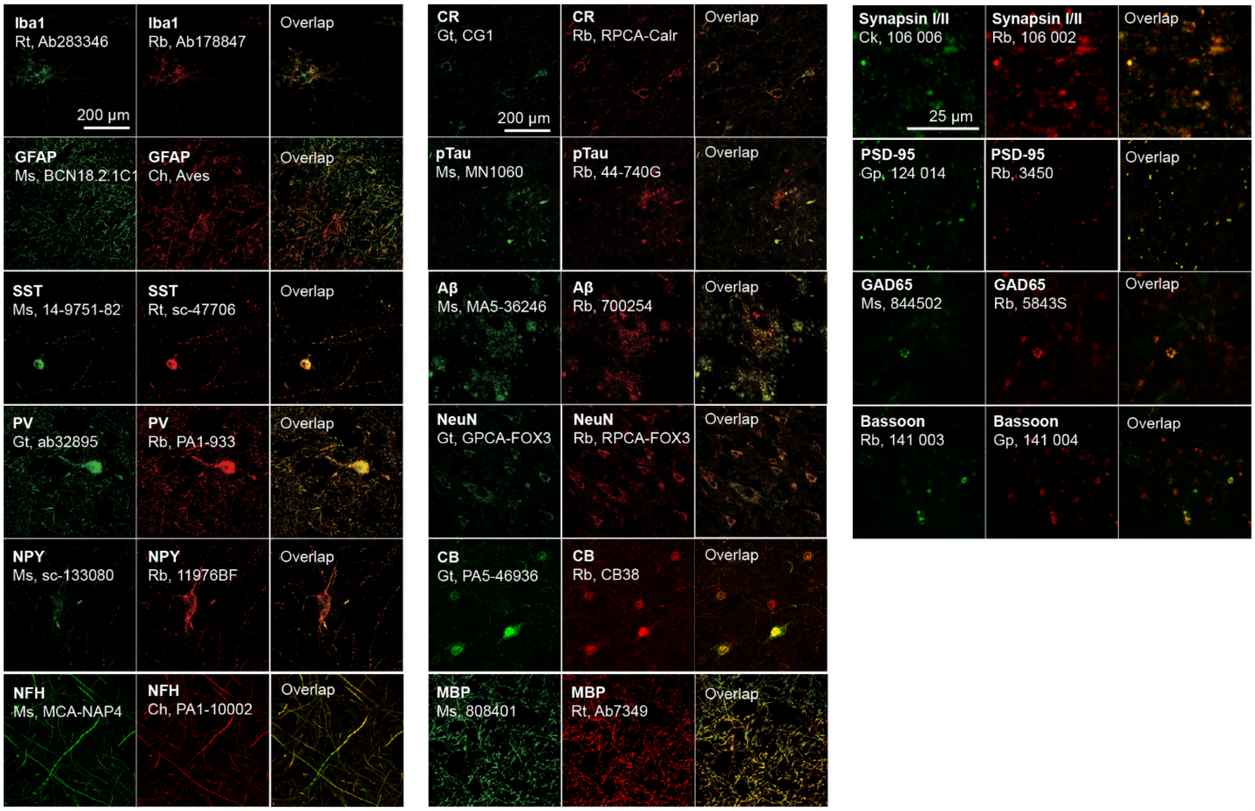


**Fig. S9.**

**Cross-species antibody validation for mELAST human brain tissues.** mELAST human brain tissues were co-stained with two antibodies from different host species but against same target proteins and imaged after expansion in water using a confocal microscope with the 20x/0.5NA water objective for the first and second columns and 63x/1.2NA water objective the third column.

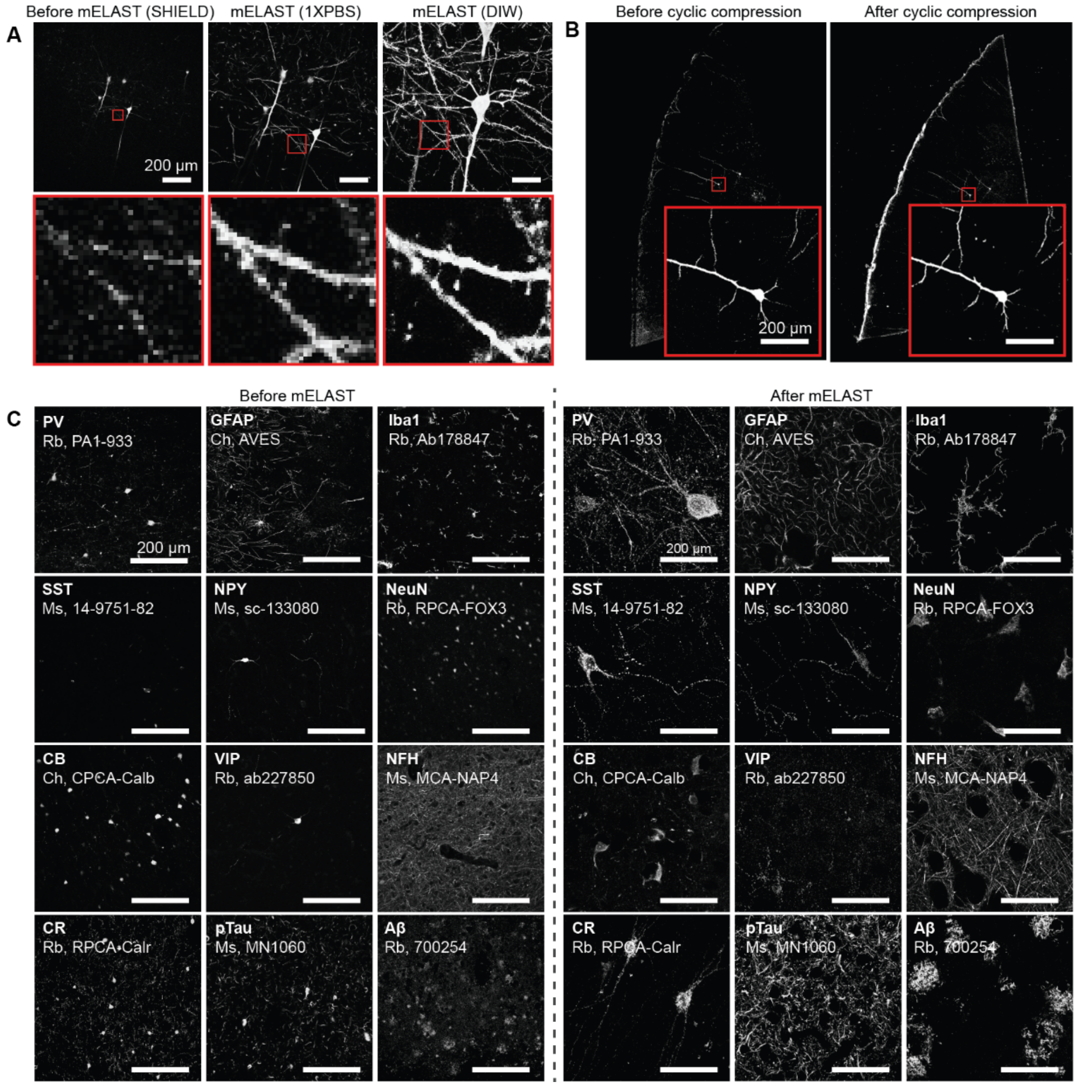


**Fig. S10.**

**mELAST validation.** (**A**) The same neurite in Thy1-eGFP M line mouse brain tissue was imaged before (SHIELD-cleared) and after mELAST processing (1XPBS, DIW). mELAST processing and tissue expansion does not damage neuron morphology while increasing imaging resolution using the same microscope conditions. (**B**) The overall integrity and microscale neuronal morphology of the mELAST mouse brain tissue were preserved without damage after 2000 cycles of 5-fold compression. (**C**) Antibody compatibility test to human brain tissue before and after mELAST processing and expansion in water. Scale bars, 200 µm. The10x/0.4NA objective was used for A-B and the 20x/0.5NA objective was used for C.

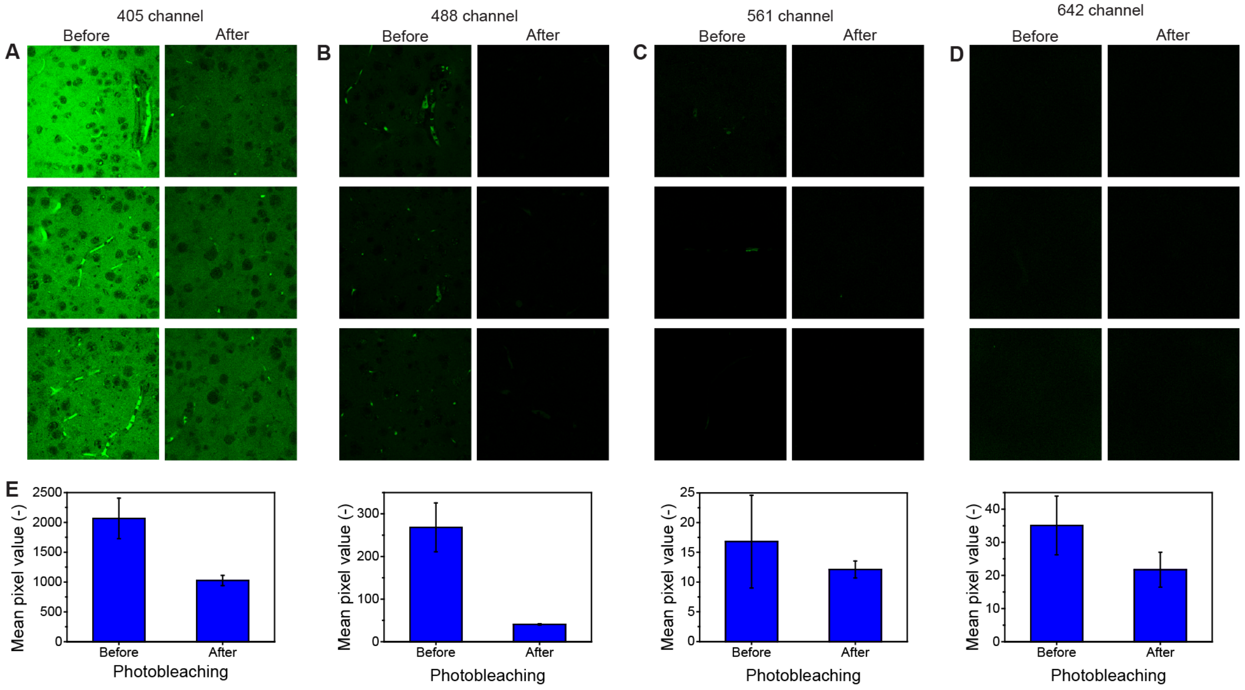


**Fig. S11.**

**Characterization of photobleaching of mELAST human brain tissues.** Autofluorescence imaging using MegaSPIM (2x objective) before and after 2 days of photobleaching for 405 channels (**A**), 488 channel (**B**), 568 channel (**C**), and 647 channel (**D**). The mean pixel values for all the cases were calculated and compared with each other.

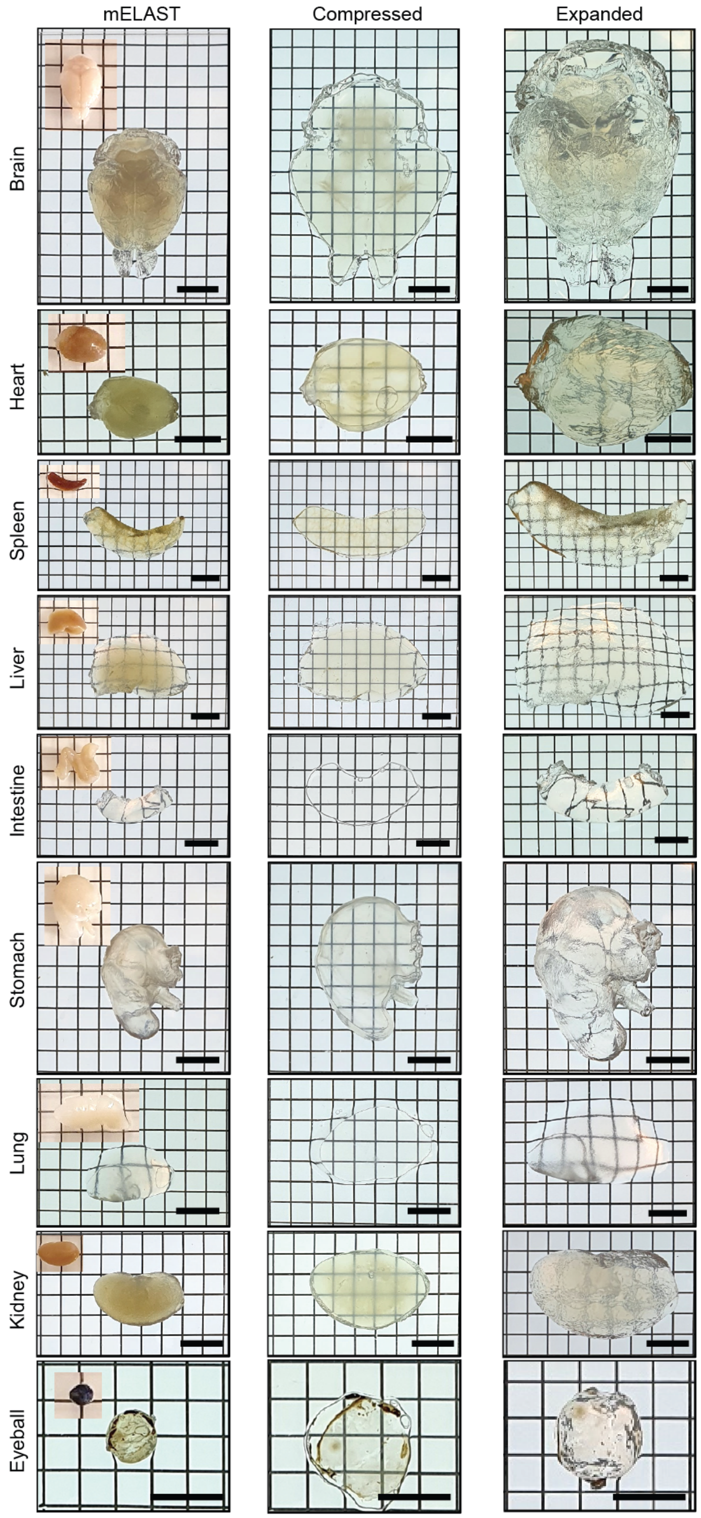


**Fig. S12.**
**mELAST processing of mouse organs**. Scale bars, 1 cm.

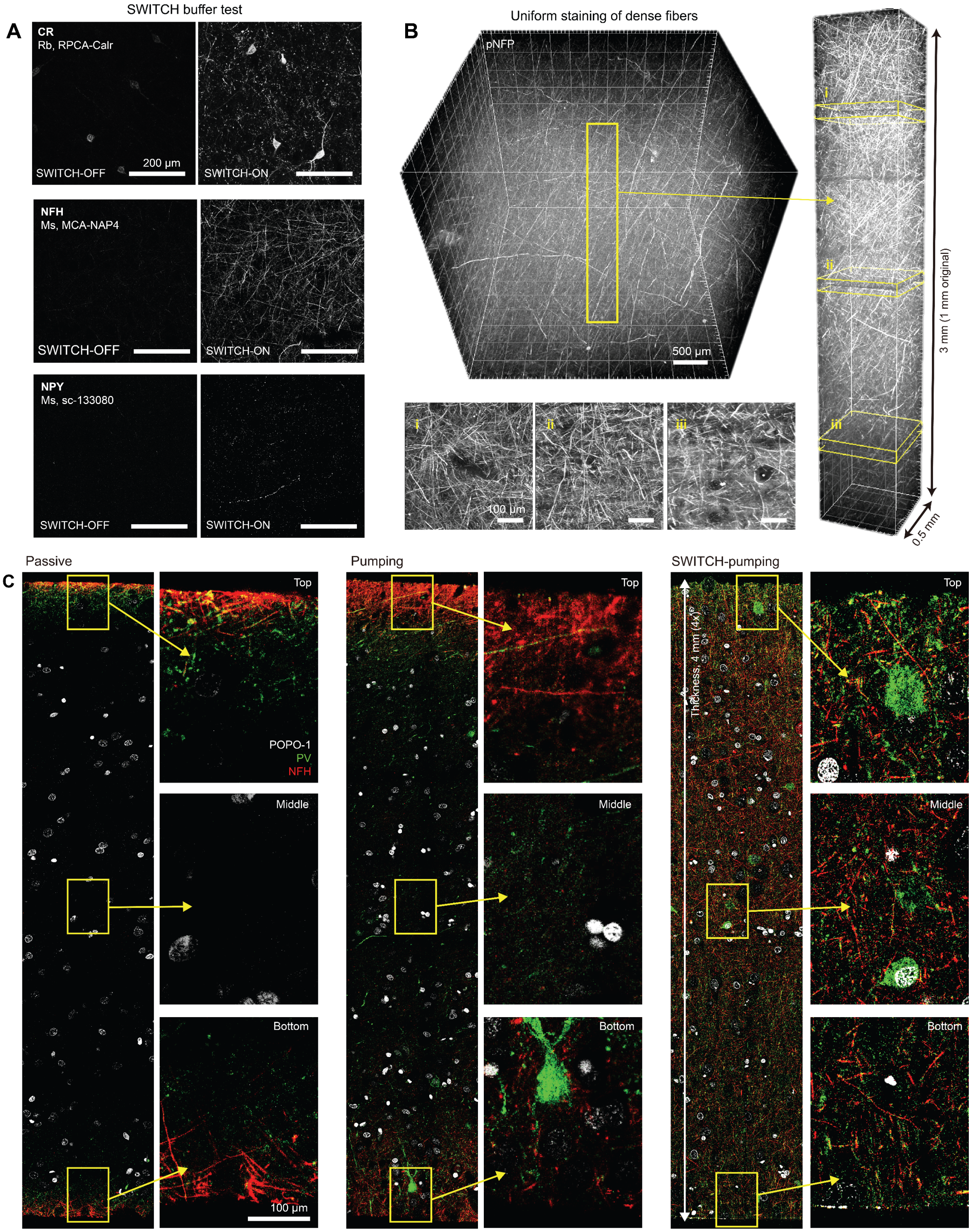


**Fig. S13.**

**SWITCH-pumping method.** (**A**) Staining test of NaCh-based SWITCH buffers to mELAST human brain tissues. SWITCH-OFF buffer was 1XPBSN with 5% NaCh and SWITCH-ON buffer is 1XPBSN with 5% NaCh and 1% Triton-X. The signal was compared with each other after 2 hours of primary and secondary staining. The result shows that the SWITCH-OFF buffer successfully slows down antibody-binding and the binding affinity can be recovered by ON step. (**B**) SWITCH-pumping staining of mELAST human brain tissues with anti-pNFP antibody (a pan-axonal marker) and MegaSPIM imaging after 3x expansion (16.7x objective). (**C**) SWITCH-pumping staining allows rapid, uniform, and cost-effective immunostaining of mELAST tissue-hydrogels. Three same-sized mELAST human brain tissue-hydrogels were stained with POPO-1 (nuclear dye), anti-PV, and anti-NFH antibodies using passive, pumping and SWITCH-pumping methods, and the staining uniformity was compared. Scale bar, 100 μm.

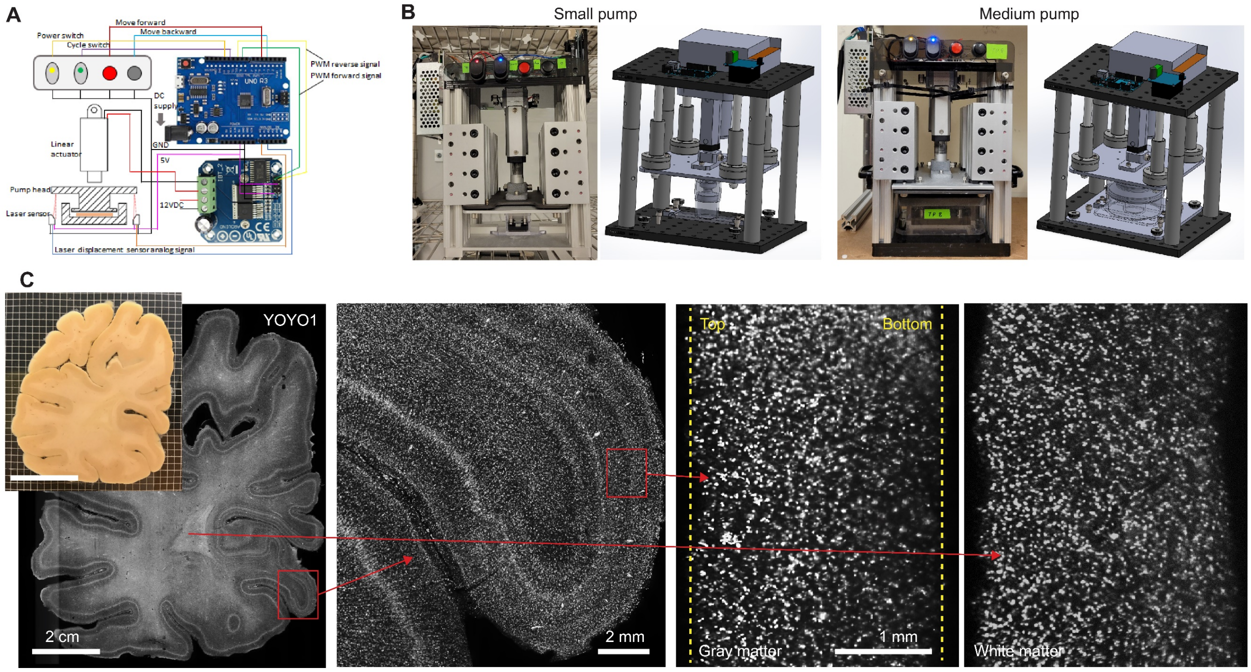


**Fig. S14.**

**Design and fabrication of tissue pumps of various sizes.** (**A**) Circuit design of tissue pumps. (**B**) Photos and 3D models of the small and medium-sized tissue pumps (the big-sized pump is displayed in Fig. 4G). (**C**) Ultra-large-scale tissue staining (mELAST processed, ~10 cm x 8 cm x 2.5 mm) with YOYO-1 nuclei dye using the big tissue pump.

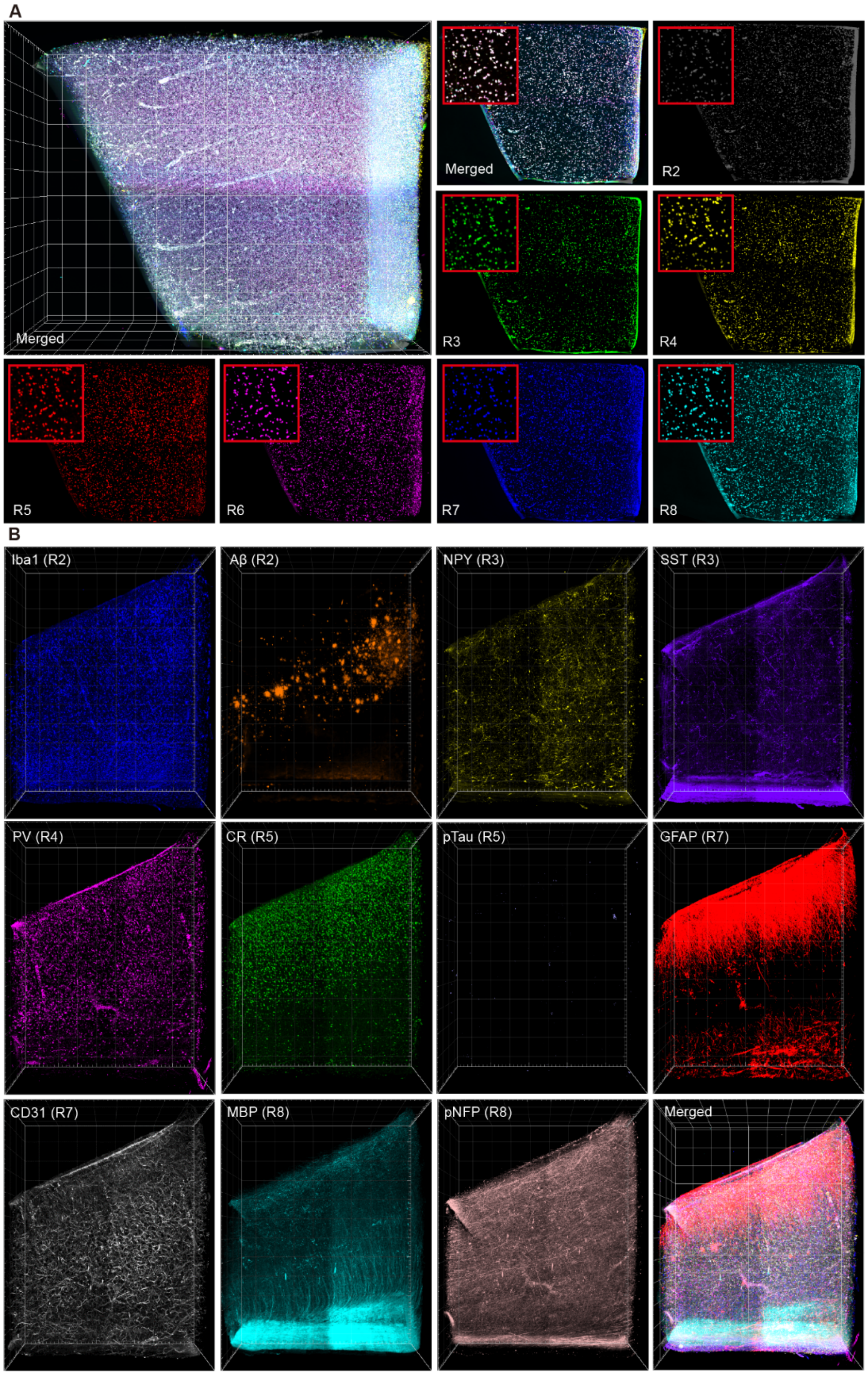


**Fig. S15.**

**Co-registration of multi-round images from the mELAST human brain tissue (Control).** (**A**) Co-registration of seven rounds images (rounds 2-8) by detecting nuclei

(YOYO-1) and warping based on the matching points. (**B**) The same transformation matrix is applied to other channels for multiplexing.

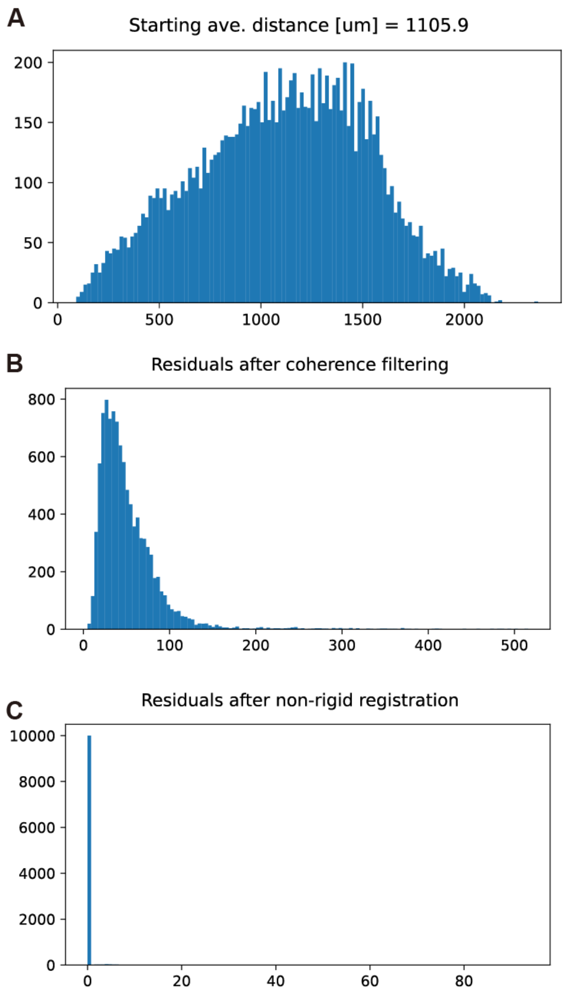


**Fig. S16.**

**Near perfect co-registration of nuclei (YOYO-1) from multi-rounds images.** (**A**) Histogram of distances between corresponding nuclei in different rounds (round 2 and 8), where the moving image has been projected into the fixed image coordinates. (**B**) Histogram of errors between matching nuclei after affine transform, RANSAC, and coherence filtering. (**C**) Histogram of distances between matching nuclei after final round of nonrigid registration.

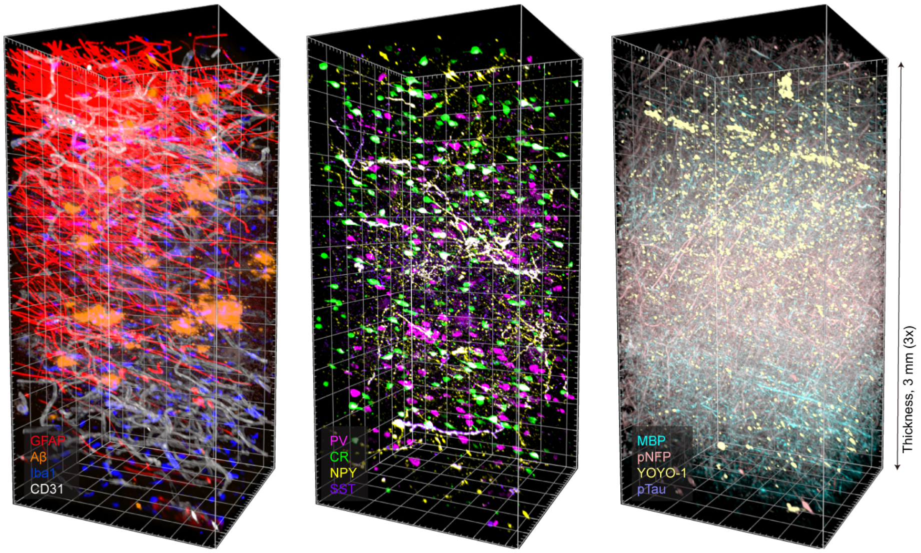


**Fig. S17.**

**Highly multiplexed imaging of an mELAST human brain tissue.** The mELAST cortex tissue was multi-round SWITCH-pumping stained and MegaSPIM-imaged, and the images were computationally co-registered based on nuclei dye patterns included in each round.

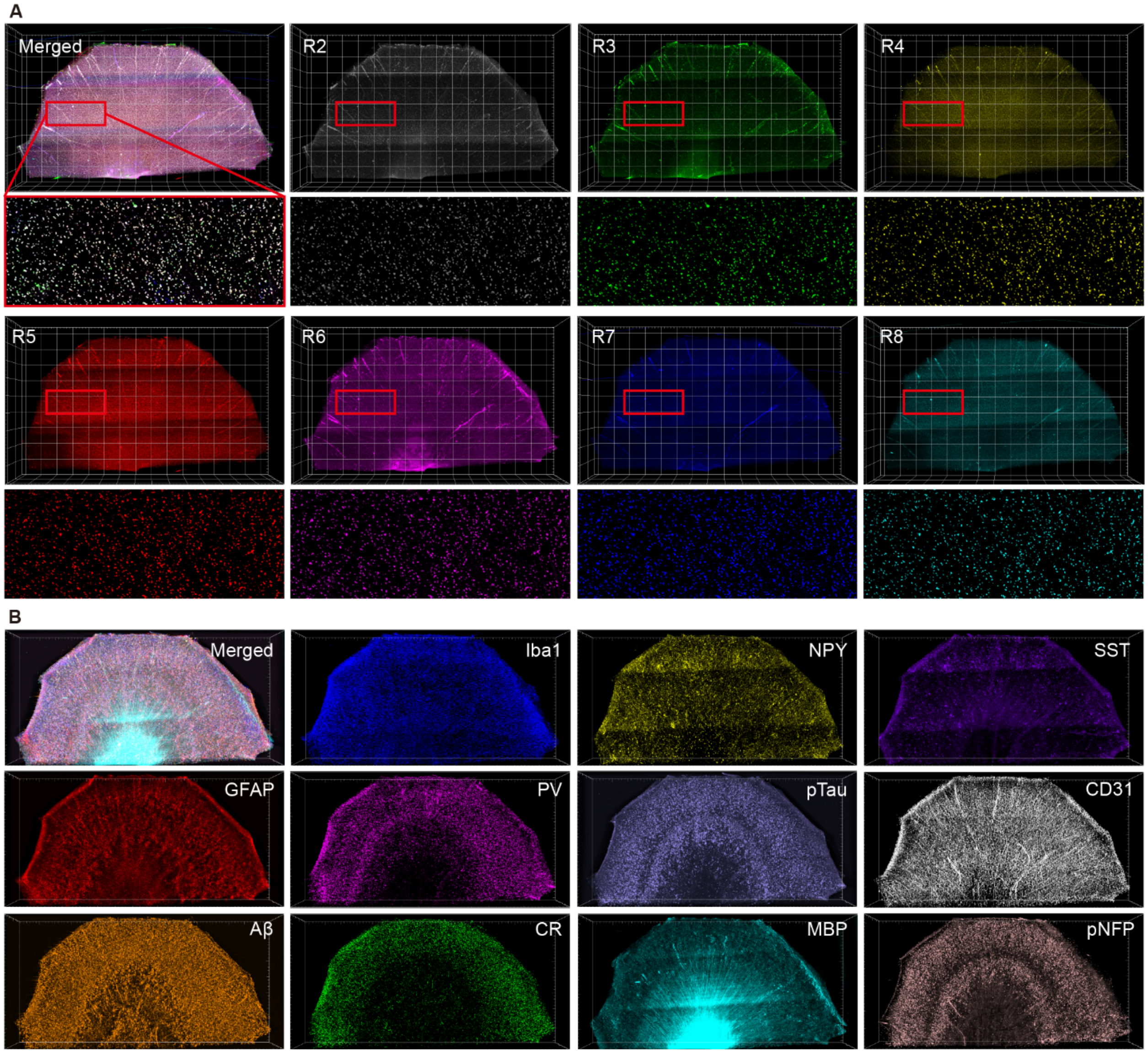


**Fig. S18.**

**Co-registration of multi-round images from the mELAST human brain tissue (AD).** (**A**) Co-registration of seven rounds images (rounds 2-8) by detecting nuclei (YOYO-1) and warping based on the matching points. (**B**) The same transformation matrix is applied to other channels for multiplexing.

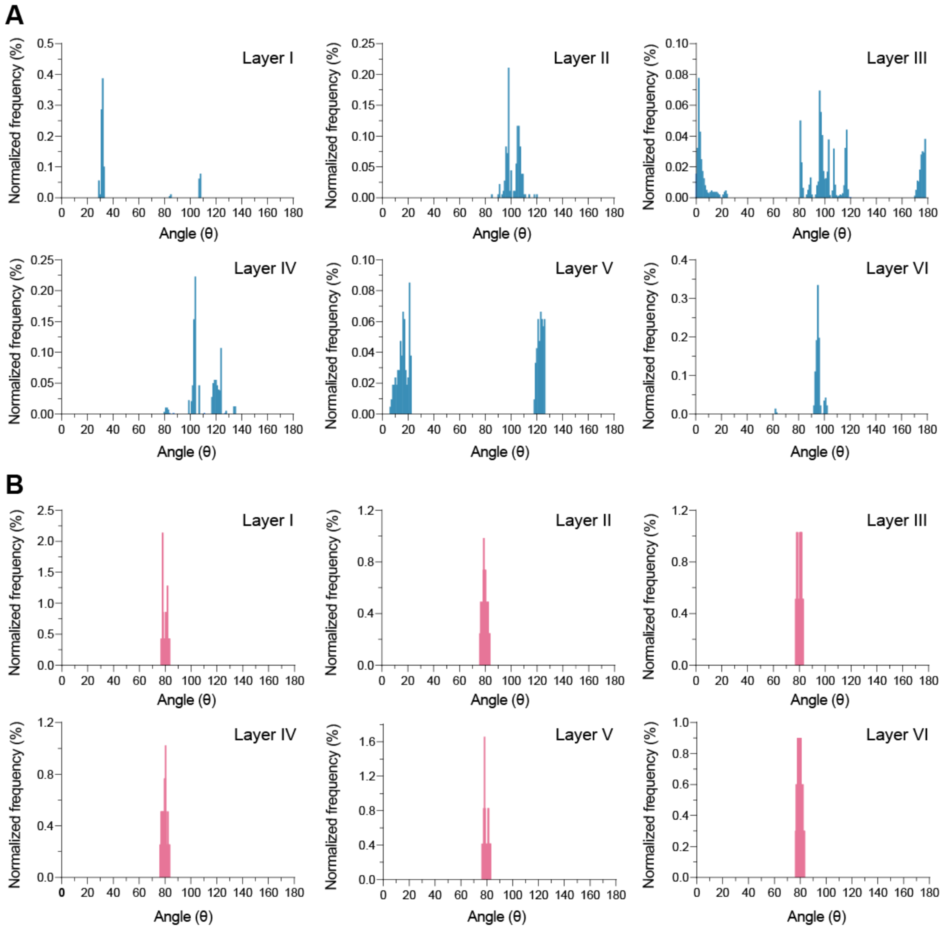


**Fig. S19.**

**Layer-specific histograms of pNFP+ neural fiber angular orientation with respect to cortical column direction (θ) of individual cortical layers in the control and AD tissues**; (**A**) Control (**A**) and AD (**B**) tissues.

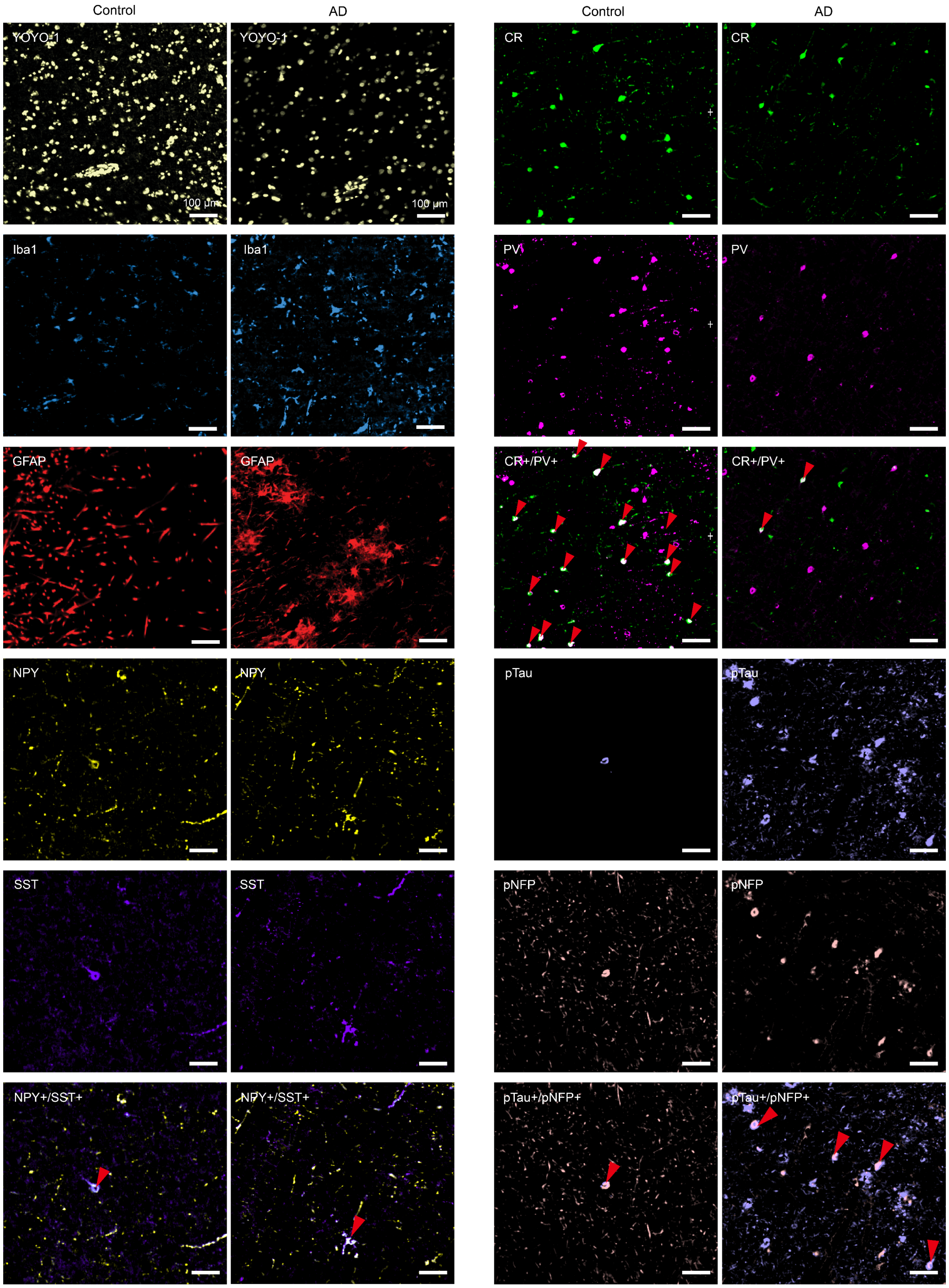


**Fig. S20.**

**Representative images of each cell type in layer III of the mELAST-processed control and AD human brain tissues.**

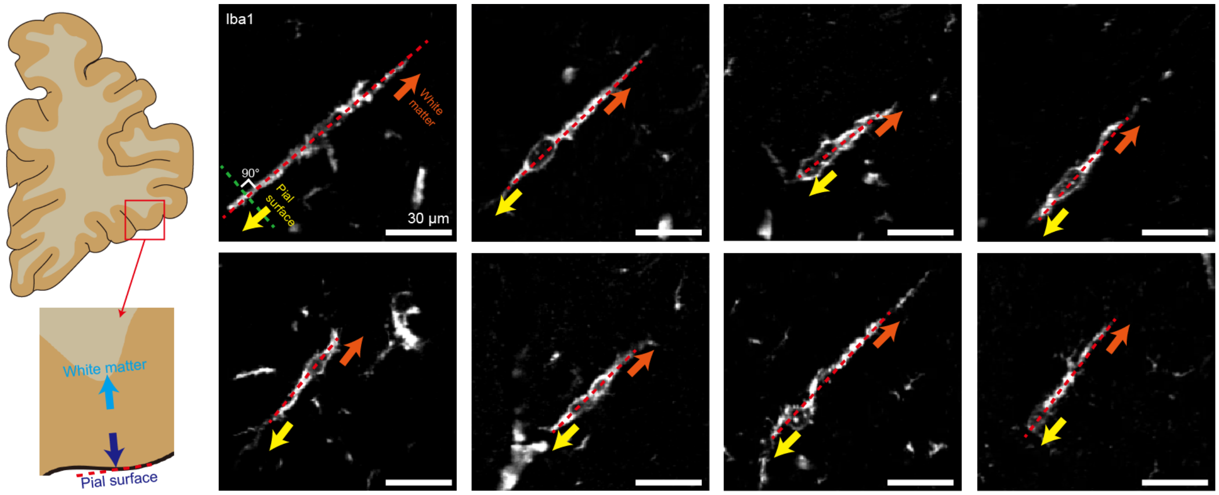


**Fig. S21.**

**Representative images of Iba1+ rod-shaped microglia in layer III in the mELAST-processed AD human brain tissue.**

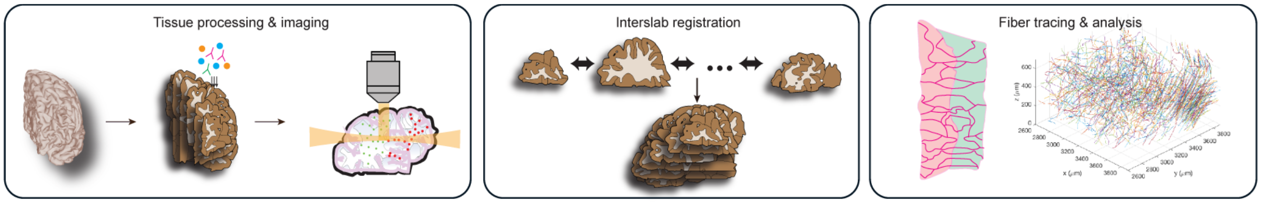


**Fig. S22.**

**Projectome mapping of large intact tissue requires processing, staining, and imaging of sliced mm-thick slabs followed by computational reconstruction, axon tracing, and downstream analysis.**

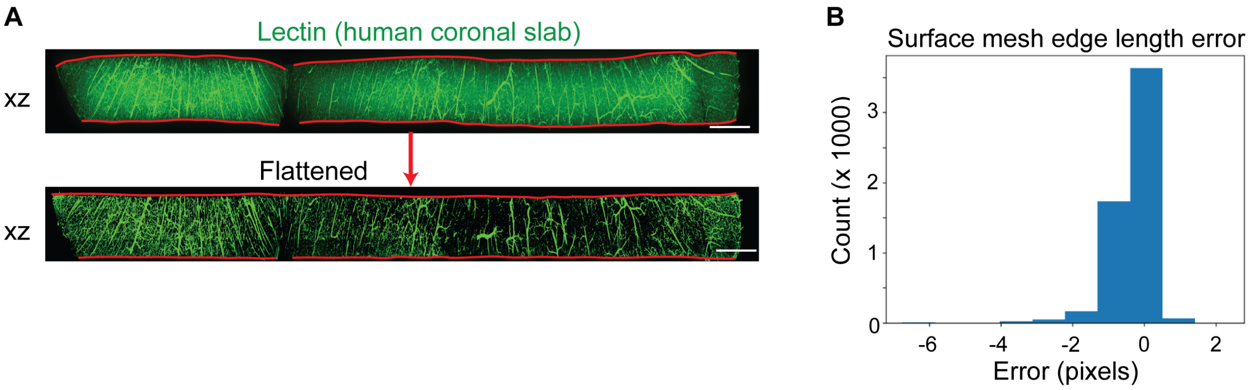


**Fig. S23.**

**UNSLICE surface flattening preserves surface topology.** (**A**) Human coronal slabs stained with lectin corresponding to the top slab from Fig. 7D-E were computationally flattened using Boundary First Flattening. Top and bottom surfaces were outlined in red. Scale bars, 2 mm. (**B**) Histogram showing the differences in edge lengths of the original slab surface mesh and the edge lengths of the flattened surface mesh. The root mean square error (RMSE) of the edge lengths is 0.67 pixels, with a variance of 0.33 pixels.

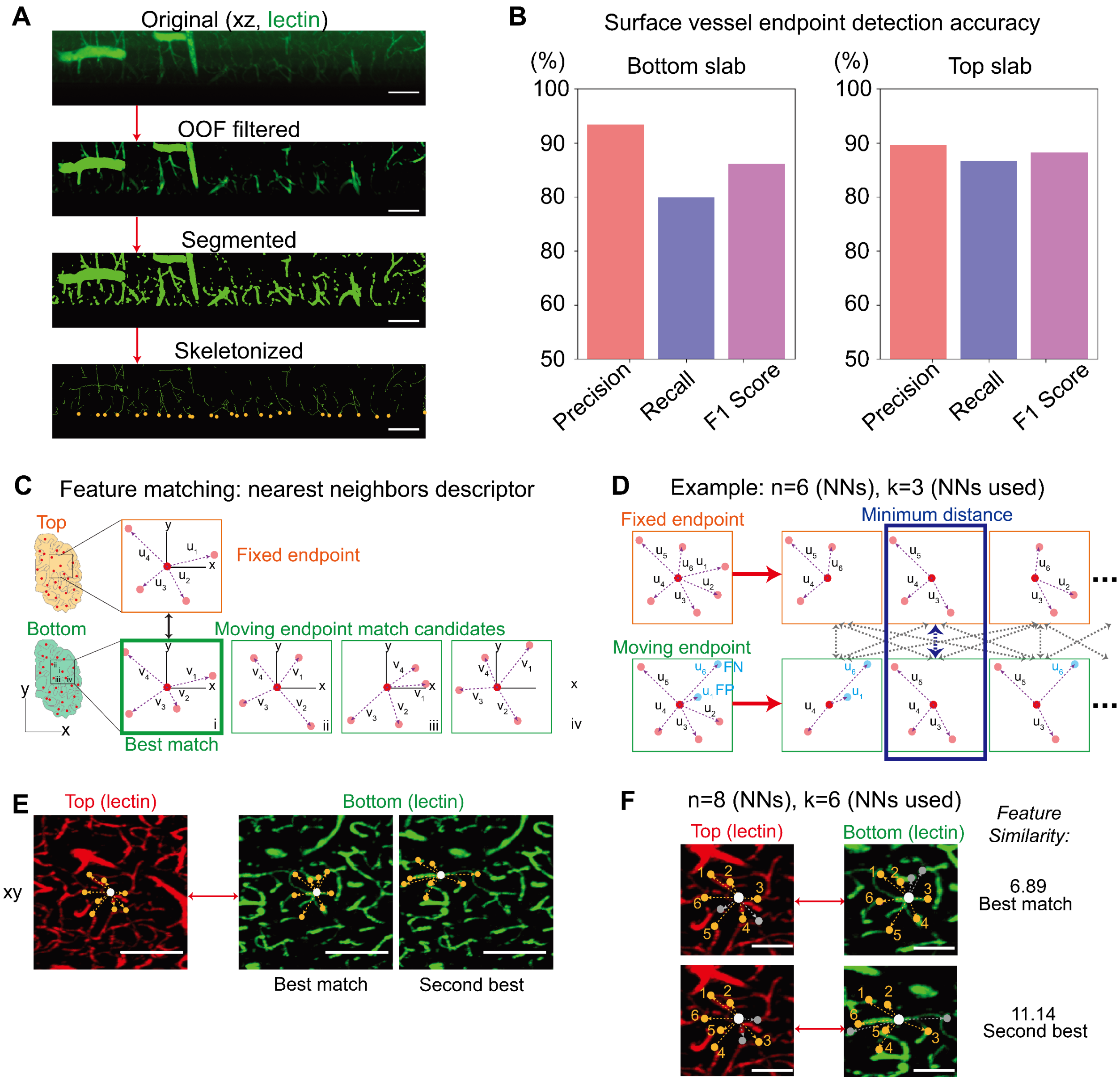


**Fig. S24.**

**UNSLICE endpoint detection and keypoint descriptor enables robust feature matching.** (**A**) Vessel (or fiber) surface endpoints were detected via the Optimally-Oriented Flux filter, intensity threshold segmentation, and skeletonization. The image subvolume was from the coronal human slab data in Fig. 7D-E. Scale bars, 200um. (**B**) F1 scores for endpoint detection in the human hemisphere slab data were 86.1% in the bottom slab and 88.1% in the top slab. Precision, which measures the ratio of true positives detected to all detected points (i.e., true positives + false positives), was 93.4% and 89.6% in the bottom and top slabs, respectively. Recall, which measures the ratio of true positives to all actual endpoints (i.e., true positives + false negatives), was 79.8% and 86.7% in the bottom and top slabs, respectively. (**C**) The keypoint descriptor is the concatenation of relative vectors to an endpoint's *n* nearest neighbors. The Euclidean distance between the fixed and candidate moving descriptors determines the “best match” for each endpoint. (**D**) To be robust to false negatives (FN) and positives (FP) depicted in cyan, UNSLICE’s feature matching strategy used the Hungarian algorithm to compute the minimum distance between a given candidate moving and fixed keypoint descriptor when using any *k* of the *n* nearest neighbors. (**E**) Two candidate bottom slab endpoints potentially matched with a top slab endpoint, with each point’s 8 nearest neighbors shown in orange. Scale bars, 200 μm. (**F**) When using 8 nearest neighbors and *k*=6 to compute the feature similarity, we found that the feature similarity ratio of the best match to the second-best match is 6.89/11.14 = 62% < 85%, satisfying the criterion to accept this pair as a feature match. Scale bars, 100 μm.

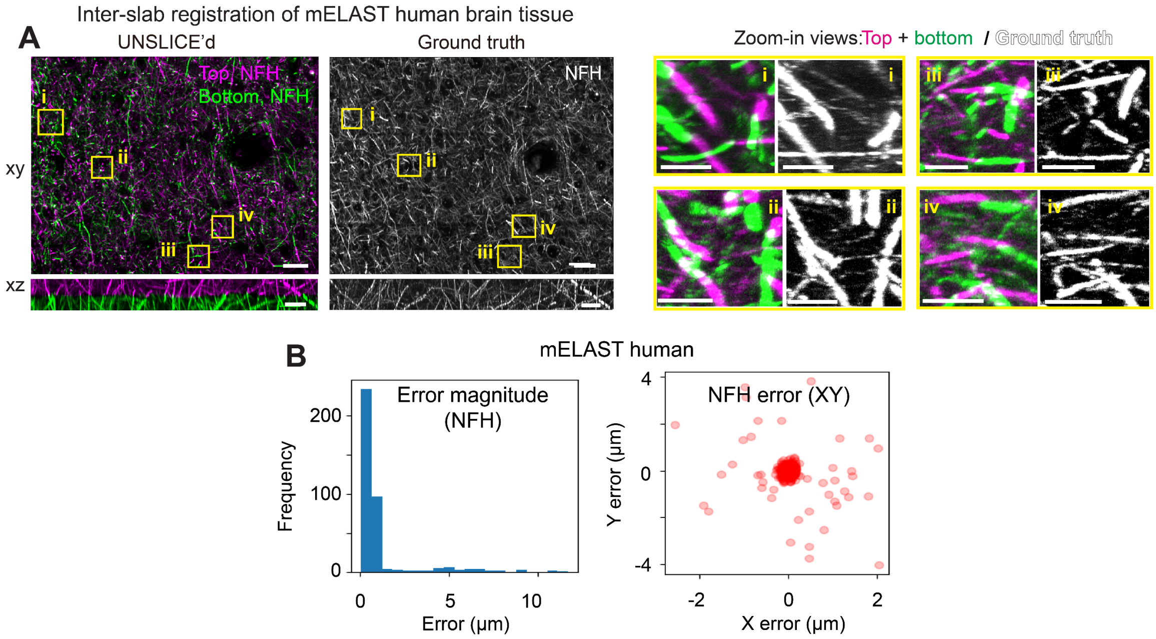


**Fig. S25.**

**Ground-truth validation of UNSLICE for axon-level connectivity.** (**A**) UNSLICE enables 3D reconstruction of sliced, expanded human mELAST-processed tissue at axon-scale resolution. Compared to the intact tissue imaged pre-slicing (ground truth), UNSLICEd images show restoration of axonal connectivity between slabs. Scale bars, 100 µm (macroscopic view) and 25 µm (zoom-in view). (**B**) Validation of UNSLICE evaluated by quantifying the distance (error) between fiber endpoints at the cut surface in three sampled subsections based on ground-truth image data before and after slicing at subcellular resolution; the expanded mELAST human brain tissue; average error 1.13 µm.

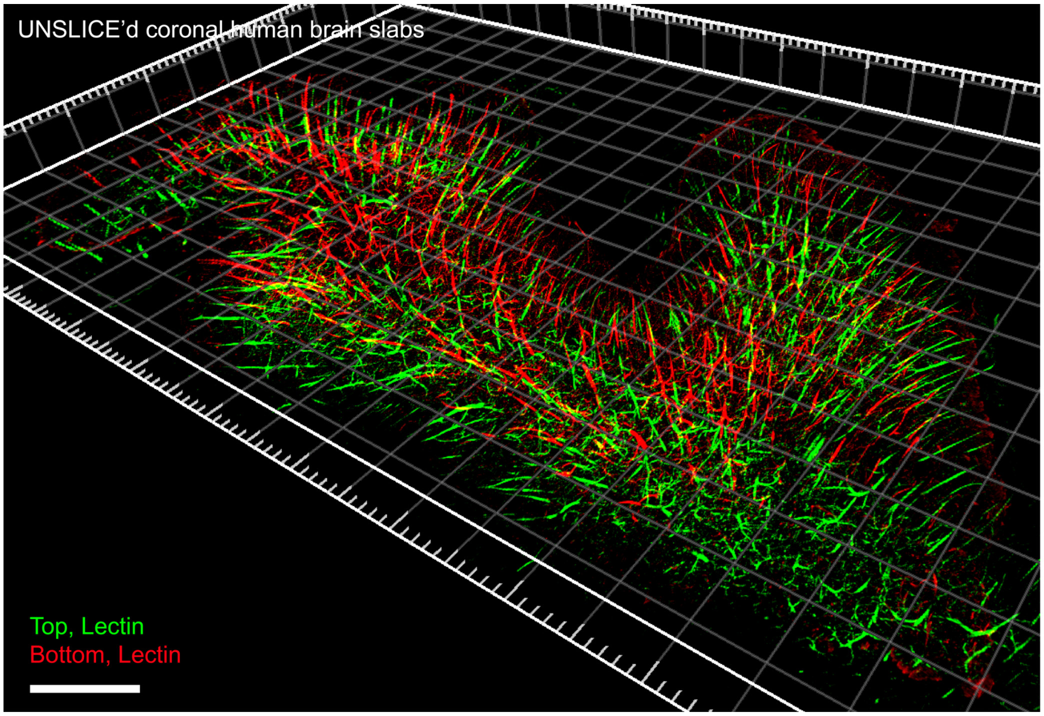


**Fig. S26.**

**3D rendering of the UNSLICE’d coronal human brain hemisphere slabs. Scale bar, 5 mm**.

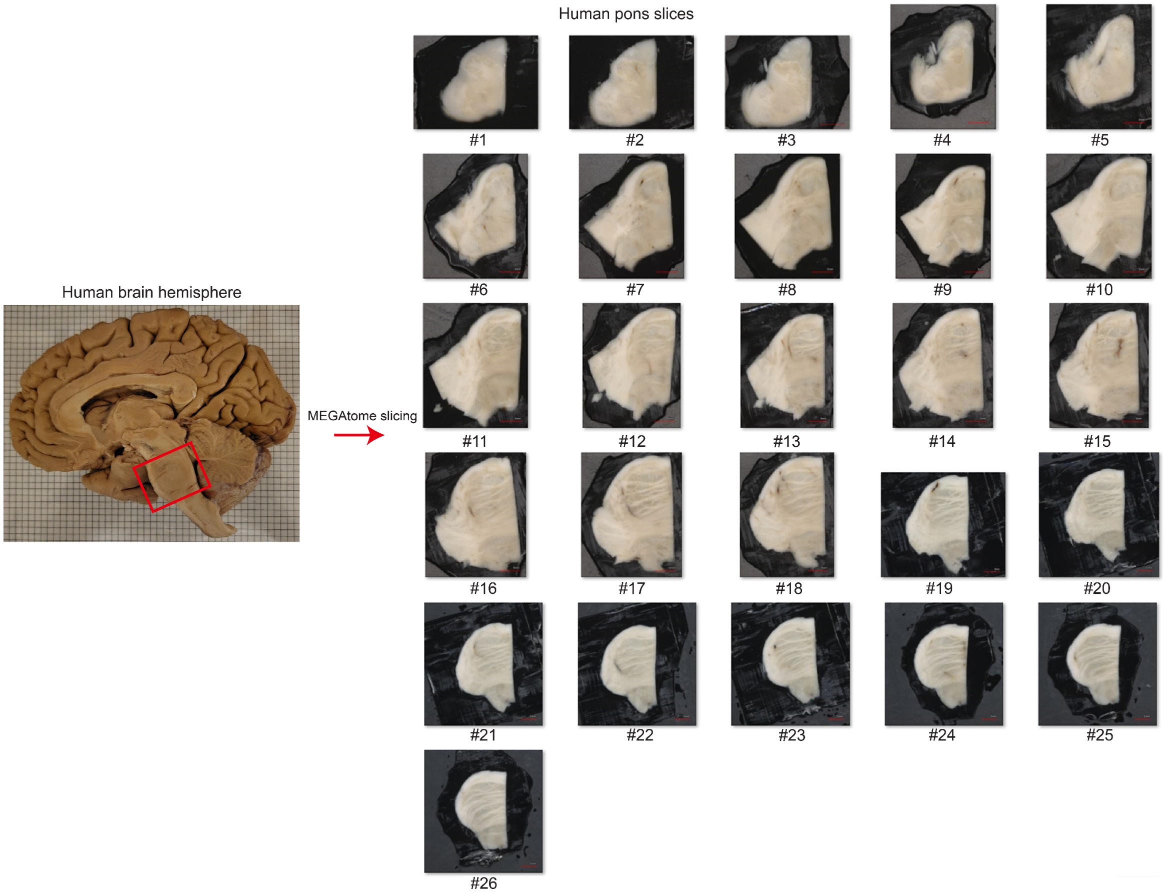


**Fig. S27.**

**Human pons was cut from a human brain hemisphere and sliced into 1 mm-thick slabs using MEGAtome.**

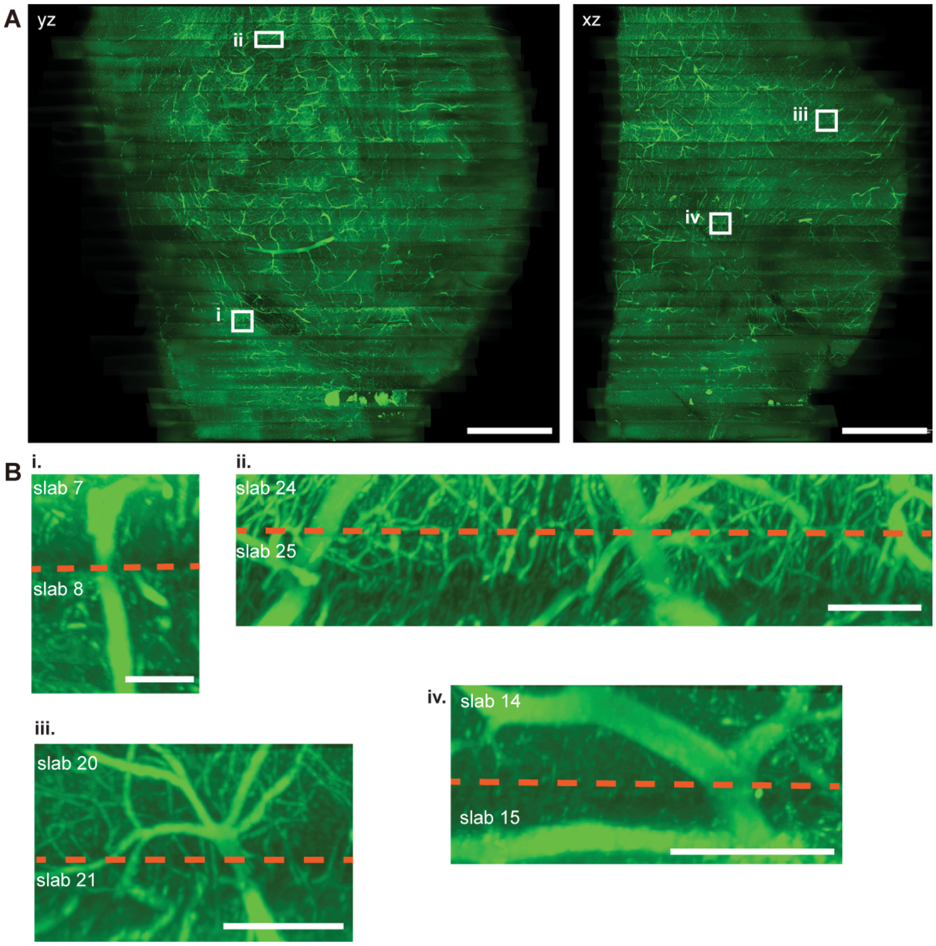


**Fig. S28.**

**3D pons reconstruction using UNSLICE.** (**A**) xz and yz profile views of the reconstructed pons for lectin, maximum intensity projection over the middle 50 slices in y and x, respectively. Scale bars, 5 mm. (**B**) Zoomed in high resolution views of the lectin channel at the interface between (i) the #7 and #8 slabs, (ii) the #24 and #25 slabs, (iii) the #20 and #21 slabs, and (iv) the #14 and #15 slabs. Scale bars, 200 μm.

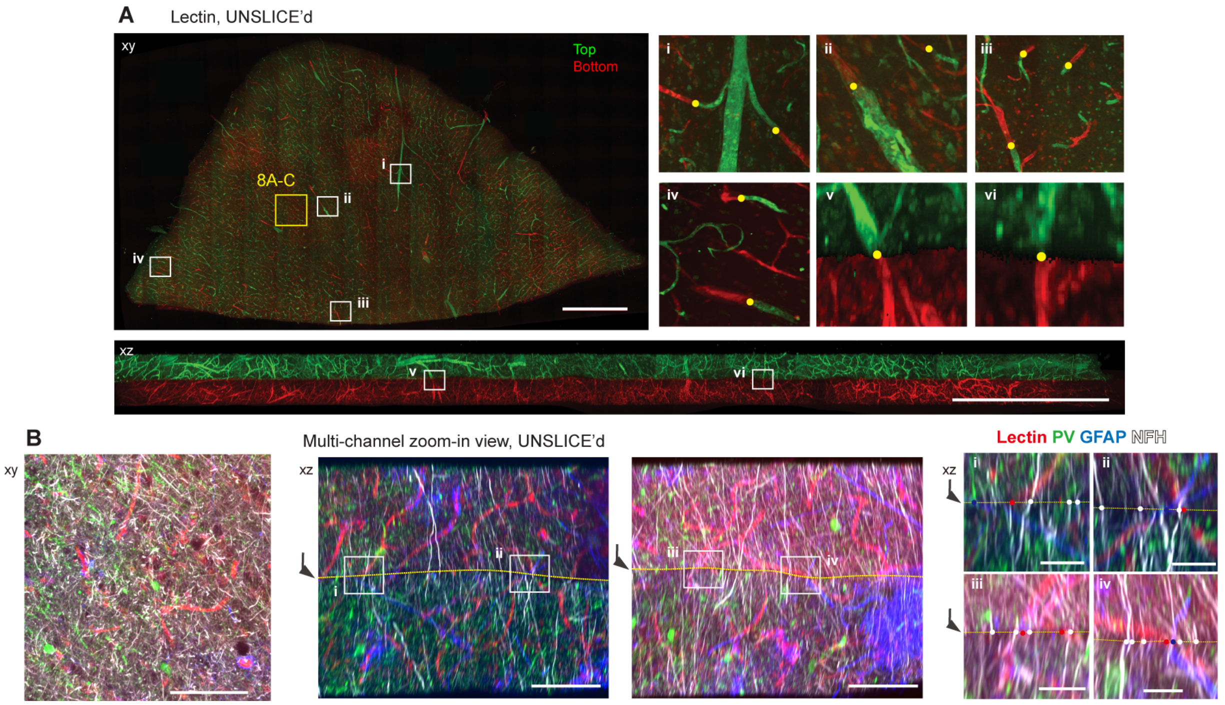


### Fig. S29.

**Multi-channel projectome mapping.** (**A**) Using lectin, two cortical human tissue slabs were inter-slab-registered to single-vessel resolution, providing a groundwork for fine-tuning using the other neural fiber channels. Scale bars, 1 mm. (**B**) UNSLICE enables multi-channel reconstruction at single-axon resolution. Zoomed in section from (A) with overlay of GFAP, lectin, NFH, and PV showing multi-channel reconstruction. Scale bars, 100 µm and 25 µm (zoomed-in images).

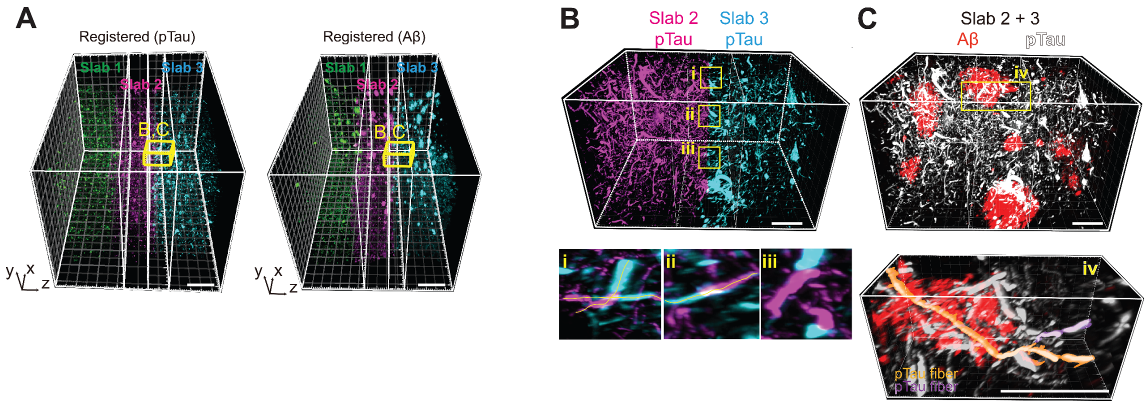


**Fig. S30.**

**Zoom-in view of pTau+ fiber reconstruction, segmentation, tracing, and colocalization with Aβ at slab 2-3 interface.** (**A**) Macroscopic view of UNSLICE'd mELAST-processed AD human brain tissues, stained for pTau and Aβ. Scale bars, 0.5 mm. (Same as Fig. 8D). (**B**) 3D rendering of a subvolume at the slab 2 / slab 3 interface, as indicated by the prisms drawn in (A). pTau+ fibers were reconstructed and traced across the interface, as shown in the xy maximum intensity projections shown in insets (i)-(iii). Scale bars, 150 µm. (**C**) Overlay of the reconstructed Aβ channel (red) and pTau channel (white) of the same subvolume in (B). Inset (iv) shows two pTau fibers that was segmented via our axon tracing algorithm. Scale bars, 150 µm.

**Table S1.**

Resolution information of each objective for MegaSPIM imaging. Raw optical xy resolution is the pixel size as seen by the camera in the 45-degree tilted image plane through which the light-sheet illuminates. De-obliqued xyz voxel size is after the voxels are transformed back into traditional xyz coordinates based on the calculated choice of x-step size, typically the xy voxel size divided by √2. Effective resolution is relative to the tissue expansion, and the scale at which we visualize the biological structures or cell types of interest.

| Objective Magnification | Numerical Aperture | Raw optical xy pixel resolution (µm) | De-obliqued xyz voxel size (µm) | Effective xyz resolution with 3x tissue expansion (µm) |
|---|---|---|---|---|
| 2x | 0.1 | 3.46 x 3.46 | 2.44 x 3.46 x 2.44 | 0.81 x 1.15 x 0.81 |
| 4x | 0.2 | 1.74 x 1.74 | 1.23 x 1.74 x 1.23 | 0.41 x 0.58 x 0.41 |
| 15x | 0.4 | 0.48 x 0.48 | 0.34 x 0.48 x 0.34 | 0.11 x 0.16 x 0.11 |

**Table S2.**

Antibodies and dyes compatible with mELAST human brain tissues.

| Target | Vendor | Catalog # | Host species | Clone |
|---|---|---|---|---|
| Aβ | Invitrogen | MA5-36246 | Mouse IgG2b | M |
| Aβ | Invitrogen | 700254 | Rabbit | M |
| Bassoon | SYSY | 141 004 | Guinea pig | P |
| Bassoon | SYSY | 141 003 | Rabbit | P |
| CB | Encor | CPCA-Calb | Chicken | P |
| CB | SWANT | CB38 | Rabbit | M |
| CB | Invitrogen | PA5-46936 | Goat | P |
| CD31/PECAM-1 | R&D systems | AF3628 | Goat | P |
| CR | Abcam | Ab702 | Rabbit | P |
| CR | Encor | RCPA-Calr | Rabbit | P |
| CR | Swant | CG1 | Goat | P |
| GAD2 | CST | 5843S | Rabbit | M |
| GAD65/GAD2 | Biolegend | 844502 | Mouse IgG1 | M |
| GFAP | Santa Cruz | sc-33673 | Mouse IgG2b κ | M |
| GFAP | Aves | | Chicken | P |
| NeuN | Encor | RPCA-FOX3 | Rabbit | P |
| NeuN | Encor | GPCA-FOX3 | Goat | P |
| NeuN | Abcam | ab104225 | Rabbit | P |
| PV | Invitrogen | PA1-933 | Rabbit | P |
| PV | Abcam | ab32895 | Goat | P |
| pNFP (SMI-312) | Biolegend | 837904 | Mouse IgG1 & IgM | P |
| MBP | Abcam | Ab7349 | Rat | M |
| MBP | Biolegend | 808401 | Mouse IgG2b | M |
| Iba1 | Abcam | Ab178847 | Rabbit | M |
| Iba1 | Abcam | Ab283346 | Rat IgG2a | M |
| NFH | Thermo | PA1-10002 | Chicken | P |
| NFH | Encor | MCA-NAP4 | Mouse IgG1 | M |
| NPY | CST | 11976s | Rabbit | M |
| NPY | Santa Cruz | Sc133080 | Mouse IgG1 | M |
| SST | Santa Cruz | Sc-47706 | Rat | M |
| SST | eBiosci | 14-9751-82 | Mouse IgG1 | M |
| Synapsin I/II | SYSY | 106 102 | Rabbit | M |
| Synapsin I/II | SYSY | 106 006 | Chicken | P |
| pTau | Invitrogen | MN1060 | Mouse IgG1 | M |
| pTau | Invitrogen | 44-740G | Rabbit | P |
| PSD-95 | Abcam | ab12093 | Goat | P |
| PSD-95 | CST | 3450 | Rabbit | M |
| PSD-95 | SYSY | 124 014 | Guinea Pig | P |
| YOYO-1 | Invitrogen | Y3601 | - | - |
| POPO-1 | Invitrogen | P3580 | - | - |
| Lectin-649 | Vector | DL-1177 | Tomato | - |

**Table S3.**

Number of NeuN+ cells in the non-dementia control and AD intact human brain coronal slabs (Fig. 5A-B).

| Region | Control | AD |
|---|---|---|
| Superior frontal gyrus [SFG] | 2,204,877 | 753,033 |
| Mid frontal gyrus [MFG] | 2,408,979 | 898,503 |
| Orbital gyrus [OrG] | 1,153,566 | 464,514 |
| Gyrus rectus [ReG] | 490,300 | 182,397 |
| Rostral gyrus [RoG] | 447,005 | 189,261 |
| Cingulate gyrus [CgG] | 760,000 | 403,150 |
| Total | 7,464,727 | 2,890,858 |

**Table S4.**
Tissue processing, staining and imaging information for data collection of each figure.

| Figure | Tissue type | Microscope | Staining buffer | Imaging buffer | Antibody |
|---|---|---|---|---|---|
| Fig. 2L | SHIELD human spinal cord | Dfly 20x obj. | 1XPBSNaCh | dPROTOS | Anti-NFH (Mouse IgG1, MCA-NAP4, Encor) + RRX donkey anti-mouse (715-295-150, JacksonImmunoResearch Lab)<br>Lectin-649 (DL-1177, Vector) |
| Fig. 2M | SHIELD mouse brain | MegaSPIM 4x obj. | 1XPBSNaCh | dPROTOS | Lectin-488 (DL-1174, Vector)<br>Anti-V5 (Chicken, RPCA-FOX3, Encor) + RRX donkey anti-rabbit (711-295-152, JacksonImmunoResearch Lab) |
| Fig. 3D-E | SHIELD human brain | MegaSPIM 2x obj. | 1XPBSNaCh | dPROTOS | Anti-NeuN (Rabbit, RPCA-FOX3, Encor) + Alexa647plus donkey anti-rabbit (A32795, Invitrogen) |
| Fig. 3G | SHIELD mouse, marmoset brains | MegaSPIM 2x obj. | 1XPBSNaCh | dPROTOS | Anti-NeuN (Rabbit, RPCA-FOX3, Encor) + Alexa647plus donkey anti-rabbit (A32795, Invitrogen) |
| Fig. 4E | mELAST human brain | MegaSPIM 2x obj. | - | - | No staining |
| Fig. 4F | SHIELD human brain | Leica 20x obj. | 1XPBSTN | dPROTOS | Anti-CR (Rabbit, RPCA-Calr, Encor) + Alexta488 donkey anti-rabbit (ab150073, abcam) |
| fig. S13B | mELAST human brain | MegaSPIM 16.7x obj. | 0.2XPBSNaCh | dPROTOS | Anti-pNFP (Mouse IgG1 & IgM, 837904, Biolegend) + Setau647 goat anti-mouse (115-007-185, JacksonImmunoResearch Lab) |
| fig. S13C | mELAST human brain | Leica 20x obj. | i. 1XPBSTN<br>ii. 1XPBSTN | Water | Nuclei dye (POPO-1, Invitrogen)<br>Anti-PV (Rabbit, PA1-933, Invitrogen) |

| | | | | | |
|---|---|---|---|---|---|
| | | | iii. 0.2XPBSNaCh | | + RRX donkey anti-rabbit (711-295-152, JacksonImmunoResearch Lab) |
| | | | | | Anti-NFH (Mouse IgG1, MCA-NAP4, Encor)<br>+ Setau647 goat anti-mouse (custom) |
| Fig. 4I | mELAST human brain | MegaSPIM 2x and 16.7x objectives | 0.2XPBSNaCh | dPROTOS | Nuclei dye (YOYO-1, Invitrogen) |
| | | | | | Anti-CR (Rabbit, RPCA-Calr, Encor)<br>+ RRX donkey anti-rabbit (711-295-152, JacksonImmunoResearch Lab) |
| | | | | | Anti-SST (Rat, Sc-47706, Santa Cruz)<br>+ Alexa647plus donkey anti-rat (A48272, Invitrogen) |
| Fig. 4J<br>Fig. 5C<br>Fig. 5F, left<br>Fig. 6A, upper<br>Fig. 6H, left<br>Fig. 6J, left<br>fig. S15<br>fig. S17<br>fig. S20, left | mELAST human brain | MegaPIM 2x and 16.7x objectives | 0.2XPBSNaCh | dPROTOS | (For all rounds) Nuclei dye (YOYO-1, Y3601, Invitrogen) |
| | | | | | (R1) Before mELAST processing, Anti-NeuN (Rabbit, RPCA-FOX3, Encor)<br>+ RRX donkey anti-rabbit (711-295-152, JacksonImmunoResearch Lab) |
| | | | | | (R1) Before mELAST processing, Lectin-649 (DL-1177, Vector) |
| | | | | | (R2) Anti-Iba1 (Rabbit, Ab178847, Abcam)<br>+ RRX donkey anti-rabbit (711-295-152, JacksonImmunoResearch Lab) |
| | | | | | (R2) Anti-Aβ (Mouse IgG2b, MA5-36246, Invitrogen)<br>+ Alexa647plus donkey anti-mouse (A32787, Invitrogen) |
| | | | | | (R3) Anti-NPY (Rabbit, 77100, CST)<br>+ RRX donkey anti-rabbit (711-295-152, JacksonImmunoResearch Lab) |
| | | | | | (R3) Anti-SST (Rat, Sc-47706, Santa Cruz)<br>+ Alexa647plus donkey anti-rat (A48272, Invitrogen) |

| | | | | | |
|---|---|---|---|---|---|
| | | | | | (R4) Anti-GFAP (Mouse IgG2b κ, sc-33673, Invitrogen)<br>+ RRX donkey anti-mouse (103-297-008, JacksonImmunoResearch Lab) |
| | | | | | (R4) Anti-PV (Rabbit, PA1-933, Invitrogen)<br>+Alexa647plus donkey anti-mouse (A32795, Invitrogen) |
| | | | | | (R5) Anti-CR (Rabbit, RPCA-Calr, Encor)<br>+ RRX donkey anti-rabbit (711-295-152, JacksonImmunoResearch Lab) |
| | | | | | (R5) Anti-pTau (Mouse IgG1, MN1060, Invitrogen)<br>+ Alexa647plus donkey anti-mouse (A32787, Invitrogen) |
| | | | | | (R6) Anti-MBP (Rat, Ab7349, Abcam)<br>+ RRX donkey anti-rat (712-295-153, JacksonImmunoResearch Lab) |
| | | | | | (R6) Anti-pNFP (Mouse IgG1 & IgM, 837904, Biolegend)<br>+ Alexa647plus donkey anti-mouse (A32787, Invitrogen) |
| | | | | | (R7) Anti-GFAP (Mouse IgG2b κ, sc-33673, Invitrogen)<br>+ RRX donkey anti-mouse (103-297-008, JacksonImmunoResearch Lab) |
| | | | | | (R7) Anti-CD31 (Goat, AF3628, R&D systems)<br>+ Alexa647plus donkey anti-goat (A32787, Invitrogen) |
| | | | | | (R8) Anti-MBP (Rat, Ab7349, Abcam)<br>+ RRX donkey anti-rat (712-295-153, JacksonImmunoResearch Lab) |
| | | | | | (R8) Anti-pNFP (Mouse IgG1 & IgM, 837904, Biolegend) |

| | | | | | |
|---|---|---|---|---|---|
| | | | | | + Setau647 goat anti-mouse (115-007-185, JacksonImmunoResearch Lab) |
| Fig. 5A | SHIELD human brain | MegaSPIM 2x obj. | 1XPBSNaCh | dPROTOS | Nuclei dye (YOYO-1, Invitrogen) |
| | | | | | Anti-NeuN (Rabbit, RPCA-FOX3, Encor)<br>+ RRX donkey anti-rabbit (711-295-152, JacksonImmunoResearch Lab) |
| Fig. 5D<br>Fig. 5F, right<br>Fig. 6A, lower<br>Fig. 6H, right<br>Fig. 6J, right<br>fig. S18<br>fig. S20, right<br>fig. S21 | mELAST human brain | MegaSPIM 2x and 16.7x objectives | 0.2XPBSNaCh | dPROTOS | (For all rounds) Nuclei dye (YOYO-1, Y3601, Invitrogen) |
| | | | | | (R1) Before mELAST processing, Anti-NeuN (Rabbit, RPCA-FOX3, Encor)<br>+ RRX donkey anti-rabbit (711-295-152, JacksonImmunoResearch Lab) |
| | | | | | (R1) Before mELAST processing, Lectin-649 (DL-1177, Vector) |
| | | | | | (R2) Anti-Iba1 (Rabbit, Ab178847, Abcam)<br>+ RRX donkey anti-rabbit (711-295-152, JacksonImmunoResearch Lab) |
| | | | | | (R2) Anti-Aβ (Mouse IgG2b, MA5-36246, Invitrogen)<br>+ Alexa647plus donkey anti-mouse (A32787, Invitrogen) |
| | | | | | (R3) Anti-NPY (Rabbit, 77100, CST)<br>+ RRX donkey anti-rabbit (711-295-152, JacksonImmunoResearch Lab) |
| | | | | | (R3) Anti-SST (Rat, Sc-47706, Santa Cruz)<br>+ Alexa647plus donkey anti-rat (A48272, Invitrogen) |
| | | | | | (R4) Anti-GFAP (Mouse IgG2b κ, sc-33673, Invitrogen)<br>+ RRX donkey anti-mouse (103-297-008, JacksonImmunoResearch Lab) |
| | | | | | (R4) Anti-PV (Rabbit, PA1-933, Invitrogen) |

| | | | | | |
|---|---|---|---|---|---|
| | | | | | +Alexa647plus donkey anti-mouse (A32795, Invitrogen) |
| | | | | | (R5) Anti-pTau (Mouse IgG1, MN1060, Invitrogen)<br>+ RRX donkey anti-mouse (115-297-185, JacksonImmunoResearch Lab) |
| | | | | | (R5) Anti-CR (Rabbit, RPCA-Calr, Encor)<br>+Alexa647plus donkey anti-mouse (A32795, Invitrogen) |
| | | | | | (R6) Anti-CD31 (Goat, AF3628, R&D systems)<br>+ RRX donkey anti-goat (805-025-180, JacksonImmunoResearch Lab) |
| | | | | | (R6) Anti-Aβ (Mouse IgG2b, MA5-36246, Invitrogen)<br>+ Alexa647plus donkey anti-mouse (A32787, Invitrogen) |
| | | | | | (R7) Anti-MBP (Rat, Ab7349, Abcam)<br>+ RRX donkey anti-rat (712-295-153, JacksonImmunoResearch Lab) |
| | | | | | (R7) Anti-pNFP (Mouse IgG1 & IgM, 837904, Biolegend)<br>+ Setau647 goat anti-mouse (115-007-185, JacksonImmunoResearch Lab) |
| Fig. 6L | mELAST human brain | Leica 63x obj. | 1XPBSNaCh | Water | (R8) Anti-Aβ (Mouse IgG2b, MA5-36246, Invitrogen)<br>+ Alexa488plus donkey anti-mouse (A32766, Invitrogen) |
| | | | | | (R8) Anti-PSD95 (Guinea Pig, 124 014, SYSY)<br>+ Alexa568plus donkey anti-Guinea Pig (A11075, Invitrogen) |
| | | | | | (R8) Anti-Synapsin I/II (Rabbit, 106 102, SYSY)<br>+ Alexa647plus donkey anti-rabbit (A32795, Invitrogen) |

| | | | | | |
|---|---|---|---|---|---|
| Fig. 7B<br>fig, S26 | SHIELD mouse brain | MegaSPIM 4x obj. | 1XPBSNaCh | dPROTOS | Lectin-488 (DL-1174, Vector) |
| | | | | | Anti-V5 (Chicken, RPCA-FOX3, Encor)<br>+ RRX donkey anti-rabbit (711-295-152, JacksonImmunoResearch Lab) |
| fig. S25 | mELAST human brain | MegaSPIM 16.7x obj. | 0.2XPBSNaCh | dPROTOS | Lectin, Biotinylated (B-1175-1)<br>+ RRX Streptavidin (S6366, Invitrogen) |
| | | | | | Anti-NFH (Mouse IgG1, MCA-NAP4, Encor)<br>+ Setau647 goat anti-mouse (115-007-185, JacksonImmunoResearch Lab) |
| Fig. 7D | SHIELD human brain | MegaSPIM 2x obj. | 1XPBSNaCh | dPROTOS | Lectin-649 (DL-1177, Vector) |
| Fig. 7E<br>fig. S28 | SHIELD human pons | MegaSPIM 2x obj. | 1XPBSNaCh | dPROTOS | Lectin-649 (DL-1177, Vector) |
| Fig. 8A-B<br>fig. S29 | SHIELD human brain | Dfly 20x obj. | 1XPBSNaCh | dPROTOS | Lectin-488 (DL-1174, Vector) |
| | | | | | Anti-GFAP (Chicken, Aves)<br>+ RRX donkey anti-chicken (703-025-155, JacksonImmunoResearch Lab) |
| | | | | | Anti-NFH (Mouse IgG1, MCA-NAP4, Encor)<br>+Alexa680plus donkey anti-mouse (A32788, Invitrogen) |
| | | | | | Anti-PV (Rabbit, PA1-933, Invitrogen)<br>+Alexa800plus donkey anti-rabbit (A32808, Invitrogen) |
| Fig. 8D, F, G, and J<br>fig. S30 | mELAST human brain | MegaSPIM 16.7x obj. | 0.2XPBSNaCh | dPROTOS | Anti-Aβ (Rabbit, RPCA-Calr, Encor)<br>+ Setau488 goat anti-rabbit (111-007-008, JacksonImmunoResearch Lab) |
| | | | | | Lectin, Biotinylated (B-1175-1)<br>+ RRX Streptavidin (S6366, Invitrogen) |
| | | | | | Anti-pTau (Mouse IgG1, MN1060, Invitrogen)<br>+ Setau647 goat anti-mouse (115-007-185, JacksonImmunoResearch Lab) |

**Table S5.**

Number of YOYO-1 (nuclei) in image sub volume of the non-demented control and AD human brain tissues (figs. S14 and S16).

| Tissue | Cortical layer | | | | | |
|---|---|---|---|---|---|---|
| | I | II | III | IV | V | VI |
| Control (5.46 $mm^3$) | 714 | 1630 | 11720 | 1768 | 4579 | 4099 |
| AD (50.02 $mm^3$) | 613 | 8221 | 65664 | 7318 | 24502 | 33399 |

**Table S6.**
Number of neuron types in image sub volume of the non-demented control human brain tissue (5.46 $mm^3$) (figs. S14).

| Cell type (antigen) | | | | | Cortical layer | | | | | |
|---|---|---|---|---|---|---|---|---|---|---|
| PV | CR | NPY | SST | Iba1 | I | II | III | IV | V | VI |
| + | - | - | - | - | 3 | 6 | 419 | 86 | 328 | 188 |
| - | + | - | - | - | 88 | 109 | 396 | 33 | 30 | 17 |
| - | - | + | - | - | 2 | 0 | 7 | 2 | 5 | 6 |
| - | - | - | + | - | 0 | 0 | 2 | 0 | 2 | 0 |
| - | - | - | - | + | 22 | 47 | 469 | 61 | 227 | 268 |
| + | + | - | - | - | 38 | 51 | 295 | 17 | 18 | 6 |
| - | - | + | + | - | 0 | 0 | 2 | 0 | 2 | 0 |
| - | - | - | + | + | 0 | 0 | 0 | 0 | 0 | 0 |
| + | - | + | - | - | 0 | 0 | 0 | 0 | 0 | 0 |
| + | - | - | + | - | 0 | 0 | 0 | 0 | 0 | 0 |
| + | - | - | - | + | 0 | 0 | 0 | 0 | 0 | 0 |
| - | + | + | - | - | 0 | 0 | 0 | 0 | 0 | 0 |
| - | + | - | + | - | 0 | 0 | 0 | 0 | 0 | 0 |
| - | + | - | - | + | 0 | 0 | 0 | 0 | 0 | 0 |
| - | - | + | - | + | 0 | 0 | 0 | 0 | 0 | 0 |
| - | - | - | + | + | 0 | 0 | 0 | 0 | 0 | 0 |
| + | + | + | - | - | 0 | 0 | 0 | 0 | 0 | 0 |
| + | + | - | + | - | 0 | 0 | 0 | 0 | 0 | 0 |
| + | + | - | - | + | 0 | 0 | 0 | 0 | 0 | 0 |
| + | - | + | + | - | 0 | 0 | 0 | 0 | 0 | 0 |
| + | - | - | + | + | 0 | 0 | 0 | 0 | 0 | 0 |
| - | + | + | + | - | 0 | 0 | 0 | 0 | 0 | 0 |
| - | + | + | - | + | 0 | 0 | 0 | 0 | 0 | 0 |
| - | - | + | + | + | 0 | 0 | 0 | 0 | 0 | 0 |
| + | + | + | + | - | 0 | 0 | 0 | 0 | 0 | 0 |
| - | + | + | + | + | 0 | 0 | 0 | 0 | 0 | 0 |
| + | + | + | + | + | 0 | 0 | 0 | 0 | 0 | 0 |

**Table S7.**

Number of neuron types in image sub volume of the AD human brain tissue (50.02 $mm^3$) (figs. S16).

| Cell type (antigen) | | | | | Cortical layer | | | | | |
|---|---|---|---|---|---|---|---|---|---|---|
| PV | CR | NPY | SST | Iba1 | I | II | III | IV | V | VI |
| + | - | - | - | - | 10 | 190 | 1883 | 204 | 633 | 428 |
| - | + | - | - | - | 20 | 253 | 1612 | 108 | 323 | 328 |
| - | - | + | - | - | 0 | 9 | 80 | 3 | 20 | 37 |
| - | - | - | + | - | 0 | 0 | 4 | 4 | 21 | 12 |
| - | - | - | - | + | 55 | 964 | 6700 | 942 | 3894 | 5532 |
| + | + | - | - | - | 1 | 261 | 821 | 12 | 38 | 16 |
| - | - | + | + | - | 0 | 0 | 6 | 0 | 7 | 1 |
| - | - | - | + | + | 0 | 0 | 0 | 0 | 0 | 0 |
| + | - | + | - | - | 0 | 0 | 0 | 0 | 0 | 0 |
| + | - | - | + | - | 0 | 0 | 0 | 0 | 0 | 0 |
| + | - | - | - | + | 0 | 0 | 0 | 0 | 0 | 0 |
| - | + | + | - | - | 0 | 0 | 0 | 0 | 0 | 0 |
| - | + | - | + | - | 0 | 0 | 0 | 0 | 0 | 0 |
| - | + | - | - | + | 0 | 0 | 0 | 0 | 0 | 0 |
| - | - | + | - | + | 0 | 0 | 0 | 0 | 0 | 0 |
| - | - | - | + | + | 0 | 0 | 0 | 0 | 0 | 0 |
| + | + | + | - | - | 0 | 0 | 0 | 0 | 0 | 0 |
| + | + | - | + | - | 0 | 0 | 0 | 0 | 0 | 0 |
| + | + | - | - | + | 0 | 0 | 0 | 0 | 0 | 0 |
| + | - | + | + | - | 0 | 0 | 0 | 0 | 0 | 0 |
| + | - | - | + | + | 0 | 0 | 0 | 0 | 0 | 0 |
| - | + | + | + | - | 0 | 0 | 0 | 0 | 0 | 0 |
| - | + | + | - | + | 0 | 0 | 0 | 0 | 0 | 0 |
| - | - | + | + | + | 0 | 0 | 0 | 0 | 0 | 0 |
| + | + | + | + | - | 0 | 0 | 0 | 0 | 0 | 0 |
| - | + | + | + | + | 0 | 0 | 0 | 0 | 0 | 0 |
| + | + | + | + | + | 0 | 0 | 0 | 0 | 0 | 0 |

**Table S8.**

Number of NFT (neurofibril tangles) types in sub image volume of the AD human brain tissue (5.83 $mm^3$) (fig. S16).

| NFT type (antigen) | | Cortical layer | | | | | |
|---|---|---|---|---|---|---|---|
| pNFP | pTau | I | II | III | IV | V | VI |
| **-** | **+** | 1 | 3 | 14 | 4 | 7 | 4 |
| **+** | **-** | 2 | 7 | 27 | 1 | 14 | 3 |
| **+** | **+** | 1 | 6 | 48 | 2 | 32 | 14 |
| Total | | 4 | 16 | 89 | 7 | 53 | 21 |

### Captions for Movies S1 to S11

#### Movie S1.

MEGAtome operation for slicing intact human brain hemisphere and mouse brain arrays.

#### Movie S2.

Navigation of the image volume in Fig. 4I (i-iii) showing YOYO-1 nuclei, CR+, SST+ cells in the mELAST-processed human brain slab (non-demented control) at cellular level.

#### Movie S3.

Navigation of the image volume in Fig. 4I (iv) showing CR+, SST+ cells in the mELAST-processed human brain slab (non-demented control) at subcellular level.

#### Movie S4.

Navigation of the image volume in Fig. 4J, Fig. 5C, and figs. S17 and S18 showing the multiplexed mELAST-processed human brain tissues of the non-demented control case.

#### Movie S5.

Navigation of the image volume in Fig. 5D and fig. S16 showing the multiplexed mELAST-processed human brain tissues of the AD case.

#### Movie S6.

Navigation of the image volume in Fig. 5C-D showing spatial distribution of pTau, Aβ, and GFAP in the mELAST-processed human brain tissues (the control and AD cases).

#### Movie S7.

Navigation of the image volume in Fig. 6A showing subcellular morphological details of the cells in the mELAST-processed human brain tissues (the control and AD cases).

#### Movie S8.

Navigation of the image volume in Fig. 6L showing nanoscopic synapse in the mELAST-processed human brain tissues (the control and AD cases).

#### Movie S9.

Navigation of the image volume in Fig. 7B and fig. S25 showing the UNSLICE'd MORF mouse brain hemisphere and mELAST human brain tissues.

#### Movie S10.

Navigation of the image volume in Fig. 7D and fig. S26 showing the UNSLICE'd SHIELD human brain slabs.

#### Movie S11.

Navigation of the image volume in Fig. 8, A and B showing multi-channel axon-level projectome mapping in UNSLICE'd SHIELD-processed human brain slabs.

**Movie S12.**

Navigation of the image volume in Fig. 8, F and G showing the pTau+ fibers tracing from UNSLICE'd mELAST-processed AD human brain tissues.
